# Rational design of T-DNA vectors enables predictable, single-copy integration in *Arabidopsis thaliana*

**DOI:** 10.64898/2026.06.08.730999

**Authors:** William M. Shaw, Anjali Gajendiran, Elizabeth I. Tchantouridze, Lindsey L. Bechen, Sean G. Clarke, Sarah Guiziou, Mary Gehring, Ahmad S. Khalil

## Abstract

*Agrobacterium*-mediated transformation is the dominant method for plant transgenesis, yet it frequently produces multi-copy, structurally complex T-DNA insertions associated with transgene silencing, unpredictable expression, and genome instability. Here, leveraging a high-throughput phenotypic reporter, we systematically dissect how T-DNA vector architecture, plasmid biology, and regulatory element choice shape transformation outcomes in *Arabidopsis thaliana*. We discover a pronounced trade-off between transformation efficiency and T-DNA copy number, uncovering the virulence enhancing overdrive sequence as a major determinant of this relationship. Guided by these insights, we engineered a new T-DNA vector that balances efficient transformation with predominantly single-copy integration. Additionally, we replaced viral elements, such as the widely used CaMV 35S promoter, with Arabidopsis-derived regulatory elements to minimise undesired enhancer effects, and developed a streamlined workflow for efficient T-DNA insertion mapping in the genome. Together, these advances form the T1 vector series, an Arabidopsis-optimised T-DNA vector system that enables clean, single-copy, and readily mappable transgene integration with predictable expression in the first generation after transformation.

## INTRODUCTION

*Agrobacterium*-mediated transformation (AMT) is the preferred method for generating transgenic plants due to its ease of use and capacity to deliver large genetic payloads^1–3^. The method harnesses the natural gene transfer machinery of *Agrobacterium tumefaciens* and related species, replacing its plasmid-encoded tumour-inducing genes with user-defined sequences to disarm the bacterium while enabling random integration of transgenes across diverse plant species^1, 4^. Subsequent technical refinements decoupled the transfer DNA (T-DNA) region from the virulence (*vir*) genes required for T-DNA processing and delivery, giving rise to a modular binary vector system in which virulence functions and transgene cargo are maintained on separate plasmids^5, 6^. DNA positioned between the left and right border (LB and RB) sequences of the T-DNA binary vector is efficiently transferred to the plant genome, allowing new sequences to be readily cloned between these sites and establishing AMT as a versatile and indispensable tool for plant research and biotechnology^1, 5^.

Despite these advances, transformation remains a major bottleneck in plant research (**Fig. 1a**). Transformation efficiencies are often low and highly variable across species and even among different ecotypes. Moreover, successful events frequently involve multi-copy or structurally complex insertions that are prone to epigenetic transcriptional silencing, resulting in unstable or unpredictable transgene expression and, in some cases, genomic instability^7–9^. The inherently random nature of T-DNA integration further exacerbates expression variability due to positional effects^10, 11^. Consequently, researchers typically generate large numbers of independent transformants and perform extensive downstream screening to identify lines with single-copy insertions and appropriate expression profiles. In tractable species this process is at best a laborious and time-consuming inconvenience, while in recalcitrant plants it can become a critical rate-limiting step^1^. These constraints are particularly restrictive for plant synthetic biology, where rapid iteration through the design–build–test–learn cycle is essential^12^.

**Fig. 1:**
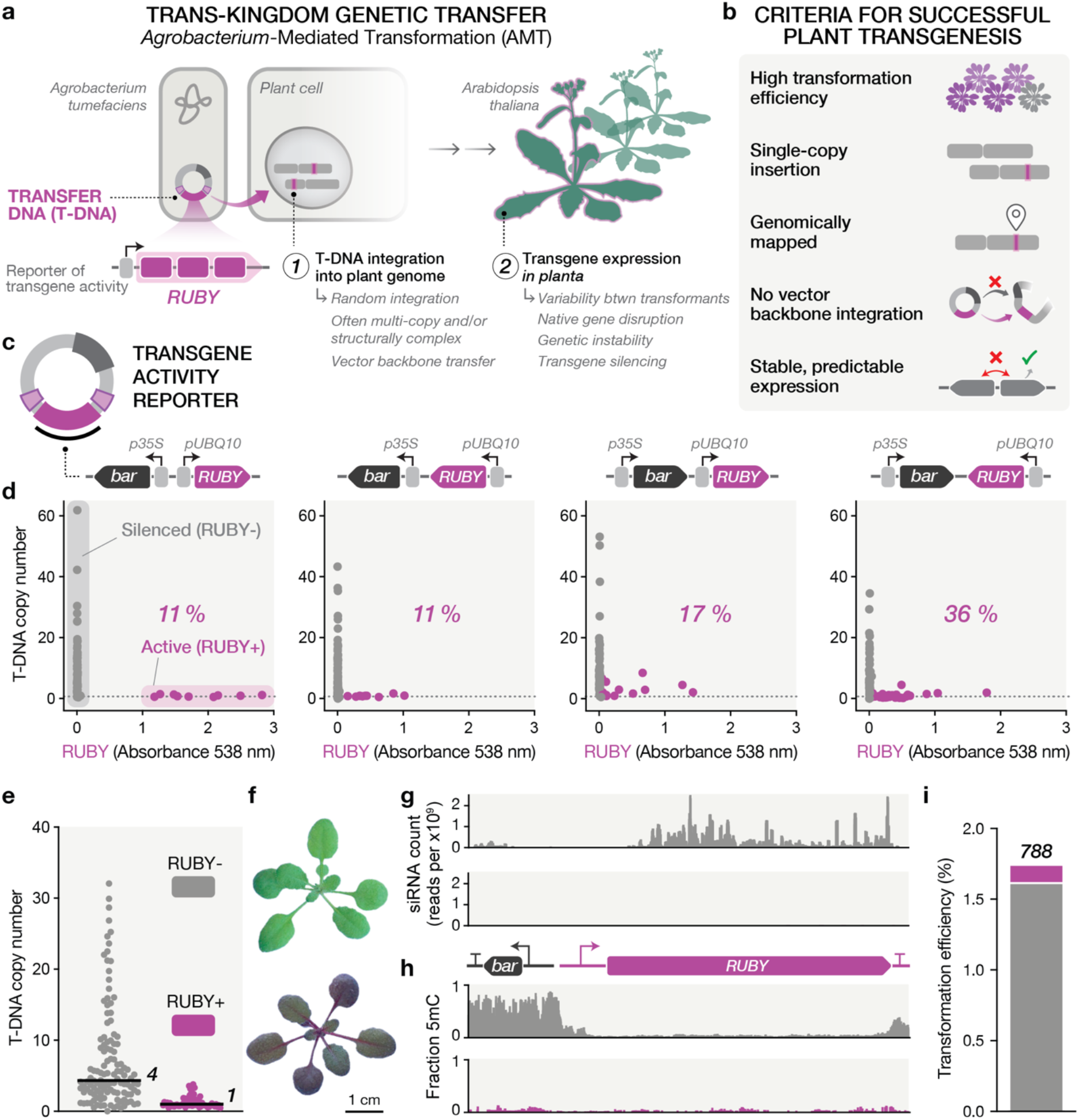
RUBY transgene activity serves as a visual reporter of low-copy T-DNA integration in *A. thaliana*. **a**, Schematic overview of *Agrobacterium*-mediated transformation and current associated challenges. **b**, Criteria for successful plant transgenesis. **c**, Organisation of the BASTA resistance (*bar*) and RUBY expression cassettes within the T-DNA across four construct arrangements. **d**, Relationship between T-DNA copy number (measured by qPCR) and relative betalain absorbance at 538 nm for 90 randomly selected transformants per construct. Pink (RUBY+) and grey (RUBY-) points indicate samples above and below an absorbance threshold of 0.05, respectively. The percentage of RUBY+ plants is shown within each panel. qPCR copy number measurements were normalised to validated single- and double-copy reference lines. All data points are shown except one out-of-range sample in the third condition (T-DNA copy number = 134; absorbance at 538 nm = 0.002) In a small number of cases, qPCR failed to detect RUBY sequences, consistent with T-DNA truncation events. **e**, T-DNA copy number distributions for 120 RUBY- and 64 RUBY+ plants derived from the divergent-gene T-DNA arrangement (**d**, left-most construct), as determined by qPCR. Black horizontal lines denote median values. Numerical labels indicate the corresponding median, rounded to one significant figure. **f**, Representative transformants illustrating the binary nature of RUBY expression in green (RUBY-) and red (RUBY+) plants at 21 days post-germination (DPG). **g**, Small RNA profiles (21-24-nt) mapped across the T-DNA from representative RUBY-(top, grey) and RUBY+ (bottom, pink) plants. **h**, Whole-genome EM-seq analysis showing 5-methylcytosine (5mC) profiles across the T-DNA from representative green (top, grey) and red (bottom, pink) plants in CG, CHG, and CHH sequence contexts. **i**, Transformation efficiency of the divergent-gene T-DNA construct, showing 58 RUBY+ (pink) and 730 RUBY-(grey) plants among all transformants obtained from 45,000 seeds (788 seedlings total).

To accelerate plant research, an effective transformation system should satisfy five key criteria: (i) high transformation efficiency; (ii) single-copy T-DNA insertion; (iii) straightforward mapping of integration sites; (iv) backbone-free, or “clean”, integration of the intended T-DNA sequence; and (v) predictable and stable transgene expression (**Fig. 1b**).

To date, efforts to improve AMT have focused almost exclusively on maximising transformation efficiency, through optimisation of virulence induction conditions^13^, *Agrobacterium* strain selection^14^, or bacteria–plant co-cultivation parameters^15^. Other approaches have targeted the virulence machinery more directly, for example by modifying or overexpressing *vir* proteins or host-derived factors, often using ternary vector systems to supply additional functions^16, 17^. More recently, complete refactoring of the virulence plasmid and engineering of binary vector copy number have emerged as promising strategies^18, 19^. However, few studies have systematically examined how these efficiency-focused interventions influence downstream transformation outcomes, such as transgene copy number, insertion complexity, and epigenetic silencing, which ultimately determine the utility and reliability of the resulting transgenic lines.

Here, we set out to rationally design a new T-DNA vector system for *Arabidopsis thaliana* that addresses the key criteria of successful plant transgenesis. Central to this effort was the use of the RUBY reporter to distinguish predominantly single-copy T-DNA integration events. Using this assay, we systematically dissected T-DNA border architecture and uncovered a central role for the overdrive sequence, a cis-acting virulence enhancer, in shaping integration outcomes. This revealed a clear trade-off between transformation efficiency, T-DNA copy number, and vector backbone integration. Guided by these insights, we engineered a series of vectors with tuneable transfer rates and evaluated their performance across diverse experimental conditions. In parallel, we addressed the pervasive bidirectional enhancer activity of the widely used CaMV 35S promoter by identifying alternative Arabidopsis regulatory elements that support robust selection without perturbing neighbouring transgene expression. Finally, we established a streamlined and cost-effective workflow for efficient mapping of T-DNA insertion sites in the plant genome. By integrating these advances, we present an Arabidopsis-optimised T-DNA vector system that balances transformation efficiency with backbone-free, predictable, single-copy integration.

## RESULTS

### RUBY expression reports predominantly single-copy T-DNA integration

Assessing plant transformation outcomes beyond simple efficiency metrics remains a major challenge (**Fig. 1a**). Traditional approaches to evaluate transgene expression rely on low-throughput measurement of mRNA levels or reporter gene activity^20^, while determining copy number and integration complexity requires laborious or costly techniques such as quantitative PCR or whole-genome sequencing^21^. The development of the RUBY reporter represents a significant advance by enabling direct visual detection of transgene expression in living plants^22^. RUBY encodes a polycistronic, three-gene biosynthetic pathway that converts tyrosine into the red pigment betalain, producing visible tissue colouration that allows rapid assessment of expression levels. Notably, RUBY expression is frequently silenced in a substantial fraction of transformants, particularly under conditions of high constitutive expression^23, 24^. Recent work has linked this phenomenon to specific sequence features that trigger siRNA production, with higher T-DNA copy numbers observed in silenced lines^25^. This property provides an opportunity to exploit RUBY as a scalable, visual indicator of additional transformation outcomes, including transgene copy number and stability (**Fig. 1b**).

To investigate the basis of this behaviour and determine whether it could be leveraged as a visual readout of transformation outcomes beyond simple efficiency metrics, we analysed RUBY expression across four distinct genetic arrangements (**Fig. 1c**). This experimental design allowed us to systematically assess the influence of T-DNA gene organisation and selection marker expression on transgene expression in Arabidopsis.

Our parental T-DNA vector was derived from the Plant MoClo toolkit plasmid pAGM4723 (Addgene ID: 48015)^26^, allowing us to leverage the Golden Gate cloning framework while providing an adaptable backbone without a predefined selection marker. This vector incorporates a pVS1 replicon and border sequences from *Agrobacterium tumefaciens* C58 nopaline-type Ti plasmid pTiC58, closely resembling the plasmid architecture of the widely adopted pCAMBIA series^27^. In all constructs, the RUBY coding sequence was positioned adjacent to the right border (RB) and was under the control of the strong constitutive Arabidopsis *UBQ10* promoter and *HSP18*.*2 (HSP)* terminator, while a BASTA resistance marker (*bar*) under control of the CaMV 35S promoter and terminator was placed at the left border (LB) in all possible arrangements.

*Arabidopsis thaliana* Col-0 plants were transformed via the floral dip method^15, 28^, using *Agrobacterium tumefaciens* GV3101. Approximately 10,000 seeds were subsequently sown on soil and selected by BASTA herbicide application over two weeks. This large-scale screen revealed distinct expression patterns across constructs, with the majority of transformants exhibiting no visible betalain pigmentation (**Extended Data Fig. 1**).

We next sought to relate RUBY expression to T-DNA copy number using quantitative PCR (qPCR). Genomic DNA (gDNA) was extracted from 90 randomly selected individuals from all four construct architectures, and RUBY pigmentation was quantified by measuring absorbance in the gDNA extraction supernatant (**Extended Data Fig. 2**). This approach enabled a direct comparison between T-DNA copy number and reporter expression (**Fig. 1d**). Distinct expression profiles were observed for each expression cassette arrangement, with only 11–36% of plants exhibiting detectable RUBY absorbance above 0.05. Surprisingly, converging expression led to the least silencing (**Fig. 1d**, right-most construct), arguing against overlapping transcription and double-stranded RNA production as a dominant contributor to transgene silencing.

**Fig. 2:**
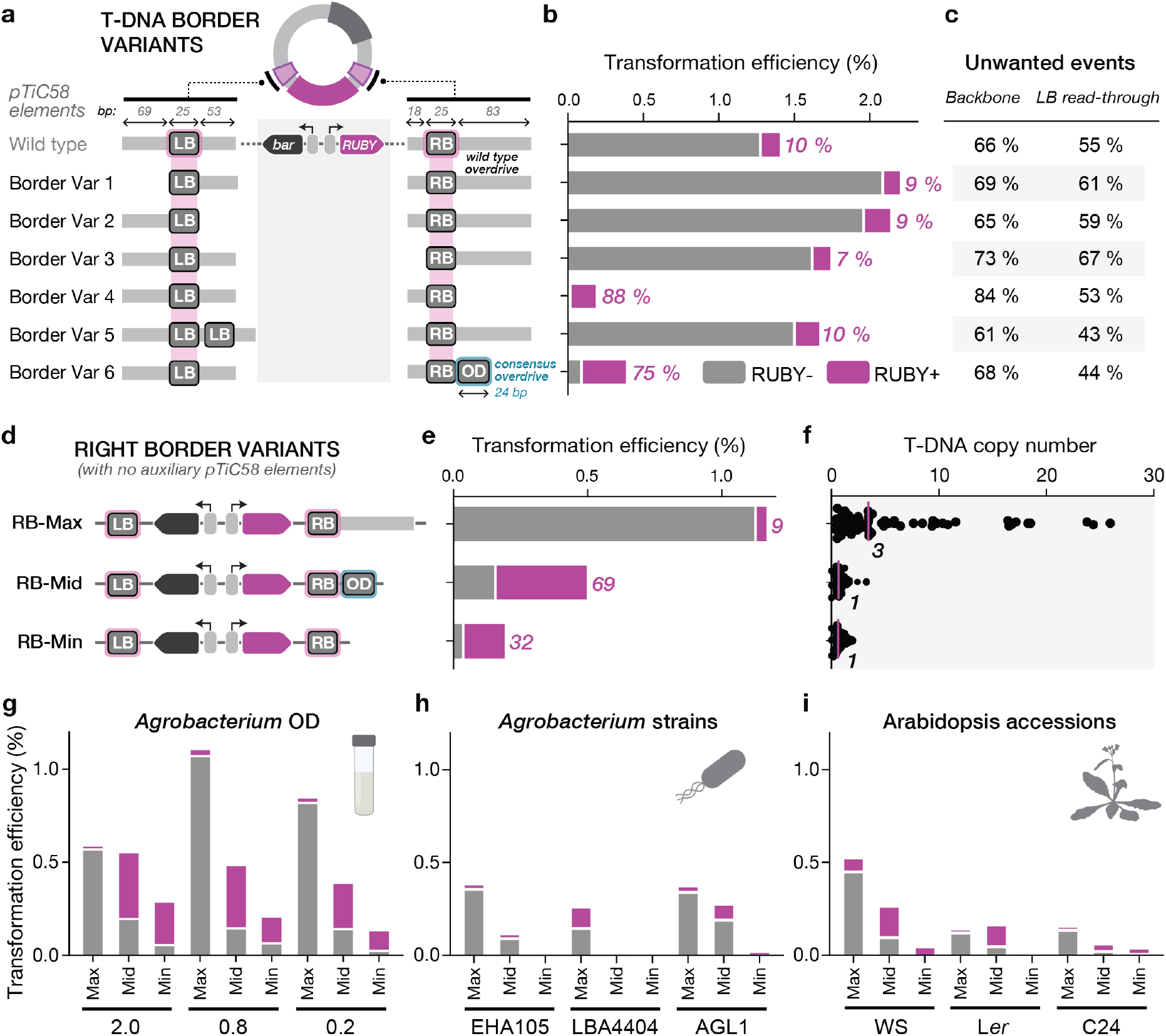
Systematic screening of *Agrobacterium* pTiC58 border sequence variants reveals changes to integration behaviour. **a**, Schematic overview of T-DNA border sequence variants. Grey regions indicate auxiliary sequences from pTiC58. The blue box (OD) denotes the 24-bp synthetic overdrive sequence. **b**, Transformation efficiency of T-DNA vectors carrying the indicated border variants, showing the proportion of RUBY-(grey) and RUBY+ (pink) plants. Percentage of RUBY+ plants shown in pink. Variable numbers of seeds were screened to obtain at least 100 independent transformants per condition, with a minimum of 20,000 seeds analysed for each. **c**, Frequency of vector backbone integration attributed to LB read-through and total backbone integration, assessed by low-cycle PCR on tissue from 96 randomly selected transformants per condition. **d**, Design of RB overdrive variants with all auxiliary pTiC58 sequences removed, retaining either the native pTiC58 overdrive (pRB-Max), the minimal synthetic overdrive (pRB-Mid), or no overdrive sequence (pRB-Min). **e**, Transformation efficiency of pRB-Max, pRB-Mid, and pRB-Min vectors, showing the proportion of RUBY- and RUBY+ plants, assessed from 20,000 seeds per construct. Total number of red plants shown in pink. **f**, T-DNA copy number distributions for pRB-Max, pRB-Mid, and pRB-Min, determined by qPCR from 62 randomly selected transformants per vector. Pink horizontal lines denote median values. Numerical labels indicate the corresponding median, rounded to one significant figure. Copy number measurements were normalised to validated single- and double-copy reference lines. **g–i**, Transformation efficiency of pRB-Max, pRB-Mid, and pRB-Min across varying experimental conditions: different *Agrobacterium* optical densities (OD) measured at 600 nM (**g**), different *Agrobacterium* strains (**h**), and different Arabidopsis accessions (**i**), with 20,000 seeds analysed per condition. Unless otherwise stated, experiments were performed using *Arabidopsis thaliana* Col-0 and *Agrobacterium tumefaciens* GV3101 at OD_600_ = 0.8.

Strikingly, across all arrangements, RUBY expression showed a strong inverse relationship with T-DNA copy number, with visible pigmentation largely restricted to low-copy insertions. This L-shaped distribution was most pronounced for the divergent gene orientation (**Fig. 1d**, left-most construct), in which RUBY expression was observed almost exclusively in plants carrying a single T-DNA insertion. Reporter output was highly binary, with individuals showing either robust pigmentation (RUBY+) or no visible colouration (RUBY-). The strong RUBY expression observed in this arrangement is likely driven by the combined effects of the 35S promoter’s potent enhancer activity and the favourable transcriptional environment created by divergent gene orientation^29, 30^. Notably, some plants containing a single T-DNA lacked red pigmentation, suggesting that RUBY expression in this context lies near a silencing threshold of approximately one T-DNA.

The observed relationship between integration and expression provides a potentially powerful visual marker for screening predominantly single-copy T-DNA insertion events. We therefore focused on the divergent gene orientation to refine this relationship and define features associated with single-copy integration (**Fig. 1e,f**). Additional transformants were grown, and 64 red and 120 green BASTA-resistant plants were randomly chosen based on their clear binary pigmentation phenotypes (**Extended Data Fig. 3**). Relative T-DNA copy number analysis confirmed our initial observations: RUBY expression was restricted to plants with low-copy integrations, with 67% of red (RUBY+) plants carrying a single T-DNA insertion and the remainder harbouring two (25%), three (5%), or four (3%) copies. Consistent with this trend, late-onset silencing was observed in all plants carrying more than two copies (**Extended Data Fig. 4**). In contrast, green (RUBY-) plants exhibited much greater copy number variation and a higher median copy number (4 T-DNAs), with a maximum of 32 T-DNAs observed.

To confirm whether RUBY-plants resulted from transgene silencing, we selected six green plants and two red plants for small RNA sequencing (**Fig. 1g and Extended Data Fig. 5**). This analysis revealed accumulation of 21-24-nt RNAs mapping across the RUBY coding sequence in green plants. Whole-genome methylation sequencing of the same plants revealed high methylation levels over the 35S promoter but low levels across the RUBY coding sequence (**Fig. 1h and Extended Data Fig. 6**). The presence of siRNAs and absence of DNA methylation suggest that in green plants RUBY silencing occurred at the post-transcriptional level, with transcriptional gene silencing not yet established in the T1 generation after three weeks. High rates of silencing appeared to be an intrinsic property of the RUBY coding sequence, consistent with recent findings^25^ (**Extended Data Fig. 7**). Alterations to codon usage, incorporation of twelve introns, and changes in promoter and terminator identity had no major effect on RUBY silencing rates.

Taken together, these results demonstrate that RUBY, particularly when expressed strongly in the divergent gene orientation, serves as a highly effective and scalable reporter for predominantly single-copy T-DNA insertions. Simply scoring plants as red or green provides an easily observable metric that complements traditional measures of transformation efficiency (**Fig. 1i and Extended Data Table 1**).

### Transformation efficiency, T-DNA copy number, and backbone transfer are largely determined by the overdrive sequence

Having established RUBY as a faithful reporter for large-scale screening of T-DNA copy number, we next investigated how modifications to the T-DNA vector might influence transformation outcomes in Arabidopsis. Since the only sequences in our vectors that are derived from the *Agrobacterium* pTiC58 virulence plasmid are the border sequences, we first explored the effects of changes to these regions. The left (LB) and right (RB) borders are 25-bp imperfect, direct repeats recognised by the virulence protein complex VirD1/VirD2, which binds the RB, nicking and covalently attaching to the 5′ end of the single-stranded T-DNA. DNA polymerase then repairs the nicked DNA by synthesising a new strand across the T-DNA region, displacing the VirD2-bound intermediate in the process. A second nick at the LB releases the single-stranded T-DNA (T-strand) for transfer into plant cells^1, 31^.

While the roles of the left and right border sequences are well established and essential for T-DNA transfer, their influence on downstream transformation quality, as defined by our criteria for successful plant transgenesis, has not been systematically examined in a comprehensive manner. In addition, these border elements are typically embedded within larger stretches of pTiC58-derived sequence in binary vectors, whose roles are poorly defined and highly context dependent^32^. A notable exception is the sequence immediately upstream of the RB, termed the overdrive sequence, a cis-acting virulence enhancer that facilitates VirD2 recruitment to the RB, thereby promoting efficient T-DNA processing and transfer.^33, 34^. To assess the contribution of auxiliary regions to transformation outcomes, we systematically truncated sequences internal and external to the border elements in our RUBY reporter T-DNA constructs (**Fig. 2a**, Border Var 1-4). We also evaluated a design containing tandem left borders (Border Var 5), which has been reported to reduce transfer of vector backbone sequences^35, 36^, as well as constructs in which the native 83-bp pTiC58 overdrive region was replaced with a minimal 24-bp overdrive sequence^33, 36^ (Border Var 6).

Our transformation assay revealed no major differences in integration behaviour between the original T-DNA vector and constructs missing the auxiliary sequences either side of the LB or internal to the RB (**Fig. 2b**, Border Var 1-3). In contrast, substantial effects were observed when altering sequences external to the RB, where the overdrive element resides. Removal of the native pTiC58 sequence (Border Var 4), or its replacement with a minimal overdrive motif (Border Var 6), led to dramatic reductions in transformation efficiency, underscoring the importance of this region. Notably, these modifications also shifted the ratio of red to green plants, indicating marked enrichment for low-copy T-DNA insertions at the expense of overall transformation efficiency. This effect was most pronounced in the complete absence of an overdrive sequence, where nearly all recovered transformants were red (88 %), consistent with predominantly single-copy integration events. Remarkably, despite a 6.7-fold lower transformation efficiency (Border Var 4) relative to the original vector (Wild type), the absolute number of red, low-copy transformants per 10,000 seed (18) exceeded that obtained with the standard T-DNA design (14), highlighting the practical appeal of this trade-off for applications prioritising low-copy insertions.

To determine whether modifications to the border regions affected proper T-DNA processing, we performed large-scale genotyping to detect transfer of vector backbone sequences into the plant genome using an optimised, low-cycle PCR assay (**Fig. 2c and Extended Data Fig.8–11**). This analysis revealed a high frequency of backbone integration across all conditions, consistent with previous reports^37, 38^. Surprisingly, removal of auxiliary sequences flanking the LB and inside of the RB had little effect on backbone transfer (Border Var 1-3), in contrast to earlier studies^39^. In most cases, backbone transfer could be attributed to read-through at the LB, whereby T-DNA transfer initiates at the RB but fails to terminate at the LB, which was reduced with tandem LB sequences, as previously reported (Border Var 5)^35, 36^. Notably, the highest levels of backbone integration were observed under conditions with the lowest transformation efficiencies, when the overdrive sequence was absent (Border Var 4). Although readthrough frequencies were comparable to those of the original design (Wild type), the overall incidence of backbone integration was increased, suggesting increased initiation of T-DNA transfer from the LB. Reintroduction of a minimal overdrive sequence restored backbone integration frequencies to baseline levels (Border Var 6).

To further dissect the contribution of sequences outside the RB, we generated a panel of T-DNA vectors in which all auxiliary border-associated sequences were removed, while omitting or selectively retaining either the minimal or pTiC58 overdrive (**Fig. 2d**). These constructs, which yielded predominantly low, intermediate, or high transformation efficiencies, were designated pRB-Min, pRB-Mid, and pRB-Max, respectively. The resulting transformation patterns closely mirrored those observed in our earlier experiments, demonstrating that transformation efficiency and T-DNA copy number are largely dictated by the presence and composition of the overdrive sequence (**Fig. 2e**). To independently validate the red/green visual assay, we performed quantitative copy number analysis on 62 randomly selected transformants per vector (**Fig. 2f**). As expected, pRB-Max produced higher-copy integrations, with a median copy number of 3 and events approaching 30 copies, whereas both pRB-Min and pRB-Mid yielded predominantly low-copy insertions, with a maximum of 2 or 3, respectively, and a median copy number of 1 for both.

Finally, to evaluate the robustness of these three vectors across a range of experimental conditions, we transformed each construct using varying densities of *Agrobacterium tumefaciens*, different *Agrobacterium* strains, and several *Arabidopsis thaliana* accessions (**Fig. 2g–i**). Across a wide range of bacterial densities, the relative performance of the vectors was remarkably consistent. However, increasing *Agrobacterium* density imposed a clear penalty for pRB-Max, likely caused by excessive T-DNA delivery triggering an immune response^40^ (**Fig. 2g**). In contrast, the choice of *Agrobacterium* strain had a more pronounced impact, with all alternative strains resulting in reduced transformation efficiency while preserving the overall trends among vectors (**Fig. 2h**). Lastly, substantial variation was observed among different Arabidopsis accessions, consistent with previous reports^14^, yet each accession displayed a similar ranking of vector performance, underscoring the generality of these transformation properties (**Fig. 2i**).

These results identify pRB-Mid as providing the optimal balance between transformation efficiency (0.51%) and predominantly single-copy T-DNA integration (median copy number of 1), while avoiding the elevated vector backbone integration observed in the absence of an overdrive sequence (42% backbone) (**Extended Data Fig. 12**). In parallel, our analyses underscore the importance of *Agrobacterium* strain choice, establishing GV3101 as the most effective strain for transformation in Arabidopsis using this vector and the floral dip method, at least in the Col-0 background. Based on these findings, all subsequent vector development was performed using the pRB-Mid backbone in combination with GV3101 at a standard density of OD_600_ = 0.8 and the Col-0 accession.

### Higher T-DNA vector copy number in *Agrobacterium* increases integration copy number in Arabidopsis

A recent study demonstrated that transformation efficiency can be enhanced by introducing mutations into the T-DNA vector backbone that increase plasmid copy number in *Agrobacterium*^18^ (**Fig. 3a**). To assess how increased plasmid copy number influences transformation outcomes in our system, we introduced the high-copy RepA R106H mutation in the pVS1 replicon into the pRB-Mid vector. For direct comparison with the original study, we used the reported pCAMBIA-based vectors carrying kanamycin resistance and constitutive GFP reporter cassettes (*p35S:nptII:t35S* and *pCL2:EGFP:tUBQ3*). We cloned the same T-DNA construct into pRB-Mid to create a panel of equivalent vectors with and without the R106H mutation (**Fig. 3b**). We first confirmed the effect of this mutation in *Agrobacterium*, observing an approximately twofold increase in plasmid copy number for both vectors (**Fig. 3c**). Consistent with this, transient expression assays in *Nicotiana benthamiana* revealed a corresponding increase in GFP fluorescence, validating the functional impact of elevated plasmid copy number (**Fig. 3d**).

**Fig. 3:**
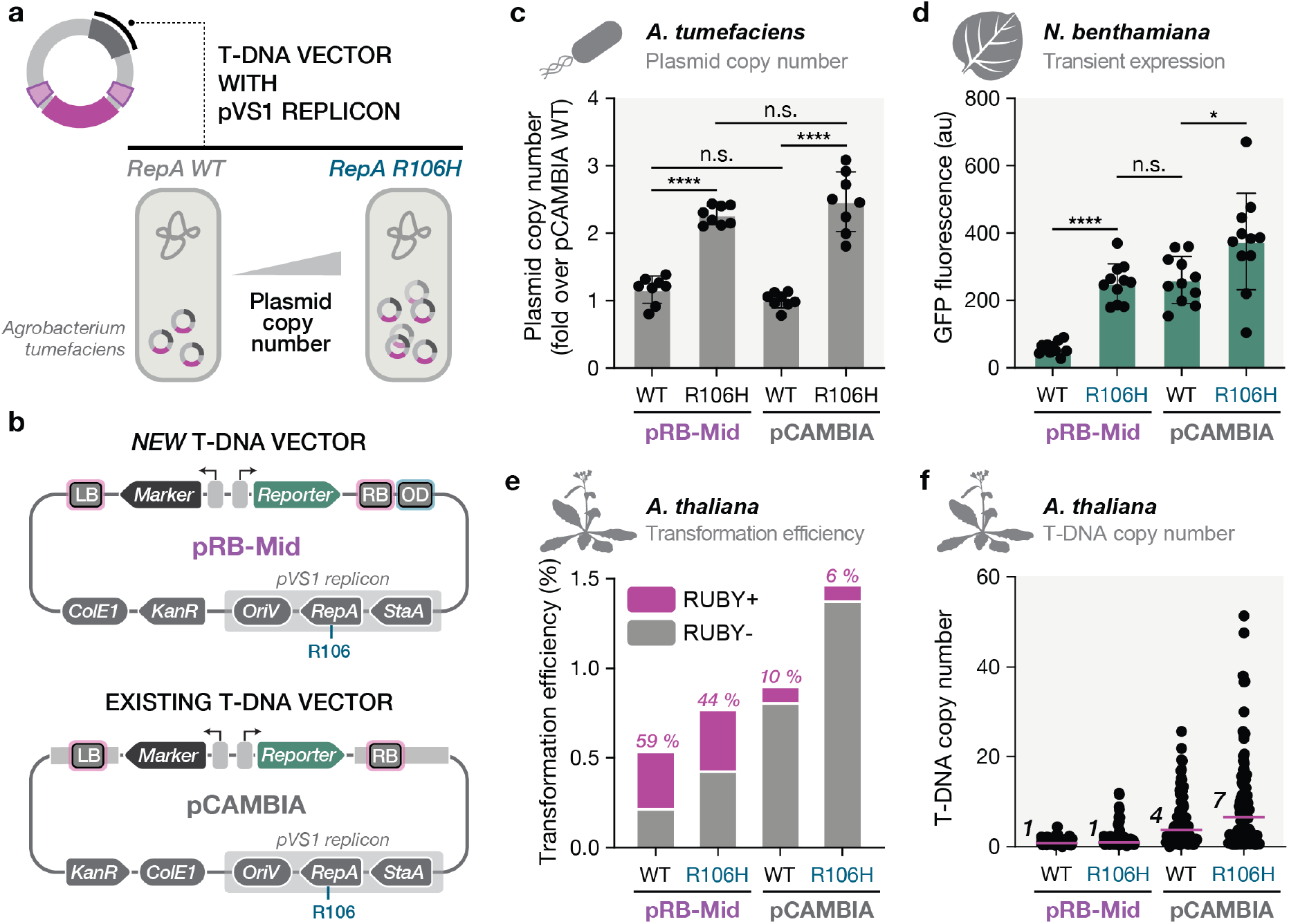
Increased plasmid copy number elevates T-DNA integration copy number in Arabidopsis floral dip transformation. **a**, Schematic overview of increased T-DNA plasmid copy number in *Agrobacterium tumefaciens* conferred by the R106H mutation in the pVS1 RepA replication protein. **b**, Plasmid architectures of pRB-Mid and pCAMBIA vectors, highlighting the RepA R106 residue targeted for mutation. **c**, Relative plasmid copy number of four T-DNA vector variants, pRB-Mid and pCAMBIA with (R106H) or without (WT) the high-copy mutation, as determined by qPCR in *A. tumefaciens* GV3101. Data were normalised to pCAMBIA (WT) and represent mean ± SD from eight biological replicates. **d**, GFP fluorescence from *N. benthamiana* leaf infiltrations using the four vector variants expressing GFP under the control of the low-strength constitutive promoter *pCL2*. Data represent mean ± SD from 11 independent infiltrations (averaging two leaf discs per infiltration) measured by plate reader at three days post-infiltration (DPI). Statistical significance in c,d was assessed by two-way ANOVA with multiple comparisons (GraphPad Prism). Significance levels are indicated as *p* < 0.05 (*), *p* < 0.0001 (****), and not significant (n.s.). **e**, Transformation efficiency of the four vector variants in Arabidopsis, showing the proportion of RUBY+ and RUBY-plants. For these experiments, T-DNA was replaced with *p35S:bar:t35S* and *pUBQ10:RUBY:tHSP*. Data represent 20,000 seeds analysed per construct. Percentage of RUBY+ plants shown in pink. **f**, T-DNA copy number distributions for the four vector variants in Arabidopsis, determined by qPCR from 96 randomly selected transformants per construct. Pink horizontal lines denote median values. Numerical labels indicate the corresponding median, rounded to one significant figure. Copy number measurements were normalised to validated single- and double-copy reference lines.

Introduction of the R106H mutation increased overall transformation efficiency in Arabidopsis for both pRB-Mid and pCAMBIA (**Fig. 3e**). However, this gain was accompanied by a higher proportion of green plants (94% vs 90% for pCAMBIA), consistent with increased T-DNA copy number and enhanced transgene silencing. Notably, boosting plasmid copy number raised the transformation efficiency of pRB-Mid to levels approaching that of the unmodified pCAMBIA vector, while maintaining a high fraction (44%) of RUBY+ plants. To directly test whether this effect reflected increased T-DNA integration, we isolated genomic DNA from 96 randomly selected transformants per condition and performed quantitative PCR-based copy number analysis (**Fig. 3f**). As expected, the R106H mutation resulted in higher T-DNA copy numbers in Arabidopsis. In the pCAMBIA-R106H background, the median copy number (7) was nearly doubled that of the unmodified vector (4). In contrast, although the maximum copy number increased for pRB-Mid-R106H, the median copy number (1) remained unchanged, indicating that other features of pRB-Mid partially buffer against the elevated integration associated with increased plasmid abundance.

### Screening Arabidopsis regulatory elements to avoid 35S promoter enhancer effects

Our initial experiments revealed that the CaMV 35S promoter (*p35S*) used to drive the selection marker can inadvertently increase expression of neighbouring ORFs (**Fig. 1d**), likely due to its well-characterised enhancer activity^30^. Moreover, this promoter is highly susceptible to silencing^41, 42^, as reflected by the accumulation of DNA methylation over the selection marker (**Fig. 1h**). Although such effects may be tolerable for constitutively expressed transgenes, they are problematic for conditional regulation intended for activity only in specific cell types or under particular conditions.

To address this issue and further improve T-DNA vector design, we selected a panel of constitutive promoters from the *Arabidopsis thaliana* genome to evaluate their suitability for replacing *p35S* (see **Methods** for design criteria). To assess enhancer activity on a neighbouring gene, we introduced a minimal promoter (*p35Smin*) in a divergent orientation driving mStayGold expression, following the design from Jores et al^43^ (**Fig. 4a**). Any enhancer activity from the upstream promoter would upregulate mStayGold expression, allowing simultaneous quantification of promoter strength via mScarlet-I3 and unwanted enhancer effects via mStayGold. As benchmarks, we included the CaMV 35S regulatory elements (*35S*) to represent strong enhancer activity and the *Agrobacterium* nopaline synthase regulatory elements (*nos*) to define a lower expression boundary, reflecting the reduced kanamycin resistance typically observed when *nos* elements are used to drive the selection marker^44^.

**Fig. 4:**
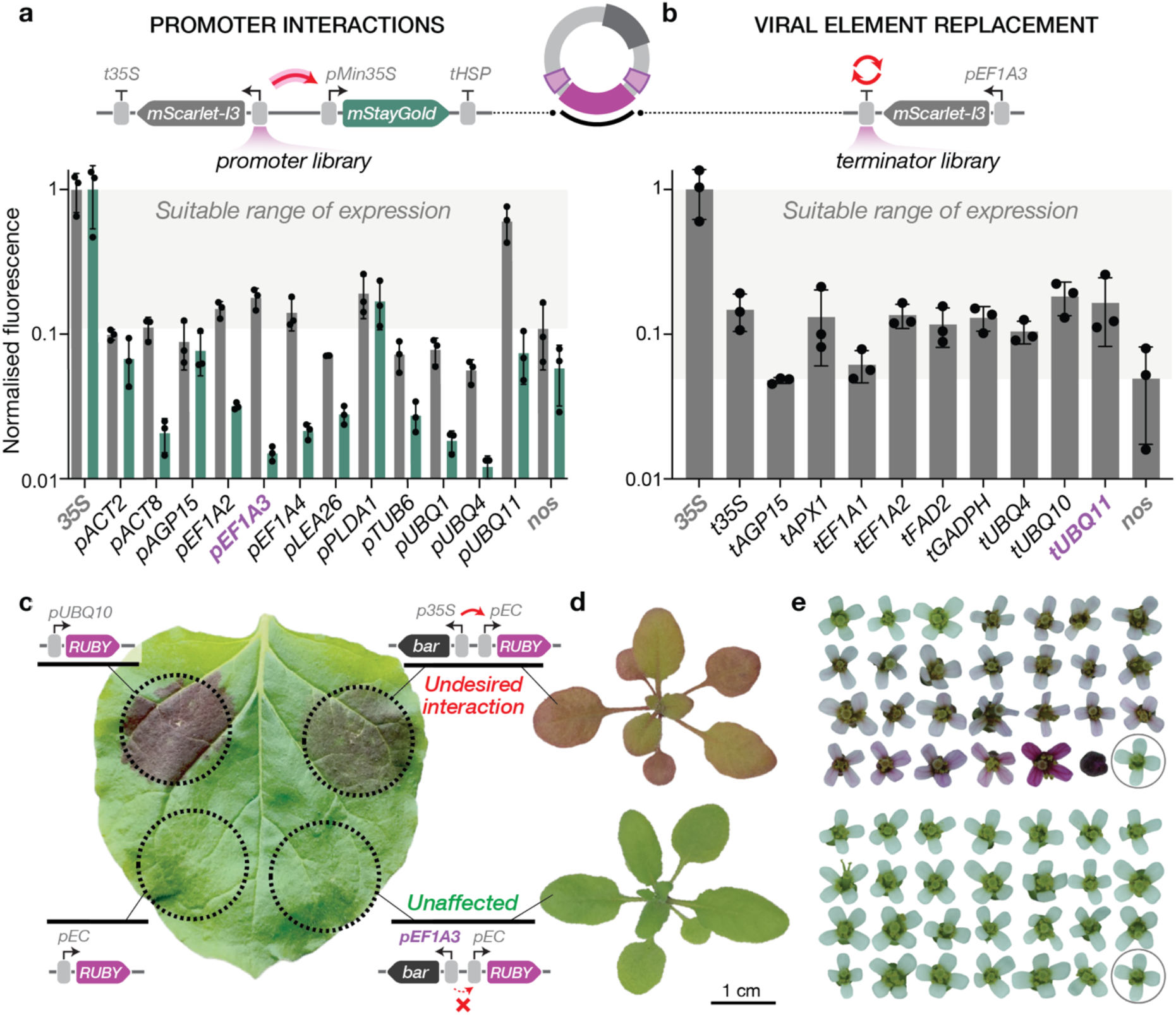
Identification of Arabidopsis regulatory elements that lack inappropriate enhancer activity. **a**, mScarlet-I3 (grey) and mStayGold (green) fluorescence from *N. benthamiana* leaf infiltrations of a panel of Arabidopsis promoters driving mScarlet-I3, cloned in a divergent orientation relative to a minimal 35S promoter (*p35Smin*) driving mStayGold to assay enhancer activity. The 35S terminator (*t35S*) was used for all Arabidopsis promoter constructs. **b**, mScarlet-I3 (grey) fluorescence from *N. benthamiana* infiltrations of the *EF1A3* promoter paired with a panel of Arabidopsis terminators. *p35S:mScarlet-I3:t35S* (*35S*) and *Pnos:mScarlet-I3:Tnos* (*nos*) were included as high- and low-expression controls, respectively. The grey shaded region indicates the desired range of expression above *nos*. Data points represent individual measurements from three independent infiltrations (averaging two leaf discs per infiltration), quantified by plate reader at 3 DPI and normalised to *35S* and non-infiltrated tissue. **c**, Representative *N. benthamiana* leaf at 5 DPI showing the infiltrated area (black dotted circle) for the indicated constructs. **d**, Representative *Arabidopsis thaliana* transformants of the indicated constructs at 21 DPG. *N. benthamiana* and Arabidopsis constructs were assembled in the pRB-Max and pRB-Min T-DNA backbone, respectively. **e**, Representative flowers from 27 randomly selected transgenic lines from the corresponding constructs in **d** (*p35S*, top; *pEF1A3*, bottom). The circled flower indicates WT.

We infiltrated these constructs into *N. benthamiana* leaves and quantified mScarlet-I3 and mStayGold fluorescence three days post-infiltration under long-day conditions (**Fig. 4a**). The tested promoters spanned a broad range of expression levels, with approximately half driving higher mScarlet-I3 expression than the *nos* regulatory elements. Among these, the *EF1A3* promoter (*pEF1A3*) exhibited the lowest enhancer activity, as evidenced by minimal mStayGold fluorescence that was indistinguishable from background, identifying it as a strong candidate for further development.

To eliminate foreign regulatory elements and create a fully Arabidopsis-derived selection cassette, we paired *pEF1A3* with a panel of Arabidopsis terminators. Transient expression assays showed that most terminators performed comparably to the CaMV 35S terminator (*t35S*), with all *pEF1A3*–terminator combinations yielding higher expression than the *nos* regulatory elements (**Fig. 4b**). The *UBQ11* terminator (*tUBQ11*) was selected for the final design because it is naturally located within an intergenic region and lacks common restriction enzyme recognition sites. We further confirmed that the new cassette does not exert repressive effects, demonstrating a genuine absence of enhancer activity (**Extended Data Fig. 13**).

Although the minimal promoter (*p35Smin*) is a convenient tool for quantifying enhancer activity, it does not reflect the behaviour of conditional or cell-type-specific promoters commonly used in plants. We therefore evaluated the redesigned selection marker cassette adjacent to the widely used egg cell–specific promoter, *pEC* (fusion of the *EC1*.*2* enhancer and *EC1*.*1* promoter), which underpins many CRISPR-based transformation strategies^45^. The original and redesigned selection cassettes were cloned in a divergent orientation upstream of the RUBY reporter. Following transient expression in *N. benthamiana*, the *pEF1A3*-based cassette showed no detectable RUBY expression, whereas the *p35S*-based design caused ectopic expression in leaves, consistent with strong enhancer activity (**Fig. 4c**). These results were recapitulated in stable Arabidopsis transformants generated using the pRB-Min T-DNA vector. In this context, RUBY expression was observed exclusively in plants carrying the *p35S*-driven selection marker and not in those containing the redesigned cassette (**Fig. 4d,e**). Egg cell-specific expression was further validated using a nuclear-localised mStayGold reporter (**Extended Data Fig. 14)**.

Together, these data demonstrate that, within the Arabidopsis genome and in the context of a cell-type-specific promoter, the redesigned selection marker lacks measurable enhancer activity on neighbouring gene expression. Importantly, the newly introduced Arabidopsis regulatory elements provided sufficient expression of the resistance marker, as evidenced by robust plant growth on standard concentrations of kanamycin (50 mg/L) when using the kanamycin resistance cassette, *pEF1A3:nptII:tUBQ11* (**Extended Data Fig. 15**). This design therefore enables reliable selection without compromising the specificity or predictability of transgene expression.

### An Arabidopsis-optimised T-DNA vector toolkit for clean, genomically mapped, single-copy integration

Having systematically explored how T-DNA vector design influences transformation outcomes and transgene expression, we next sought to integrate these advances into a unified, Arabidopsis-optimised T-DNA vector system. Because the combination of tandem LB sequences with the pRB-Mid backbone had not yet been evaluated, we first assessed the impact of this configuration (**Fig. 5a**). Consistent with our earlier observations, inclusion of tandem LBs (2xLB) substantially reduced vector backbone integration compared with a single LB (1xLB), with only 24% of transformants containing detectable backbone sequences, of which just 8% were attributable to LB read-through. Although the tandem LB design resulted in a slight reduction in overall transformation efficiency (18% lower), this decrease fell within the variability we typically observe between independent experiments. Importantly, the ratio of red to green plants remained unchanged, indicating that the desired low-copy integration behaviour was preserved. Indeed, copy number analysis confirmed that the resulting pRB-Mid vector with tandem LB sequences (pRB-Mid-2LB) produced predominantly single-copy insertions, with a median copy number of 1, compared with a median of 4 for pCAMBIA (**Fig. 5b**).

A longstanding challenge in plant transformation is the efficient identification of clean, single-copy insertion events^46^. To address this, we incorporated novel sequence elements inside of each T-DNA border that enable straightforward mapping of insertion sites by inverse PCR (iPCR) (**Fig. 5c**). These regions consist of 100-bp biologically neutral spacer sequences that buffer against the frequent truncations observed at T-DNA–plant genome junctions^47^, followed by NsiI restriction sites positioned adjacent to unique PCR barcode sequences (F, forward primer; R, reverse primer)^48^. NsiI was selected because it is predicted to generate a majority of fragments within the 0.1-3 kb range^49^ and is insensitive to cytosine methylation, facilitating reliable digestion across diverse genomic contexts (**Fig. 5d**). We first validated this strategy using wild type Arabidopsis genomic DNA, targeting native amplicons expected to arise following NsiI digestion and circularisation of fragments up to at least 4.8 kb. All three designed fragments were robustly amplified, demonstrating the efficiency and practicality of this approach for capturing large genomic regions flanking T-DNA insertion sites (**Fig. 5e**).

**Figure. 5:**
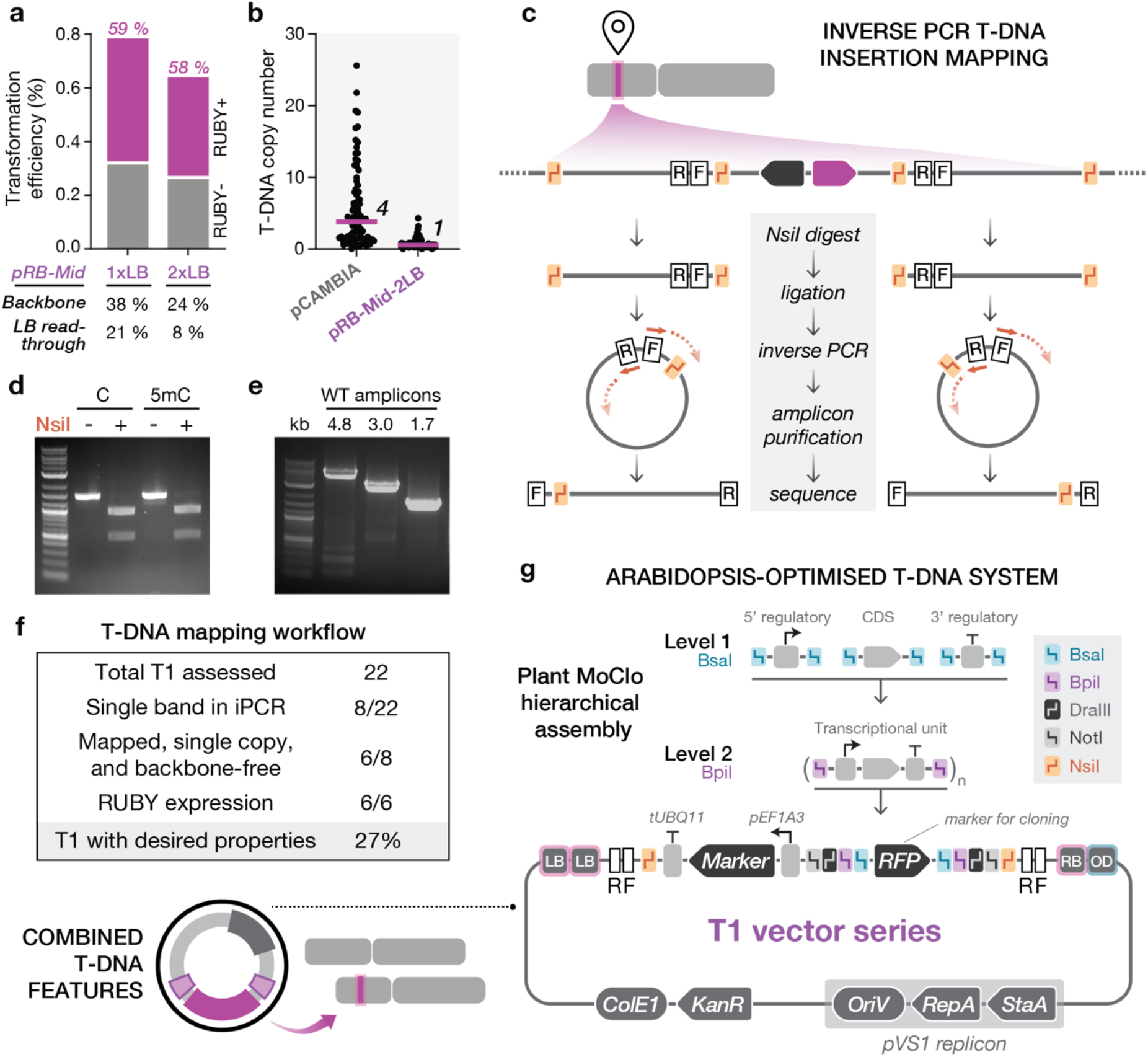
An Arabidopsis-optimised single-copy T-DNA vector system. **a**, Transformation efficiency of the pRB-Mid T-DNA vector with one (1xLB) or two (2xLB) sequences, scoring the RUBY+ and RUBY-plants, and showing the presence of backbone and LB read-through, as determined by low-cycle PCR. Percentage of RUBY+ plants shown in pink. **b**, T-DNA copy number of pRB-Mid with tandem LB sequences (pRB-Mid-2LB) in comparison to pCAMBIA. Pink horizontal lines denote median values. Numerical labels indicate the corresponding median, rounded to one significant figure. **c**, NsiI restriction sites are shown in orange. F and R denote forward and reverse primer binding sites, respectively, and orange arrows indicate primer binding and orientation. **d**, Digestion of a 1.5 kb amplicon containing cytosine (C) or 5-methyl cytosine (5mC) with (+) and without (-) NsiI. **e**, iPCR of circularised WT genomic DNA up to 4.8 kb in length. DNA ladder is NEB 1kb plus. **f**, Mapping of T-DNA insertion events using the iPCR workflow from 22 randomly selected transformants using pRB-Mid-2LB vector. **g**, Single-copy, Arabidopsis-optimised T-DNA vector toolkit, showing hierarchical plasmid assembly, based on the MoClo Golden Gate format. Plasmid architecture of T-DNA vector shown with preassembled selection marker, RFP dropout for cloning in *E. coli*, and highlighting key restriction enzymes.

When combined with the other features of our T-DNA vector, this mapping strategy enables rapid identification of single-copy T-DNA insertion events, supporting a streamlined workflow in which only a small number of plants need to be screened. This substantially reduces the extensive downstream profiling that has long bottlenecked plant research. We performed iPCR on 22 randomly chosen (RUBY+ and RUBY-) T1 lines transformed with *pUBQ10:RUBY:tHSP* in the pRB-Mid-2LB vector carrying the kanamycin resistance cassette, *pEF1A3:nptII:tUBQ11*. This identified eight lines exhibiting a single amplicon at both the LB and RB junctions, of which six could be precisely mapped at both borders, with the longest recovered amplicon spanning 6.2 kb (**Fig. 5f, Extended Data Table 2 and Extended Data Fig. 16**,**17**). All six mapped lines displayed consistent RUBY expression, resulting in 27% of transformants possessing all desired characteristics: mappable, backbone-free, single-copy insertions with stable transgene expression.

Integrating all previous optimisations, we next developed a new T-DNA vector cloning system based on the widely adopted MoClo framework, retaining the established overhangs and hierarchical Golden Gate assembly workflow for full compatibility^26^ (**Fig. 5g**). We designate these new T-DNA binary plasmids the “T1 vector series”, reflecting their ability to generate predominantly one T-DNA insertion that can be readily identified by straightforward mapping in the T1 generation. The T1 vector series is supported by a complete MoClo-compatible cloning toolkit that enables assembly of up to seven transcriptional units in two straightforward Golden Gate cloning steps, with iterative expansion allowing construction of even larger multigene assemblies. To improve accessibility, the system has been streamlined and simplified for ease of use while retaining the full functionality and compatibility of the original MoClo framework^26^. A detailed description of the toolkit is provided in the **Supplementary Information**.

## DISCUSSION

In this study we systematically dissected how T-DNA binary vector architecture, plasmid biology, and regulatory element choice shape transformation outcomes in *Arabidopsis thaliana*. We integrated these insights into the rational design of a single-copy, Arabidopsis-optimised T-DNA vector system, termed the T1 vector series. A central technical advance of this work was leveraging RUBY transgene silencing as a high-throughput phenotypic reporter for T-DNA copy number. Rather than relying solely on transformation efficiency, which obscures critical outcomes such as copy number and transgene stability, we show that RUBY expression robustly reflects these properties. Red plants primarily harbour a single T-DNA, providing a simple, visually scorable proxy for favourable transformation outcomes and enabling experiments to be scaled in ways that were previously impractical.

Leveraging this visual reporter alongside simple PCR analysis, we assayed changes to T-DNA vector architecture at scale, allowing systematic screening of multiple border variant architectures. These experiments conclusively demonstrate that the overdrive sequence is a dominant determinant of both T-DNA copy number and backbone integration in Arabidopsis, highlighting its importance as a “recruitment hub” for the virulence machinery. Guided by this insight, we engineered a suite of vectors with variable T-DNA transformation efficiencies and evaluated their performance across a wide range of experimental conditions. Among these, pRB-Mid, which incorporated a minimal overdrive motif, provided the most favourable balance, achieving moderate transformation efficiency while maintaining predominantly single-copy integration and minimal backbone transfer.

By exploiting mutations in the pVS1 replication protein RepA, we directly linked plasmid copy number in *Agrobacterium* to T-DNA integration behaviour. Increasing plasmid copy number enhanced transformation efficiency, but at the cost of elevated T-DNA copy number in stable transformants. Notably, our designed pRB-Mid vector was able to buffer against this effect, displaying comparable transformation efficiencies (with the R106H mutation) to that of unmodified pCAMBIA, while maintaining low-copy T-DNA integration. This underscores the complex and context-dependent nature of interactions between the virulence machinery, border sequences, and relative number of T-DNA copies within the bacterial cell.

More broadly, these findings may reflect a general principle underlying *Agrobacterium-mediated* transformation. Transformation efficiency appears to be limited primarily by successful infection of the target cell, rather than by the absolute amount of T-DNA delivered. Once infection has occurred, further increases in T-DNA transfer yield diminishing returns and become counterproductive. Under these conditions, restrained T-DNA delivery is not only sufficient but preferable, as it promotes low-copy integration. This work demonstrates the importance of evaluating transformation outcomes holistically, as improvements in one property can substantially compromise another.

By coupling a predominantly single-copy T-DNA vector with a streamlined insertion-mapping workflow, we enable the rapid identification of clean, single-copy integration events from a small pool of transformants. This will help avoid genome positional effects and reduces the likelihood of transgene silencing, resulting in more predictable and stable expression. In parallel, redesigning the selection marker to eliminate unwanted CaMV 35S enhancer activity increases confidence in conditional and cell-type-specific transgene regulation. Together, this platform substantially reduces the extensive downstream screening and characterisation that have long represented a major bottleneck in plant transformation **(Fig 6)**.

**Fig. 6:**
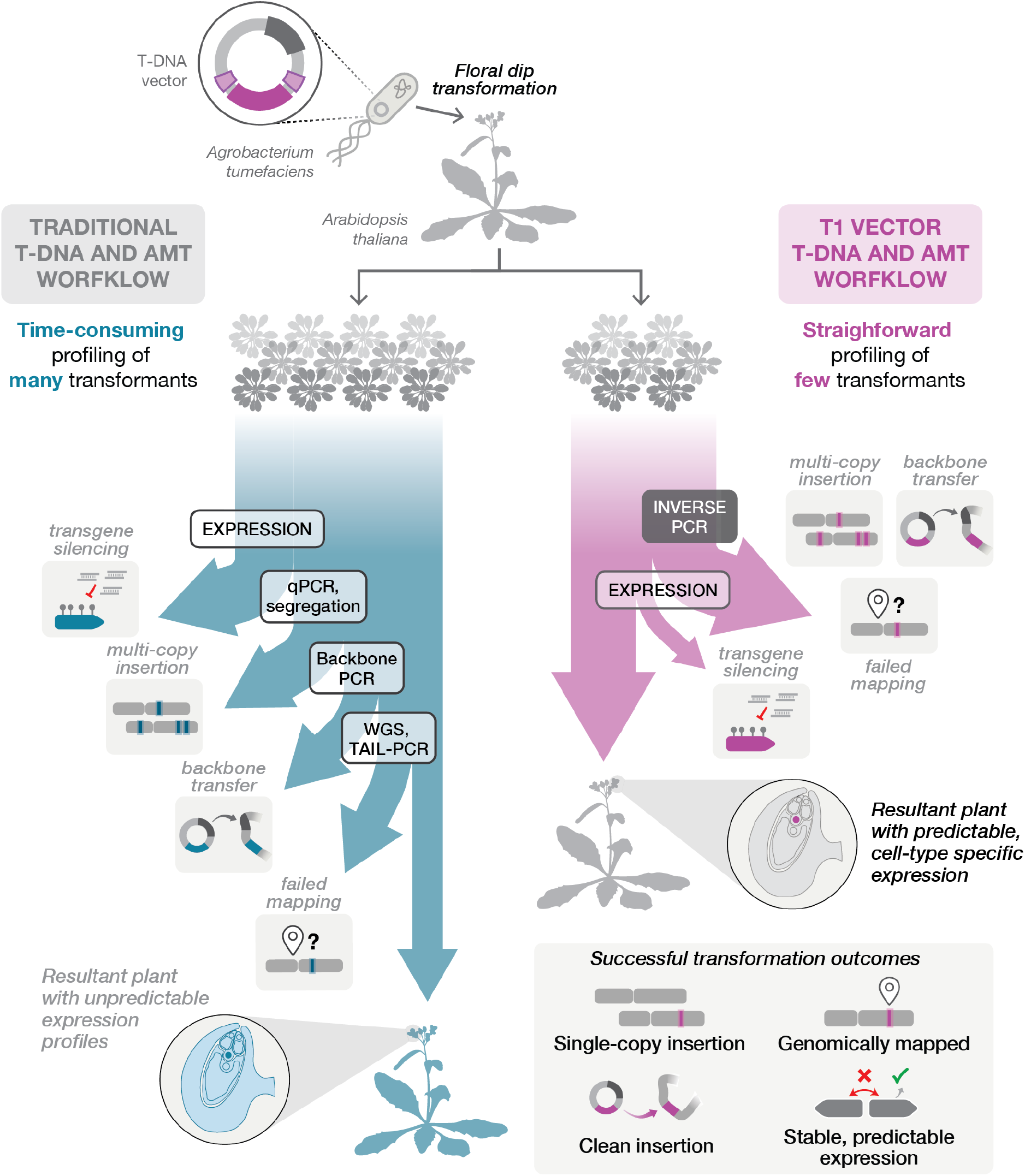
Simplified workflow for generating single-copy, mapped T-DNA lines in *A. thaliana* using the T1 vector. **Left**, Traditional *Agrobacterium*-mediated transformation (AMT) workflow using standard T-DNA vectors, highlighted in blue. Large numbers of transformants must be screened to identify lines carrying single-copy, stable transgene insertions due to frequent multi-copy integration, backbone incorporation, and transgene silencing. In addition, the enhancer activity of the widely used CaMV 35S promoter, which drives selection marker expression in many vectors, can lead to unpredictable cell-type-specific or condition-dependent transgene expression. **Right**, AMT workflow using the T1 vector, highlighted in pink. Fewer transformants are required to identify fully mapped, backbone-free, single-copy insertion events due to predominantly single-copy integration behaviour and simplified genomic mapping. The redesigned selection marker minimises enhancer interference, resulting in more predictable cell-type-specific and condition-dependent transgene expression.

This study focuses on *Arabidopsis thaliana*, the central model for plant biology and an important chassis for plant synthetic biology. How broadly these optimisation strategies translate to other species remains to be determined, particularly given the unique target cell and infection route exploited by the floral dip method. However, the identification of vector features that tune transformation quality provides a clear basis for such optimisation. In particular, our work identifies the overdrive sequence as a principal factor shaping transformation outcomes, accounting for wide-ranging differences among vectors that otherwise share the same backbone sequence. Further dissection and optimisation of this virulence element, together with systematic testing across a broader diversity of Agrobacterium strains, may provide additional opportunities to improve plant transformation.

Nevertheless, the T1 vector series achieves all criteria for successful transgenesis in Arabidopsis. By addressing long-standing challenges in *Agrobacterium*-mediated transformation, this work shifts the emphasis of T-DNA vector design from maximising efficiency to rationally engineering the quality of resulting transgenic lines. In doing so, it establishes an outcome-focused framework for developing more predictable and effective plant transformation across diverse species.

## METHODS

### *E. coli* strains, culture conditions, and transformation

*Escherichia coli* strain TOP10 (Thermo Fisher Scientific) was used for all cloning experiments. Cultures were grown in Lysogeny Broth (LB) medium at 37 °C with shaking at 225 rpm. Except during preparation of competent cells, LB medium was supplemented with the appropriate antibiotics for plasmid maintenance: chloramphenicol (34 mg/L), carbenicillin (100 mg/L), kanamycin (50 mg/L) or spectinomycin (100 mg/L). Chemically competent *Escherichia coli* cells were prepared using the TSS method for KCM transformation as previously described^50^.

### Plasmids used in this study

T-DNA vector variants described in this work were derived from pAGM4723 (Addgene ID: 48015)^26^. Additional mutations were introduced where necessary to remove all occurrences of common Type IIS and Type II restriction enzyme recognition sites, generating a vector free of the following sites: Type IIS enzymes (BsaI, BpiI, BsmBI, PaqCI, and SapI) and Type II enzymes (BamHI, BglII, EcoRI, HindIII, PstI, SpeI, XbaI, XhoI, SphI, KpnI, SalI, and SacII). The sequence between the T-DNA borders was replaced by synthesised DNA (Twist Biosciences) containing an mScarlet3 cloning dropout marker derived from YTK, using the MoClo formatting. 100 bp biological neutral spacers (GC 50) were included adjacent to the borders to buffer against truncation commonly seen in T-DNA integration^47^, designed using R2oDNA Designer^51^. The DNA barcodes for inverse PCR were chosen from Subramanian et al.^48^, first screening the sequences with low similarity to the reference *Arabidopsis thaliana* genome (TAIR10) and then choosing a smaller set based on their secondary structure and compatibility (manually assessed in Benchling). Primer pairs were validated in vivo, and the best combinations showing the greatest specific and lowest non-specific amplification were chosen. The remaining plasmid toolkit was adapted from Multiplex Yeast Toolkit (MYT)^52^, incorporating the overhangs from the MoClo system^26^. pCAMBIA was taken from Szarzanowicz et al.^18^, and recreated for this study using DNA synthesis (Twist Bioscience). For experiments using pCAMBIA with RUBY, the sequence between the T-DNA borders was transferred directly from T-DNA plasmid in this study. All cloning was performed using Golden Gate Assembly and validated using whole plasmid sequencing (Plasmidsaurus). Plasmids used in this work are listed in **Supplementary Table 1**. For a list of MoClo toolkit plasmids, see **Supplementary Table 2**.

### Golden Gate assembly

All DNA insert and plasmid concentrations were set to 50 fmol/μL and plasmid backbone concentrations were set to 25 fmol/μL prior to assembly. Golden Gate reaction mixtures were prepared as follows: 0.5 μL of each DNA insert or plasmid, 1 μL of T4 DNA Ligase Reaction Buffer (NEB), 0.5 μL of T4 DNA Ligase (400 U/μL, NEB), 0.5 μL of Type IIS restriction enzyme, and water to bring the final volume to 10 μL. The Type IIS restriction enzymes used were either BsaI (20 U/μL BsaI-HF v2, NEB), BpiI (10 U/μL, Thermo Fisher Scientific), or BsmBI (10 U/μL BsmBI v2, NEB). Reaction mixtures were incubated in a thermocycler according to the following program: 25 cycles of digestion (BsaI/BpiI, 37 °C for 5 min or BsmBI, 42 °C for 5 min) and ligation (16 °C for 5 min), followed by a final digestion step (55 °C for 10 min) and a heat inactivation step (80 °C for 10 min). The entire reaction was then transformed into *E. coli* and plated on LB medium plus the appropriate antibiotic.

### *Agrobacterium* strains, culture conditions, and transformation

Unless otherwise stated, *Agrobacterium tumefaciens* strain GV3101 (GoldBio) was used for all experiments. For experiments comparing strains, *A. tumefaciens* EHA105, LBA4404, and AGL-1 (GoldBio) were used. GV3101, EHA105 and AGL-1 were cultured in LB medium (Research Products International, L24400), and LBA4404 was grown in YEP medium (10 g/L yeast extract + 20 g/L peptone) at 28 °C with shaking at 225 rpm. Except during preparation of competent cells, growth medium was supplemented with antibiotics for strain selection and plasmid maintenance as follows: gentamicin (30 mg/L) for GV3101; rifampicin (15 mg/L) for LBA4404 and EHA105; carbenicillin (50 mg/L) for AGL-1; and kanamycin (50 mg/L) for plasmid maintenance in all strains. Cells were transformed by electroporation at 1.8 kV in an ECM 399 Electroporation System (BTX) and transferred to 1 mL of LB in a 1.5 mL tube. Cells were then incubated at 28 °C, shaking at 225 rpm for 3h, before plating 100 µL on appropriate medium and antibiotics and incubated for 3 days at 28 °C. *Agrobacterium* growing on plates were stored at 4° C for no longer than 2 weeks after transformation.

### Plant growth and selection

Plants were grown in long day conditions (16 h light, 8 h dark) at 21 °C under ~120 µmol/m^2^ full spectrum LED lights. Plants were grown on soil mix containing LM-2 Germination Mix (Lambert) and vermiculite (Whittemore Co) in a 2:1 ratio with 14-14-14 slow-release fertiliser (Osmocote). After sowing, seeds were stratified at 4 °C in the dark for 3 days to break dormancy. Seeds were then germinated under a humidity dome, which was cracked after 5 days for *Arabidopsis thaliana* or 6 days for *Nicotiana benthamiana*, and removed the following day. For BASTA selection, Arabidopsis plants were grown on soil in 10 × 20-inch flats at various seed densities (100-400 mg seed). BASTA (BASTA 120 mg/L + 0.1 % Silwet L-77) was sprayed onto seedlings at 6 DPG at 2-day intervals for 2 weeks. For kanamycin selection, Arabidopsis plants were grown on 0.5x MS medium, pH 5.7 (2.15 g/L Murashige and Skoog Basal Salt Mixture (Sigma-Aldrich M5524) + 7% agar), supplemented with 50 mg/L kanamycin and 50 mg/L Timentin to reduce *Agrobacterium* growth. Before plating, Arabidopsis seeds were surface sterilised using ethanol and Triton X-100 (0.05%).

### Arabidopsis floral dip

*Arabidopsis thaliana* plants were grown in 3-inch pots at a density of four plants per pot, with two pots typically used per transformation. Approximately 4–5 days after bolting, the primary inflorescence was removed to stimulate secondary floral growth. Plants were transformed 1 week later. Two days before transformation, a starter culture of *Agrobacterium* was inoculated into 2 mL of growth medium supplemented with the appropriate antibiotics in a 14 mL culture tube and incubated at 28 °C with shaking at 225 rpm. The following afternoon, the culture was back-diluted 1:100 into 100 mL of antibiotic-supplemented growth medium in a 500 mL flask and grown overnight at 28 °C with shaking at 225 rpm. The next morning, 50 mL of saturated culture was pelleted at 3,500 g for 10 min at 4 °C. Cells were resuspended in 50 mL transformation solution (5 % sucrose, 0.02 % Silwet L-77) and incubated at room temperature for 1 h, shaking at 225 rpm. Unless otherwise stated, the suspension was adjusted to an OD600 of 0.8 following resuspension. Immediately before dipping, all open flowers and developing siliques were removed. Plants were submerged briefly in the *Agrobacterium* suspension to ensure full coverage of floral buds, then placed horizontally between closed opaque trays for 24 h at room temperature. A second floral dip was performed 1 week later. Plants were watered for 2 weeks following the second dip, then allowed to dry on the growth shelf. Seeds were harvested approximately 2 weeks later after complete drying of the plants.

### Arabidopsis transformation counts

Arabidopsis transformants were scored 21 DPG following BASTA selection. Resistant plants were removed from the soil tray and visually inspected for RUBY pigmentation. Plants exhibiting any visible red pigmentation were classified as RUBY+, whereas plants lacking detectable pigmentation were classified as RUBY−. All scoring was performed manually by eye. A variable number of seeds was used for each experiment, with a minimum of 10,000 seeds screened in all cases. See **Extended Data Table 1** for details on seed amount used and transformation counts.

### *Nicotiana benthamiana* transient infiltration

*Nicotiana benthamiana* plants were grown individually in 3-inch pots for approximately 4 weeks prior to infiltration. Two days before infiltration, a starter culture of *Agrobacterium* was inoculated into 2 mL of LB growth medium supplemented with the appropriate antibiotics in either a 14 mL culture tube or a 12-well plate sealed with a breathable membrane and incubated at 28 °C with shaking at 225 rpm. *Agrobacterium tumefaciens* GV3101 was used for all experiments. The following afternoon, cultures were back-diluted 1:100 into 2 mL of fresh antibiotic-containing growth medium and grown overnight under the same conditions. On the day of infiltration, cultures were pelleted at 3,500 g for 10 min and resuspended in 2 mL infiltration solution (MMA buffer: 10 mM MES-KOH, 10 mM MgCl_2_, 100 µM acetosyringone, pH 5.6). Suspensions were incubated at room temperature for 2 h with shaking at 225 rpm and adjusted to an OD600 of 0.8 prior to infiltration. Intermediate-sized leaves were infiltrated using up to four infiltration spots per leaf, each approximately 2–3 cm in diameter, marking the area with a black marker. Following infiltration, plants were kept in the dark at room temperature for 16 h before being returned to standard growth conditions.

### Leaf disc fluorescent plate reader measurements

*Nicotiana benthamiana* leaf discs were collected from infiltrated regions using an 8 mm biopsy punch (Electron Microscopy Sciences). Two discs per infiltration were placed abaxial side down onto an inverted clear 96-well microplate lid (Corning 3931), using the condensation rings to position and retain the discs, with 3 µL of water added to aid adhesion. Discs were secured in place with a strip of white laboratory tape applied gently to the adaxial surface, to cover the entire bottom of the lid. The microplate lid was then placed into a SpectraMax M5 plate reader (Molecular Devices) such that fluorescence was measured from the abaxial side of the leaf discs through the clear lid. Fluorescence measurements were acquired using the following settings: mScarlet-I3, excitation 569 nm and emission 590 nm; GFP/mStayGold, excitation 485 nm and emission 515 nm; automatic gain enabled. Fluorescence values from the two discs corresponding to each infiltration were averaged prior to analysis.

### Co-extraction of genomic DNA and betalain using the DNeasy 96 Plant Kit

To enable quantitative assessment of RUBY expression while simultaneously isolating genomic DNA for copy number analysis, we established a high-throughput workflow based on the QIAGEN DNeasy 96 Plant Kit, that extracts genomic DNA in a 96-well format (**Extended Data Fig. 2a**). Building on our prior observations that betalain is recovered in the initial supernatant, following Buffer P3 addition and centrifugation, we hypothesised that absorbance of this soluble fraction could serve as a direct proxy for RUBY expression, similar to extracted samples^53^. To validate this approach, we conducted a pilot experiment extracting genomic DNA in parallel with RNA purification from eight plants with varying levels of red pigmentation (**Extended Data Fig. 2b-d**). Absorbance measurements at 538 nm of the supernatant showed strong correlation with RT–qPCR quantification of RUBY mRNA abundance (R^2^ = 0.9), confirming the robustness of this assay (**Extended Data Fig. 2e**).

The protocol is as follows: betalain was quantified during genomic DNA extraction using the DNeasy 96 Plant Kit (QIAGEN), using the initial supernatant. 50 mg of leaf tissue were lysed in 400 µL of working lysis solution and a 3 mm stainless steel bead and disrupted using a TissueLyser II (QIAGEN) for 90 s at 30 Hz. Tube racks were then reoriented and disrupted for an additional 90 s at 30 Hz to ensure complete homogenisation. Following tissue disruption, 130 µL Buffer P3 was added to each sample. Tubes were capped, mixed by shaking for 15 s, and incubated at −20 °C for 10 min. Samples were centrifuged for 5 min at 4,000 g, and 400 µL of clarified supernatant was transferred to a fresh collection tube. 200 µL of the supernatant was transferred to a clear-bottom 96-well plate and absorbance was measured at 538 nm using a SpectraMax M5 plate reader (Molecular Devices). Following measurement, the supernatant was transferred back to the tube, and the remaining genomic DNA extraction procedure was then completed according to the manufacturer’s instructions.

### RT-qPCR

Total RNA was extracted from 100 mg of leaf tissue harvested at 21 DPG using the RNeasy Plant Mini Kit (QIAGEN) with freshly added β-mercaptoethanol in Buffer RLT. Following extraction, samples were treated with DNase I for 10 min at room temperature and immediately purified using the RNA Clean & Concentrator kit (Zymo Research). RNA concentration was measured using a NanoDrop spectrophotometer, and 500 ng of total RNA was reverse transcribed in a 20 µL reaction using RevertAid Reverse Transcriptase (Thermo Scientific) with random hexamer primers. qPCR reactions were performed in 10 µL volumes containing 1 µL cDNA, 5 µL LightCycler 480 SYBR Green I Master mix (Roche), and 0.5 µM of each cDNA specific primer for RUBY CDS (FP: CTACCGTGATCAAGAATGGATC; RP: CAGCTTGGTGGTCTGCTG) or the *Arabidopsis thaliana UBC9* reference gene (FP: TCACAATTTCCAAGGTGCTG; RP: GTGGACTCGTACTTGTTCTTGTC). qPCR reactions were run on a LightCycler 480 instrument (Roche) and Ct values were calculated using the Abs Quant/2nd Derivative Max method in the LightCycler480 Software. RUBY expression levels were normalised to *UBC9*.

### Arabidopsis T-DNA copy number analysis

3 µL of genomic DNA (~10 ng) extracted using the DNeasy Plant 96 Kit (QIAGEN) was used in a 10 µL qPCR reaction containing 5 µL LightCycler 480 SYBR Green I Master mix (Roche) and 0.5 µM of each primer targeting a single *Arabidopsis thaliana* genomic locus (AT2G37620) (FP: GGAACATCCTATTCTACTTACCGAG; RP: AACAGCCTGAATAGCCACATAC) or the RUBY coding sequence (FP: CACTCATTTCCCTCAGACTCG; RP: CACTATTCGGGATCGTGC). qPCR reactions were run on a LightCycler 480 instrument (Roche) and Ct values were calculated using the Abs Quant/2nd Derivative Max method in the LightCycler480 Software. Relative T-DNA copy number was determined using the ΔΔCt method^54^, calibrated against independently validated single-copy and double-copy insertion lines.

### *Agrobacterium* T-DNA plasmid copy number analysis

Plasmid copy number in *Agrobacterium* was determined using the direct qPCR method described by Song *et al*.^55^. *Agrobacterium* cultures were inoculated into 2 mL LB medium containing 50 µg/mL kanamycin in a 14 mL culture tube and grown overnight at 28 °C with shaking at 225 rpm. The following day, cultures were back-diluted 1:100 into fresh selective medium and grown for an additional 16 h under the same conditions. For crude lysate preparation, 200 µL of saturated culture was pelleted in a 1.5 mL tube at 20,000 g for 3 min and resuspended in 800 µL of 0.5% IGEPAL. 100 µL was transferred to a PCR tube and incubated at 98 °C for 10 min to lyse cells. 4 µL of crude lysate was then used directly in a 20 µL qPCR reaction containing 10 µL LightCycler 480 SYBR Green I Master mix (Roche) and 0.5 µM of each primer targeting the *Agrobacterium tumefaciens* genome (FP: GAGTACCGGAATCTCGTCAAAGCC; RP: CGAAGATCTCTACGGCAACTACCTGG) or the T-DNA plasmid (FP: GATCATCCTGATCGACAAGACCGG; RP: CTGCCGAGAAAGTATCCATCATGGC). qPCR reactions were run on a LightCycler 480 instrument (Roche) and Ct values were calculated using the Abs Quant/2nd Derivative Max method in the LightCycler480 Software. Relative plasmid copy number was calculated by normalising plasmid Ct values to the *Agrobacterium* genomic Ct values (2^-(Ct(plasmid)-Ct(genome)) and then expressing Ct values relative to the wild type pCAMBIA plasmid.

### Low-cycle PCR detection of T-DNA backbone integration

Low-cycle PCR amplification of vector backbone sequences was performed using the Phire Plant Direct PCR Kit (Thermo Scientific). Leaf discs (~0.5 cm diameter) were collected from plants at 21 DPG and transferred into 1.2 mL 96-well cluster tubes (Corning) containing a single 3 mm stainless steel bead. Samples were sealed, frozen at −80 °C, and disrupted using a TissueLyser II (QIAGEN) for 1 min at 30 Hz. Following disruption, 50 µL of Phire Dilution Buffer was added to each well and mixed by pipetting. The entire volume was transferred to a PCR plate, sealed, and incubated at 98 °C for 10 min in a thermocycler. 1 µL of crude lysate was then used directly in a 10 µL PCR reaction containing 5 µL Phire Plant Direct PCR Master Mix and 0.5 µM of each primer (Backbone, FP: CGTGAGTTTTCGTTCCACTG, RP, CTTACCGGATACCTGTCCG; LB read-through, FP: CCACCACTTCAAGAACTCTGTAG, RP: TGCTCAGAACTCACGACTCC). PCR reactions were performed using the following cycling conditions: 98 °C for 5 min, followed by 28 cycles of 98 °C for 10 s, 61 °C (backbone) or 63 °C (LB read-through) for 10 s, and 72 °C for 30 s. PCR products (5 µL) were analysed by agarose gel electrophoresis.

### Inverse PCR mapping

Genomic DNA was extracted from 100 mg of leaf tissue using the DNeasy Plant Mini Kit (QIAGEN) and eluted in 100 µL Buffer EB (QIAGEN; 10 mM Tris-Cl, pH 8.5). DNA concentration was quantified using a NanoDrop spectrophotometer. For restriction digestion, 500 ng of genomic DNA was digested with NsiI-HF (NEB) in a 100 µL reaction containing 1 µL enzyme and 10 µL CutSmart Buffer (NEB). Reactions were incubated at 37 °C for 12 h, followed by heat inactivation at 80 °C for 20 min. The entire digest was then used in a 200 µL ligation reaction containing 1 µL T4 DNA ligase (NEB) and 20 µL T4 DNA ligase buffer (NEB), and incubated at room temperature for 2 h, followed by heat inactivation at 65 °C for 10 min. Circularised DNA was purified and concentrated using the Monarch PCR & DNA Cleanup Kit (5 µg) (NEB), eluted in 20 µL elution buffer, and measured using a NanoDrop spectrophotometer. Inverse PCR (iPCR) was performed using 25 ng circularised DNA in 50 µL reactions containing 25 µL Phire Direct PCR Master Mix (Thermo Scientific) and 0.5 µM of each primer for LB junction (FP: TTCAGCACCACGAAATAGTAGC, RP: CAGTCCCGAGCCGTCTTACTC) or RB junction (FP: CCTCGTGAATAATCAGGGTGAC, RP: CTATTGTAAAGCGAAGTGTAAGGTG). PCR reactions were performed using the following cycling conditions: 98 °C for 5 min, followed by 35 cycles of 98 °C for 10 s, 62 °C for 10 s, and 72 °C for 2 min 30 s. 5 µL of each PCR product were analysed by agarose gel electrophoresis. For lines exhibiting a single amplification band at both the LB and RB junctions, the remaining PCR product was gel purified and sequenced (Plasmidsaurus). Amplicon sequences were aligned to the *Arabidopsis thaliana* reference genome (TAIR10), and genomic coordinates were imported into Benchling for co-alignment of LB and RB junctions. Successful mapping was validated by colocalisation of LB and RB junctions and termination of amplicons at predicted NsiI restriction sites.

### Small RNA sequencing

Leaf tissue was snap-frozen in liquid nitrogen and ground using a TissueLyser II (QIAGEN). Material was maintained frozen on dry ice prior to the extraction of total RNA utilising Plant RNA Isolation Aid (Invitrogen AM9690) in combination with RNAqueous-Micro Kit (Invitrogen AM1931) according to manufacturer’s specifications. Libraries were prepared with the NEXTFLEX Small RNA Sequencing Kit V4 (Revvity NOVA-5132-31) using 0.5-2 µg total RNA as input and 17 cycles of PCR. Libraries were pooled and sequenced with Illumina NovaSeq S4 with 150 bp paired-end reads. Resulting reads were trimmed with cutadapt^56^ (-q 20 --max-n 0.5 -a TGGAATTCTCGGGTGCCAAGG -m 16 -M 40) and aligned to TAIR10 with the T-DNA construct *p35S:bar:t35S*(Reverse)-*pUBQ10:RUBY:t35S*(forward) appended as an additional chromosome using bowtie^57^ (*-v 2 --best -S*). Alignments were filtered to select for 21-24-nt sRNAs and normalised by library size.

### Enzymatic methylation sequencing

Genomic DNA was extracted from 100 mg of leaf tissue using the DNeasy Plant 96 Kit (QIAGEN) and eluted in 100 µL Buffer EB (QIAGEN; 10 mM Tris-Cl, pH 8.5). 100 ng of DNA was sheared in 50 µl of 1X TE buffer, 6 × 16 mm tubes under conditions [Peak power = 175 W, Duty factor = 10%, Cycles/burst = 200, Time = 80 sec]. Enzymatic conversion and library preparation was performed with Enzymatic Methyl-Sequencing Kit v1 (New England Biolabs), following the manufacturer’s instructions. Formamide was used for the denaturation step in the protocol, and 7 cycles of PCR were used to amplify libraries. Libraries were pooled and sequenced with Element Biosciences AVITI with 150 bp paired-end reads. Resulting reads were trimmed with cutadapt^56^ (-a AGATCGGAAGAGCACACGTCTGAACTCCAGTCA -A AGATCGGAAGAGCGTCGTGTAGGGAAAGAGTGT -m 30). Methylation analysis was performed using Bismark^58^ (--comprehensive --bedgraph --CX --ignore_r2 3) aligning to TAIR10 with the T-DNA construct *p35S:bar:t35S*(Reverse)-*pUBQ10:RUBY:t35S*(forward) appended as an additional chromosome using bowtie2^59^ (-L 10 -N 1 --non_directional). Bismark was used to remove PCR duplicates and generate methylation reports.

### Fluorescence microscopy of ovules

Flowers were emasculated at 28 DPG. Two days later, emasculated flowers were collected and pistils were dissected along both sides of the septum to expose the ovules. Dissected pistils were immediately transferred to 1.5 mL fixation solution (4% paraformaldehyde, 1× PBS, 0.05% Triton X-100) in 1.5 mL tubes. Samples were vacuum infiltrated on ice for 30 min and then incubated overnight at 4 °C with gentle rotation. The following day, pistils were washed three times with 1x PBS containing 0.05% Triton X-100 and transferred to 300 µL of freshly prepared ClearSee solution^60, 61^ (10% (w/v) xylitol, 15% (w/v) sodium deoxycholate, 25% (w/v) urea) supplemented with an additional 50 mM sodium sulphite. Samples were incubated at room temperature with gentle rotation for 7 days. Following clearing, ovules were dissected from the pistils and mounted in ClearSee solution on microscope slides. Images were acquired using a Zeiss microscope and processed using Zeiss ZEN software. Representative images are overlays of DIC and GFP fluorescence channels. Images were collected using a 20x objective with the following settings: DIC, 10 ms exposure at 5.25 V; GFP (38HE), 25 ms exposure. For image presentation, DIC exposure was increased by a factor of 1.35 and GFP brightness was adjusted to −20.

### Arabidopsis promoter and terminator design

We created a list of strongly expressed constitutive genes, including genes previously used as promoter sources and stable reference genes^26, 62–65^ (**Supplementary Table 3**). From this list, we applied rational filtering criteria to exclude candidate promoters with possible bidirectional transcription, overlapping gene bodies, or alternatively spliced 5′ UTRs, resulting in a final set of 12 promoters. Using leaf ATAC-seq data^66^, we then defined the putative minimal sequences necessary for full promoter activity based on open chromatin profiles. Promoter sequences were taken from the start of the open chromatin up to the annotated 5’ start of the coding sequence, including the full 5’ UTR region. These sequences were then domesticated for MoClo Golden Gate assembly by removing all BsaI and BpiI sites, synthesised (Twist Bioscience), and cloned upstream of the mScarlet-I3 coding sequence to measure promoter strength. We compiled a list of terminators from the same list of *Arabidopsis thaliana* genes (**Supplementary Table 4**). Terminators were selected where the sequence was in an intergenic region or in a convergent orientation with the neighbouring gene to minimise the risk of incorporating unwanted promoter elements, resulting in a list of nine terminators. The terminator region was defined as extending from the annotated 3′ end of the coding sequence to either 50 bp beyond the annotated 3’ UTR region (TAIR10) or the furthest experimentally measured transcription termination site^67^, whichever was longer. The resulting sequences were domesticated, synthesised (Twist Bioscience), and cloned downstream of the mScarlet-I3 coding sequence.

## RESOURCE AVAILABILITY

### Lead Contact

Further information and requests for resources and reagents should be directed to and will be fulfilled by the lead contact: Ahmad S. Khalil.

### Materials Availability

Key plasmids will be deposited at Addgene for distribution. DNA constructs will be available from the lead contact. Seeds will be available from Mary Gehring.

### Data and Code Availability

- Raw small RNA-seq and EM-seq data for transcriptome and DNA methylation analysis will be deposited in the NCBI GEO database. Accession numbers will be listed in key resources table. All other raw datasets will be deposited on Dryad. Associated DOIs will be listed in the key resources table.
- All original code will be available on Github and will be deposited on Zenodo. Associated DOIs will be listed in the key resources table.
- Any additional information required to reanalyse the data reported in this paper is available from the lead contact upon request.

## AUTHOR CONTRIBUTIONS

Conceptualisation, W.M.S., M.G., and A.S.K.; Experiments, W.M.S., A.G., E.I.T., L.L.B., and S.C.; Data analysis, W.M.S., A.G., E.I.T., and L.L.B.; Design of MoClo toolkit, W.M.S. and S.G.; Writing, W.M.S., M.G., and A.S.K.; Reviewing and editing, all authors.

## ACKNOWLEDGEMENTS

We thank Souraya Khouider for assistance with fluorescence microscopy, Dayishaa Daga for organisational support during this work, and the Whitehead Institute Genome Technology Innovation Center for sequencing services. This work was supported by Schmidt Sciences Polymath award G-22-63292 and the Vannevar Bush Faculty Fellowship (VBFF) under grant N00014-20-1-2825 to A.S.K. and funding from the Manton Foundation to M.G. This research was supported in part by a generous gift from Vin Ryan to Whitehead Institute in memory of Dr. Vincent J. Ryan. W.M.S. was supported by EMBO Postdoctoral Fellowship ALTF 1116-2020. S.G. and W.M.S. were supported by the UK Research and Innovation (UKRI) Biotechnology and Biological Sciences Research Council (BBSRC) via the Earlham Institute Core Capability Grant BB/CCG2220/1. S.G. was additionally supported by the UKRI Future Leader Fellowship MR/Y019008/1. A.G. and E.I.T. were supported by an NIH/NIGMS predoctoral training fellowship (grant no. T32GM154655). M.G. is an Investigator of the Howard Hughes Medical Institute.

