## Supplementary Information for "Rational design of T-DNA vectors enables predictable, single-copy integration in *Arabidopsis thaliana*"

#### EXTENDED DATA

**Extended Data Fig. 1:** Arabidopsis transformants from the four T-DNA arrangements in Fig. 1c,d.

**Extended Data Fig. 2:** Extraction of betalain using the Qiagen DNeasy Plant 96 kit

**Extended Data Fig. 3:** Transplanted green and red Arabidopsis transformants from Fig. 1e

**Extended Data Fig. 4:** Late silencing in Arabidopsis transformants from Fig 1e

**Extended Data Fig. 5:** Small RNA profiles of 6 green (RUBY-) and 2 red (RUBY+) plants

**Extended Data Fig. 6:** 5mC profiles of 6 green (RUBY-) and 2 red (RUBY+) plants

**Extended Data Fig. 7:** High prevalence of RUBY silencing cannot be explained by major features of the transcriptional unit

**Extended Data Fig. 8:** Mechanisms of T-DNA vector backbone transfer

**Extended Data Fig. 9:** Optimisation of a low-cycle PCR protocol for detecting presence of T-DNA events in Arabidopsis

**Extended Data Fig. 10:** Amplification of the ColE1 replicon to detect the integration of all backbone events in Arabidopsis transformants

**Extended Data Fig. 11:** Amplification across left border sequence (LB) to detect the integration of LB readthrough events in Arabidopsis transformants

**Extended Data Fig. 12:** Detection of backbone transfer in Arabidopsis transformants using the pRB-Mid vector

**Extended Data Fig. 13:** Evaluating the effect of the *EF1A3* promoter on strong constitutive transgene expression in the divergent arrangement

**Extended Data Fig. 14:** Egg cell-specific expression of nuclear-localised GFP using the *pEF1A3*-based selection marker system

**Extended Data Fig. 15:** Robust growth of antibiotic-resistant transformants using the *pEF1A3*-based selection cassette

**Extended Data Fig. 16:** Inverse PCR (iPCR) workflow and analysis of T-DNA insertion events in randomly selected Arabidopsis transformants

**Extended Data Fig. 17:** Genomic mapping of clean single-copy T-DNA insertions by inverse PCR

#### SUPPLEMENTARY NOTES

**Supporting guide for T-DNA MoClo toolkit**

#### SUPPLEMENTARY REFERENCES

#### SUPPLEMENTARY TABLES

**Extended Data Table 1:** Transformation counts

**Extended Data Table 2:** iPCR sequencing results

**Supplementary Table 1:** Plasmids used in this study

**Supplementary Table 2:** T-DNA plasmid toolkit

**Supplementary Table 3:** Arabidopsis promoter design

**Supplementary Table 4:** Arabidopsis terminator design

**Supplementary Table 5:** Oligonucleotides used in this study

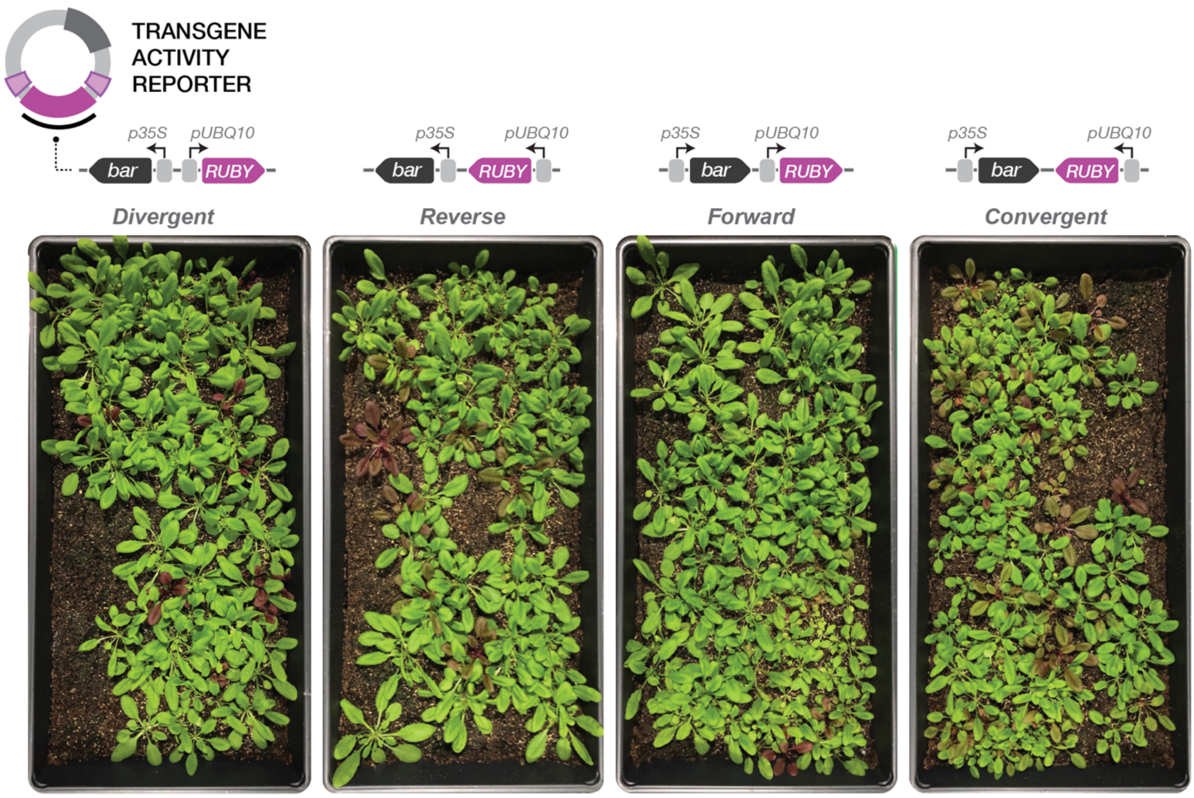


**Extended Data Fig. 1: Arabidopsis transformants from the four T-DNA arrangements in Fig. 1c,d.** Image taken of transformants at 28 DPG following selection of 10,000 seeds with a 3-week BASTA treatment.

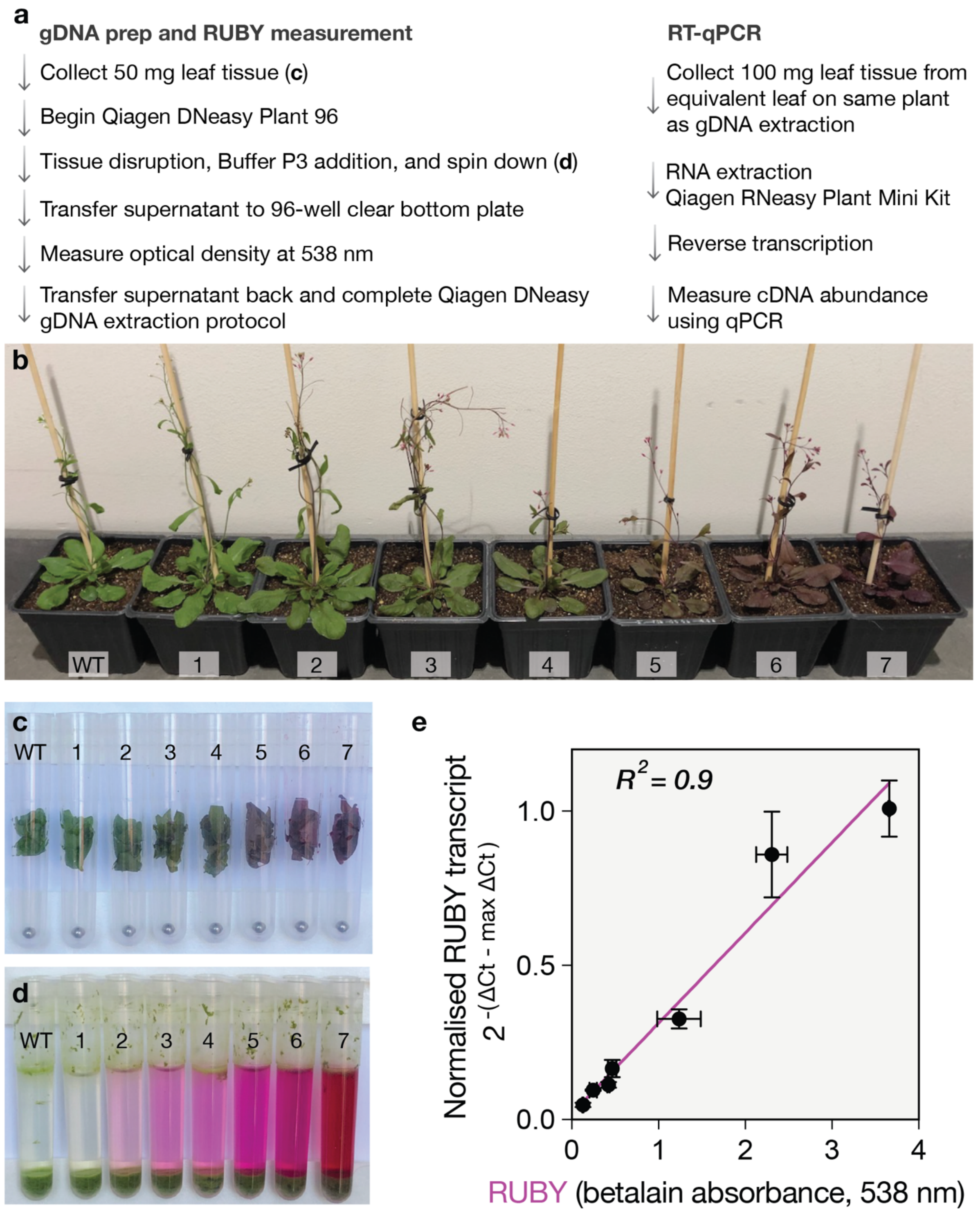


**Extended Data Fig. 2: Extraction of betalain using the Qiagen DNeasy Plant 96 kit.** **a**, Parallel workflows for gRNA and betalain extraction using QIAGEN DNeasy Plant 96 kit and RNA extraction for RT-qPCR, taking leaf tissue at 28 DPG. **b**, a panel of RUBY expressing plants (1-7) and WT control used in the betalain-RT-qPCR comparison, showing a range of expression levels from unobservable to deep red. Photos taken ~40 DPG. **c**, 50 mg of leaf tissue taken from plants in **b**. **d**, Tissue samples after the first disruption step, addition of Buffer P3, and spin down in the QIAGEN DNeasy Plant 96 protocol. **e**, Relative transcript levels compared to optical density at 538 nm of disrupted tissue. Transcript levels were normalised to the maximum measurement (line 7). A straight-line curve was fitted using GraphPad Prism linear regression fit, $R^2 = 0.9$.

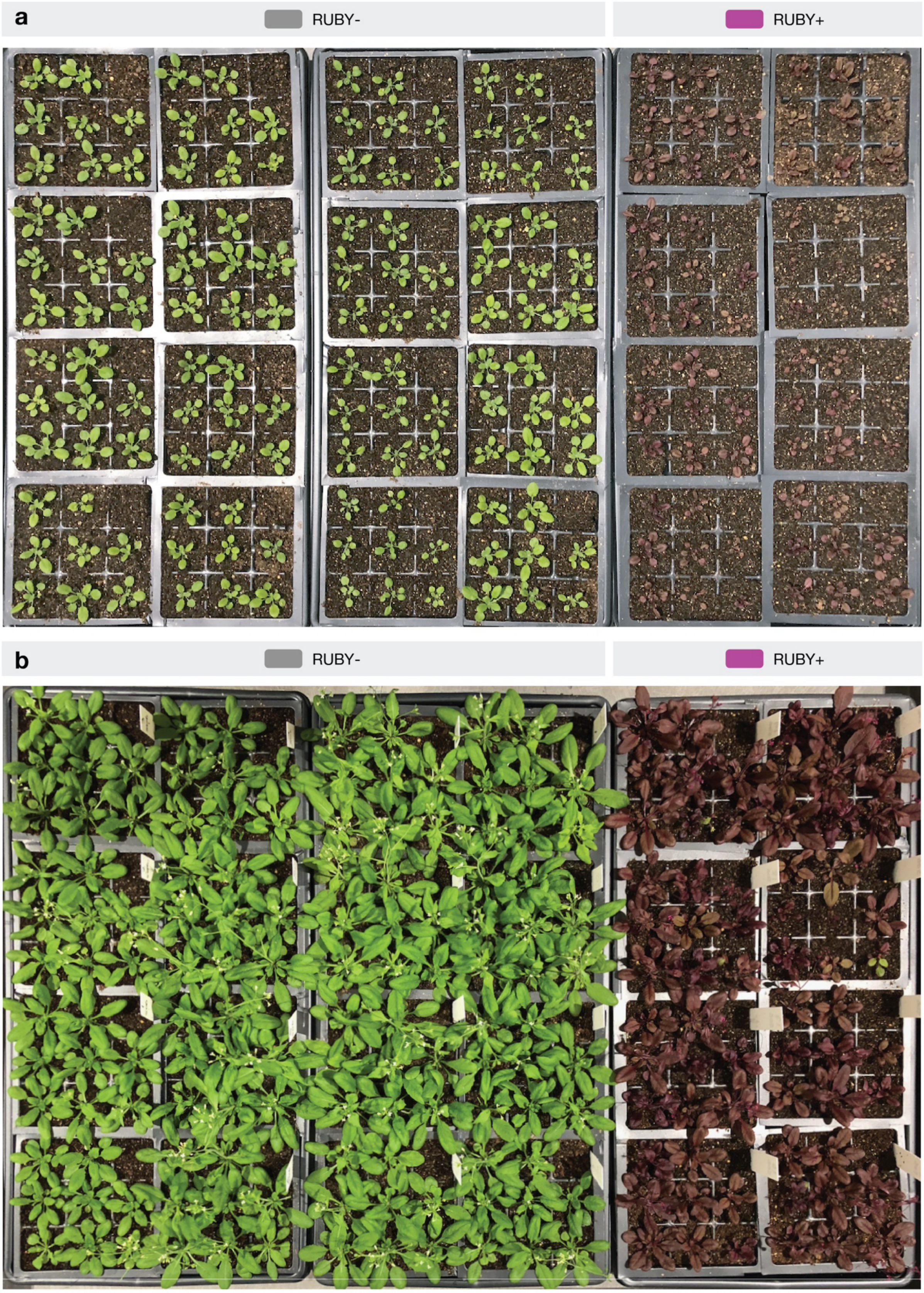


**Extended Data Fig. 3: Transplanted green and red Arabidopsis transformants from Fig. 1e**. **a**, Arabidopsis transformants at 21 DPG, showing the highly binary and uniform nature of red pigmentation. **b**, Arabidopsis transformants at ~35 DPG, showing continuation of binary pigmentation, and some late silencing in the red plants.

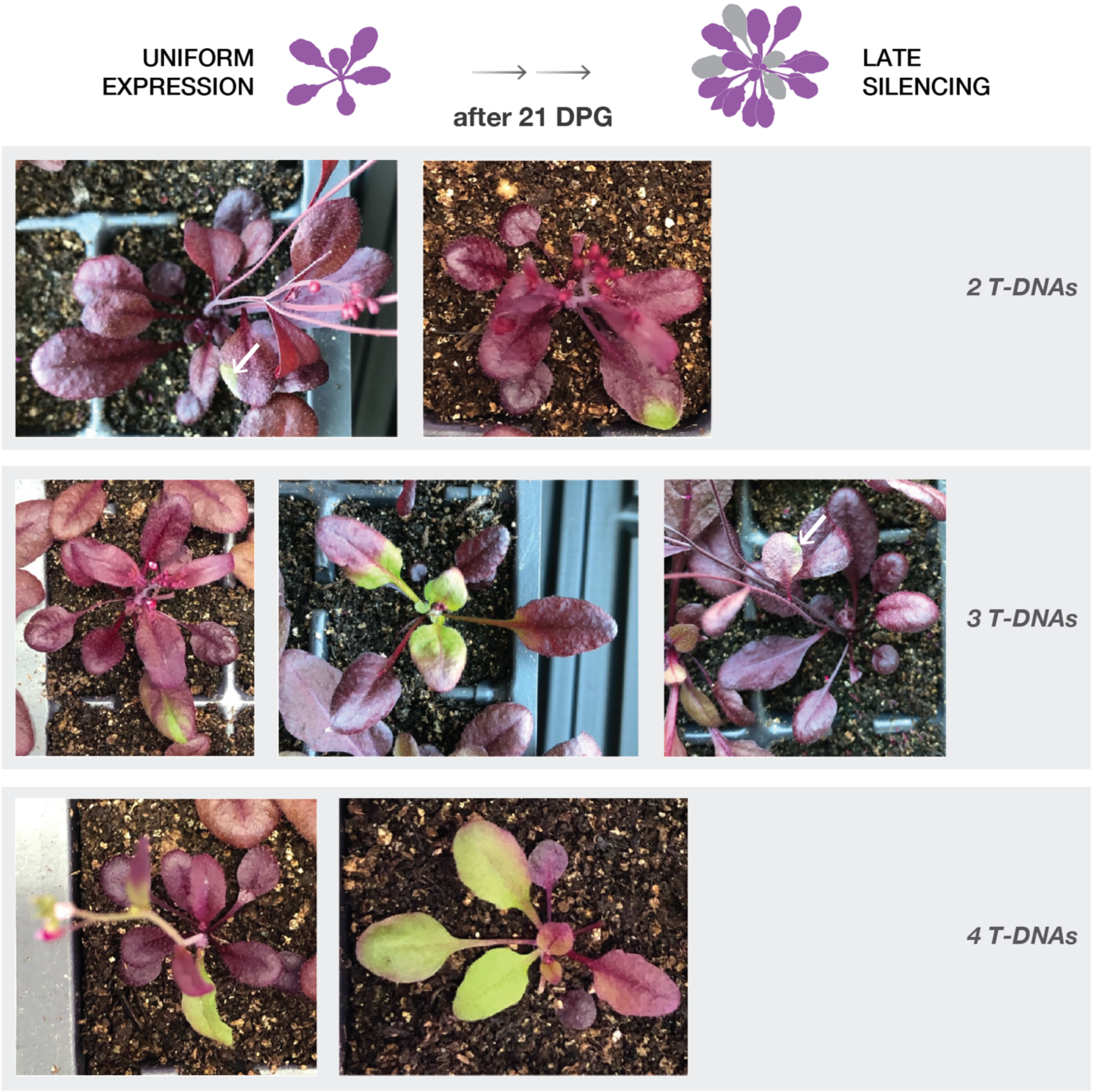


**Extended Data Fig. 4: Late silencing in Arabidopsis transformants from Fig 1e.** Visible patches of green are seen in some plants containing 2 T-DNA, and all plants containing 3 or 4 T-DNAs. White arrow used to highlight green patch in top left and mid right plants.

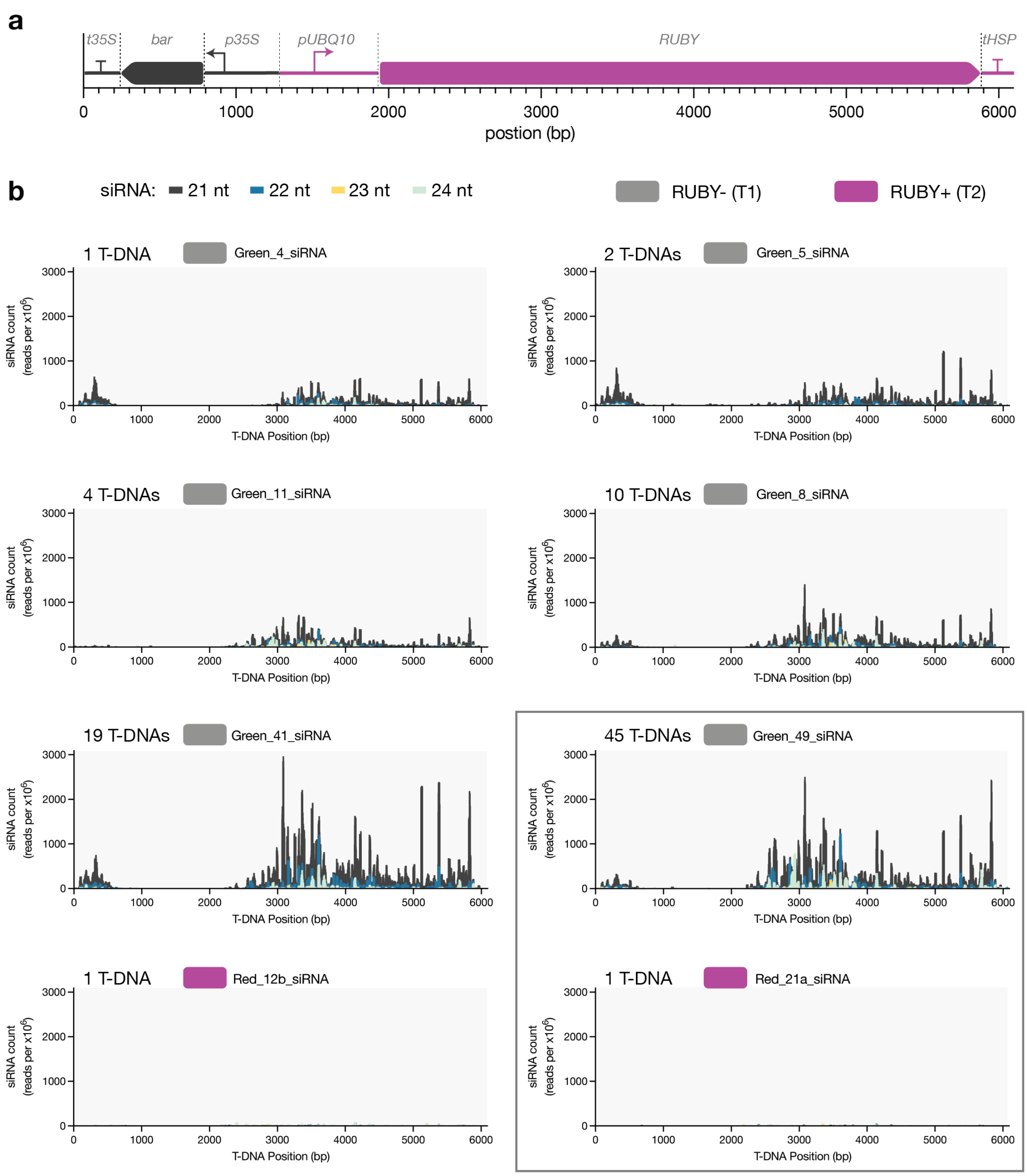


**Extended Data Fig. 5: Small RNA profiles of 6 green (RUBY-) and 2 red (RUBY+) plants. a**, Coordinates for the *p35S:bar:t35S* and *pUBQ10:RUBY:tHSP* cassettes in divergent arrangement. Small RNA profiles mapped across the T-DNA according to the coordinates in **a**, showing 21–24 nt small RNAs. Small RNAs were sequenced from green T1 plants and red T2 plants at 21 DPG. Transformants were selected to represent a range of T-DNA copy numbers, as determined by qPCR. The T-DNA copy number for each line is indicated in the upper-left corner of the corresponding plot. Box represents the data shown in Fig. 1g.

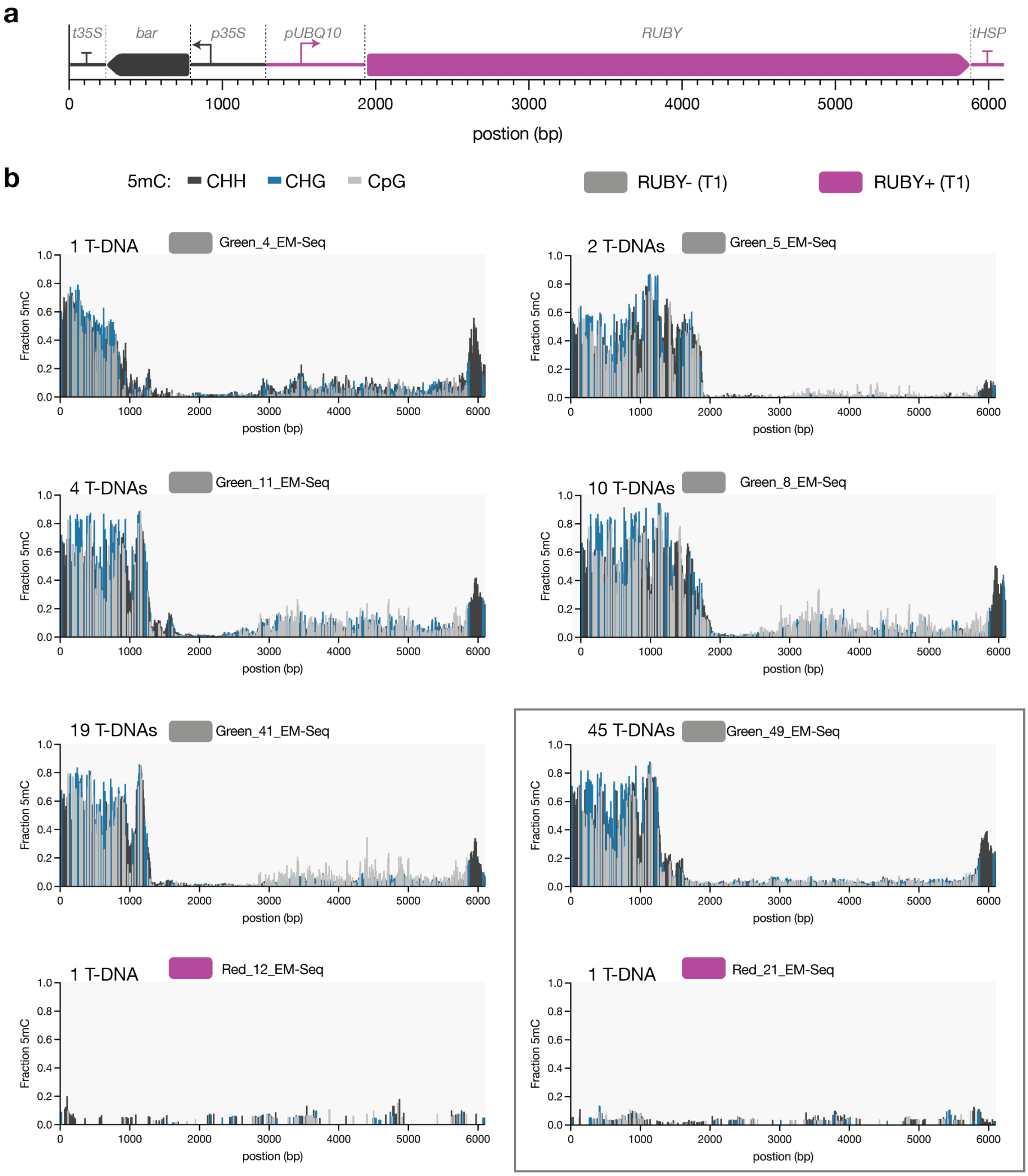


**Extended Data Fig. 6: 5mC profiles of 6 green (RUBY-) and 2 red (RUBY+) plants. a**, Coordinates for the *p35S:bar:t35S* and *pUBQ10:RUBY:tHSP* cassettes in divergent arrangement. 5mC profiles mapped across the T-DNA according to the coordinates in **a**, in all contexts (CG, CHG, and CHH). Whole genome EM-Seq was performed on green T1 plants and red T1 plants at 21 DPG. Transformants were selected to represent a range of T-DNA copy numbers, as determined by qPCR. The T-DNA copy number for each line is indicated in the upper-left corner of the corresponding plot. Box represents the data shown in Fig. 1h.

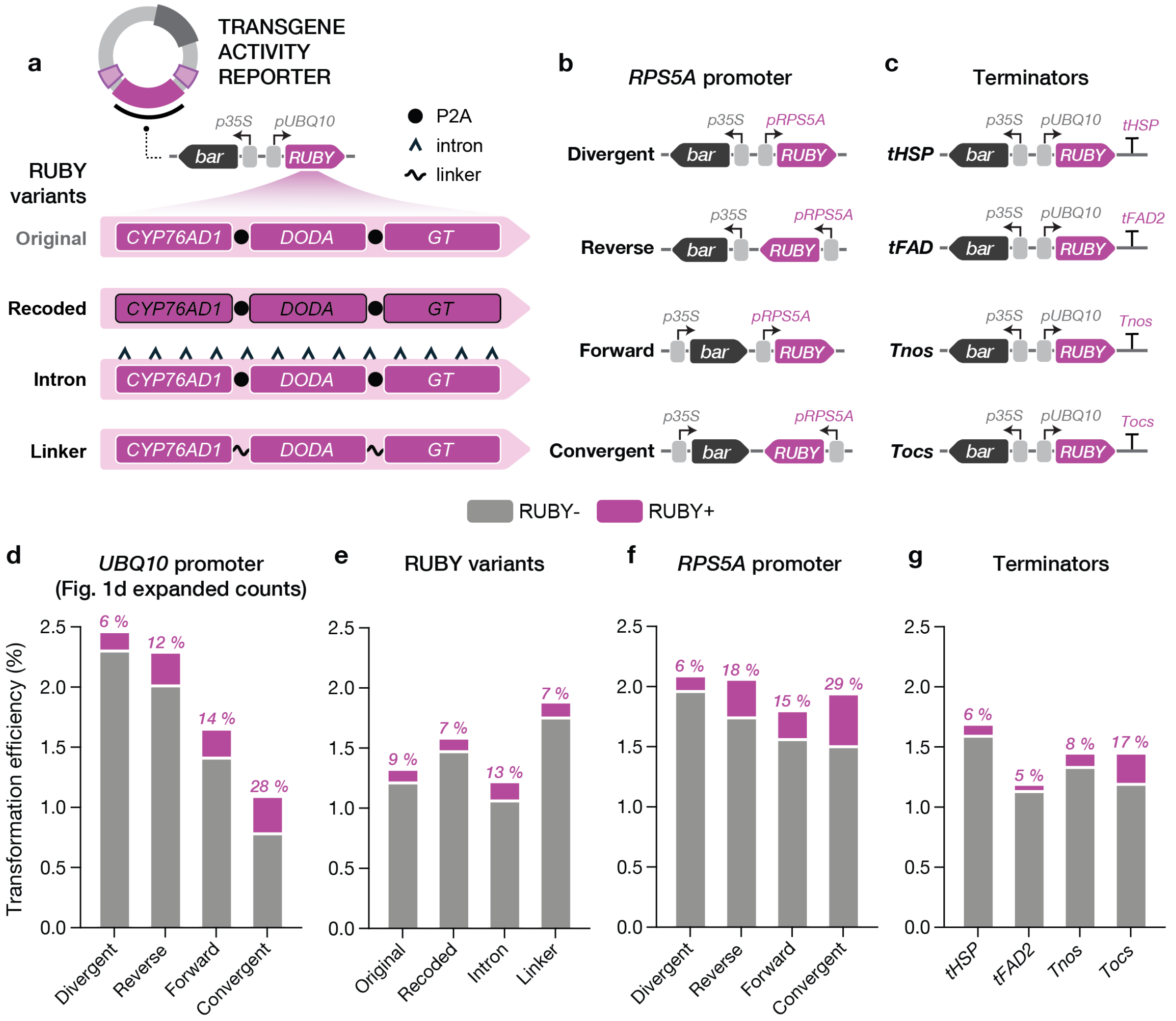


**Extended Data Fig. 7: High prevalence of *RUBY* silencing cannot be explained by major features of the transcriptional unit. a,** Schematic of RUBY coding and protein sequence variants: the original *RUBY* design, showing the three genes, cytochrome P450 *CYP76AD1*, L-DOPA 4,5-dioxygenase (*DODA*), and glucosyl-transferase (*GT*), co-expressed from a polycistronic transcript using the self-cleaving P2A peptide (Original); an Arabidopsis codon-optimised version (Recoded); a construct containing 12 introns (Intron); and a variant in which the P2A self-cleaving peptides were replaced with an equivalent-length flexible (GGGGS) linker (Linker). These designs tested potential causes of silencing, including rare codons in the original non-Arabidopsis-optimised sequence[1], the absence of introns over the long ~4 kb coding sequence[2, 3], and translational effects associated with self-cleaving 2A peptides that function via a ribosome-stalling "StopGo" mechanism[4]. **b,** The four T-DNA configurations shown in Fig. 1c,d were re-engineered by replacing the strong *UBQ10* promoter with the weaker *RPS5A* promoter to test the relationship between expression level and silencing. **c,** The Arabidopsis *HSP18.2* terminator (*tHSP*) in the original design was substituted for the Arabidopsis *FAD2* (*tFAD2*) and *Agrobacterium nos* (*Tnos*) and *ocs* (*Tocs*) terminators. **d-g,** Transformation efficiency of various genetic designs from Fig. 1c,d and panels a-c, showing the proportion of red and green plants. **d**, Expanded transformation counts from the four arrangements in Fig. 1c,d, using the strong Arabidopsis *UBQ10* promoter. **e,** No major differences were seen between RUBY variants and the equivalent divergent arrangement in panel d. **f**, No major differences were seen between terminators and the equivalent divergent arrangement in panel d. **g**, No major differences were seen between the *RPS5A* promoter arrangements and their equivalent arrangement in panel d. A variable number of seeds were assessed per condition (See **Extended Data Table 1** For a list of transformation counts). Percentage of RUBY+ plants shown in pink.

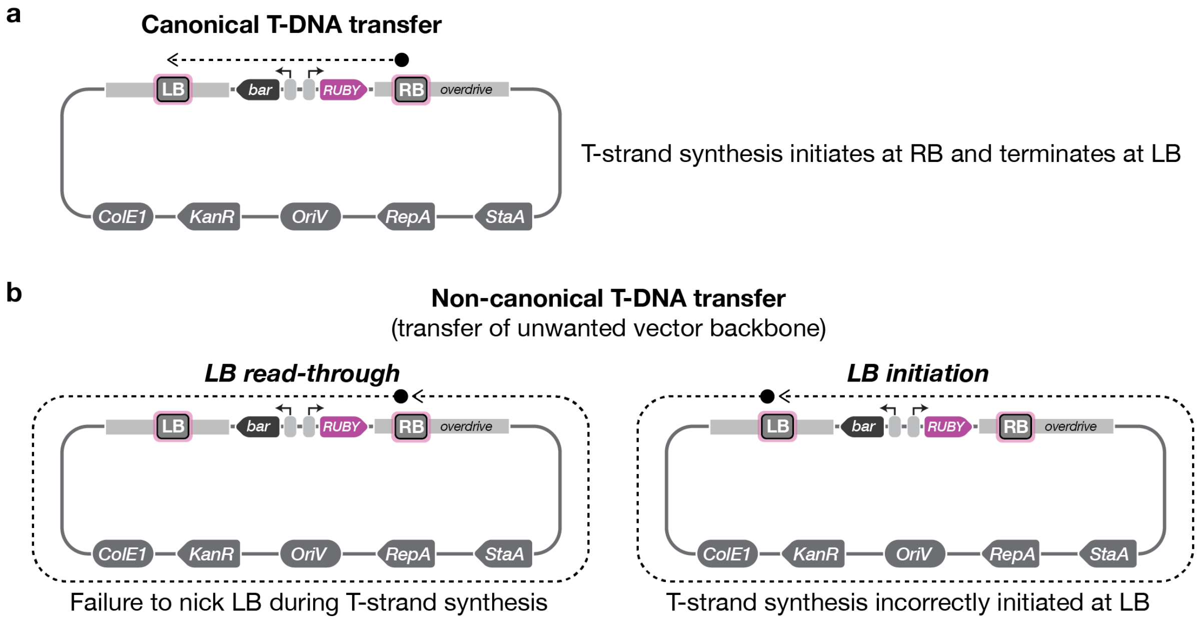


**Extended Data Fig. 8: Mechanisms of T-DNA vector backbone transfer.** **a**, Canonical T-strand synthesis without the aberrant transfer of vector backbone sequence. The left (LB) and right (RB) borders are 25-bp imperfect direct repeats recognised by the VirD1/VirD2 complex. VirD2 binds and nicks the RB, becoming covalently attached to the 5′ end of the single-stranded T-DNA. DNA synthesis then repairs the nicked strand across the T-DNA region, displacing the VirD2-bound DNA (T-strand) in the process. A second nick at the LB releases the T-strand for transfer into the plant cell. **b**, Two mechanisms of vector backbone transfer during non-canonical T-strand synthesis. Failure of VirD2 to nick the LB results in LB read-through, whereby strand displacement continues beyond the LB and incorporates adjacent vector backbone sequences into the transferred DNA (left). Alternatively, the VirD1/VirD2 complex can initiate T-strand synthesis at the LB, resulting in direct transfer of vector backbone DNA (right). Black circles indicate sites of VirD1/VirD2 binding and T-strand initiation, and dotted lines represent the displaced T-strand. Arrow endpoints are illustrative for the LB initiated T-strand synthesis and do not necessarily indicate the termination site, which may also occur at the RB.

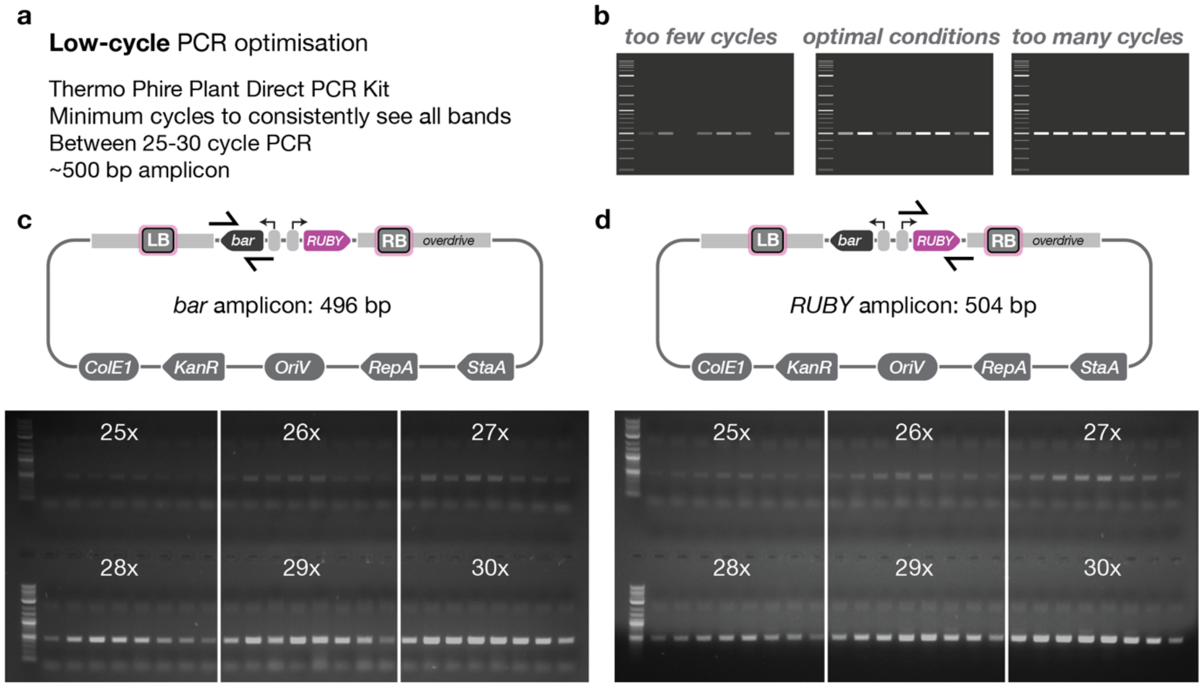


**Extended Data Fig. 9: Optimisation of a low-cycle PCR protocol for detecting presence of T-DNA events in Arabidopsis. a,** Determination of optimal PCR cycling conditions to identify a cycle number (25–30 cycles) that reliably amplifies the expected ~500 bp product while avoiding over-amplification, which increases the likelihood of detecting contaminating plasmid DNA from *Agrobacterium*. **b,** Representative agarose gels illustrating the optimisation process: *too few cycles,* where not all expected bands are visible; *too many cycles*, where bands are strongly amplified and background amplification increases; and *optimal conditions*, where all expected bands are detectable but not over amplified. **c**, PCR amplification of the BAR selection marker in low-copy red plants over variable cycles, creating a 496 bp product. **d**, PCR amplification of the RUBY reporter in low-copy red plants over variable cycles, creating a 504 bp product. ~0.5 cm of tissue was collected from Arabidopsis transformants at 21 DPG and prepared for genotyping using the Phire Direct PCR kit (see **Methods**). DNA size marker: NEB 1 kb Plus ladder.

Across these tests, **28 cycles** provided the optimal balance, consistently producing visible bands while maintaining some reactions near the detection threshold. This condition was therefore selected for the final low-cycle PCR protocol.

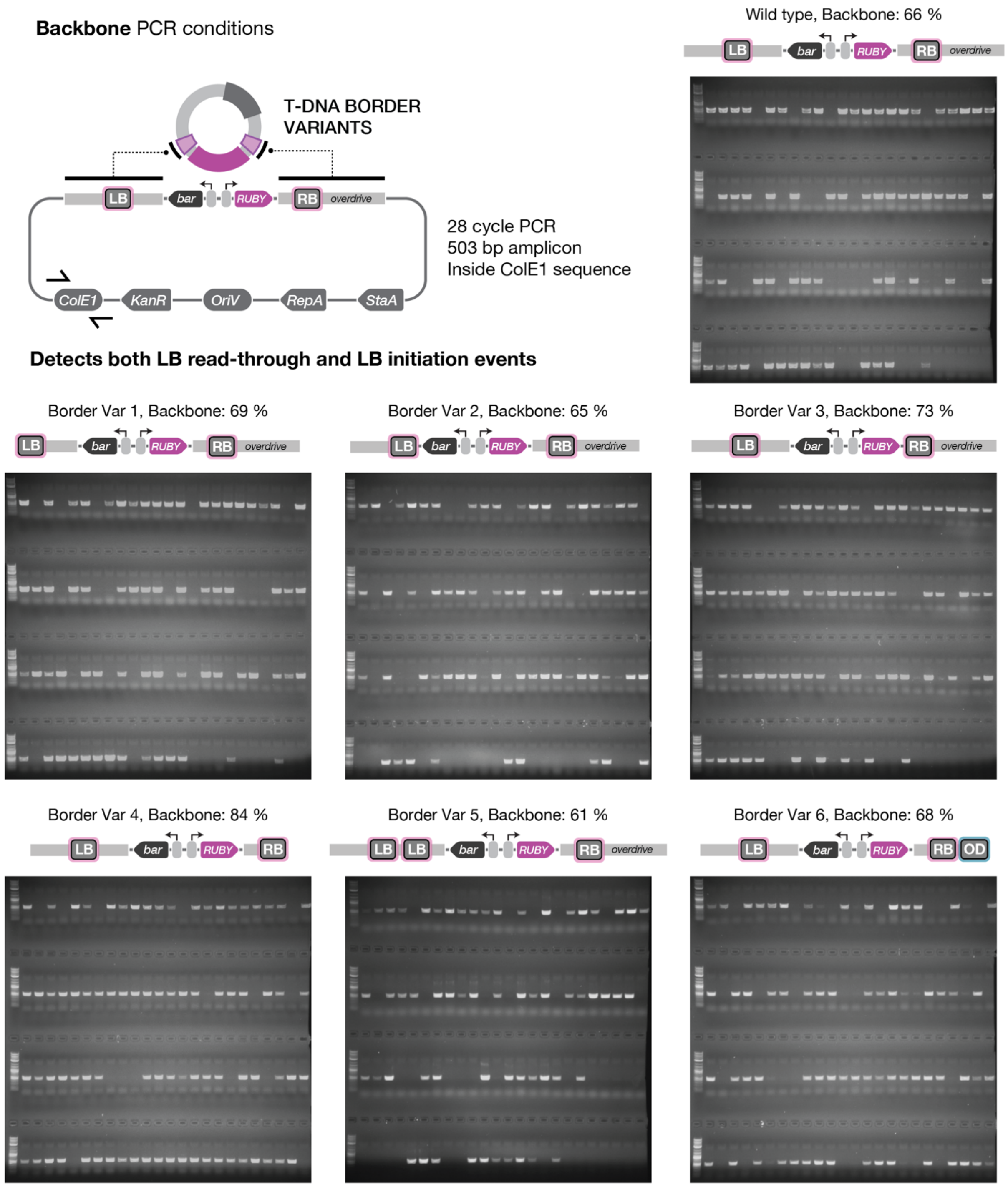


**Extended Data Fig. 10: Amplification of the ColE1 replicon to detect the integration of all backbone events in Arabidopsis transformants.** PCR amplification of the ColE1 backbone replicon was performed using the optimised low-cycle protocol (28 cycles), generating an expected 503 bp product. Leaf tissue was collected at 21 DPG from 96 randomly selected transformants for each of the seven T-DNA border sequence variants, without selection bias for RUBY+ or RUBY− phenotypes. PCR products were resolved by agarose gel electrophoresis, and individuals were scored for the presence or absence of a 503 bp band, indicating integration of vector backbone sequences. These binary scores were used to quantify total backbone integration frequencies, as summarised in Fig. 2c ("backbone"). DNA size marker: NEB 1 kb Plus ladder.

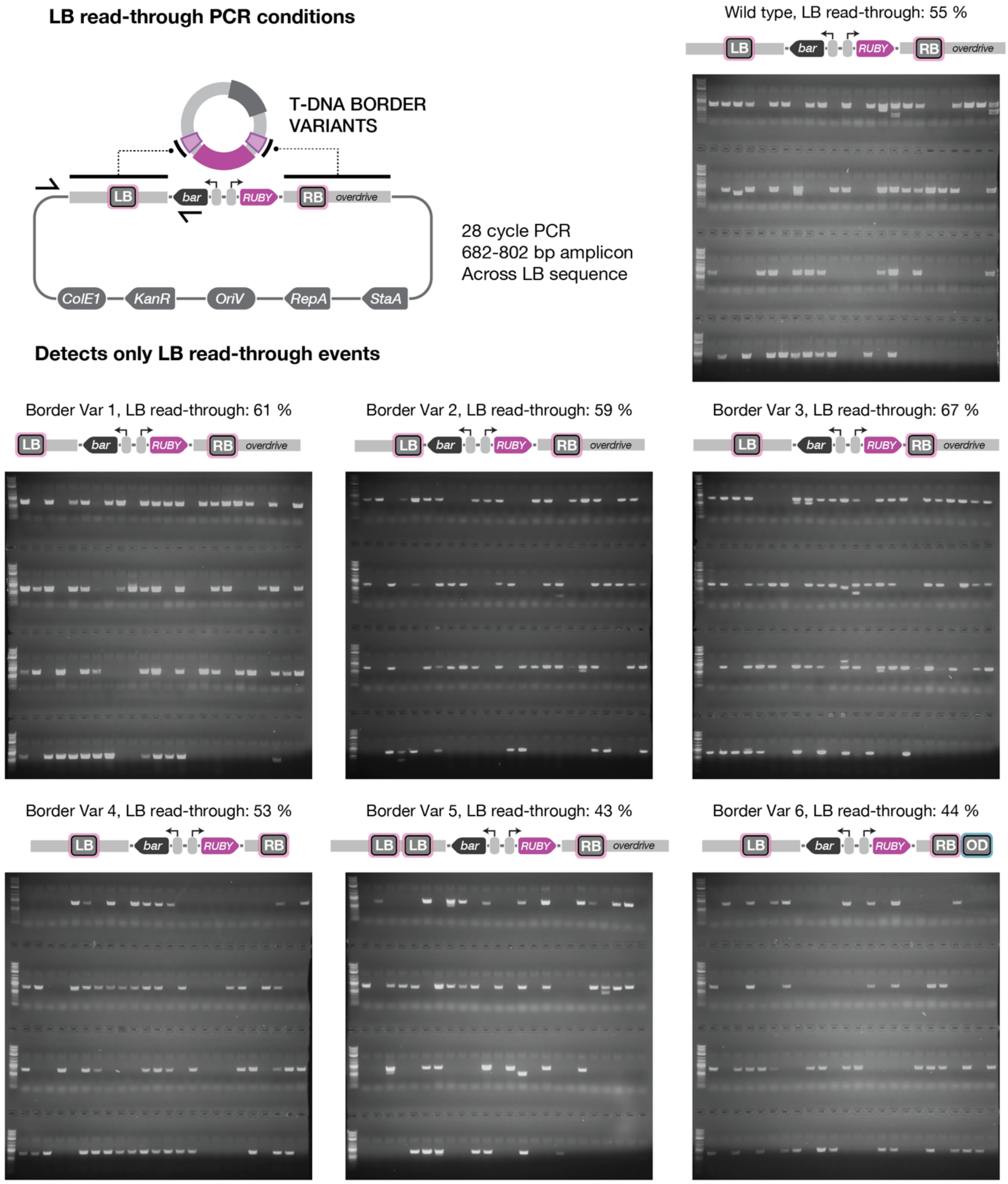


**Extended Data Fig. 11: Amplification across left border sequence (LB) to detect the integration of LB readthrough events in Arabidopsis transformants.** PCR amplification across the LB sequence was performed using the optimised low-cycle protocol (28 cycles), generating an expected product between 682-802 bp, depending on the design at the LB. Leaf tissue was sampled from 96 randomly selected transformants representing the seven T-DNA border sequence variants at 21 DPG. PCR products were resolved by agarose gel electrophoresis, and individuals were scored for the presence or absence of a band in the 682-802 bp range, indicating integration of vector backbone sequences attributed to LB read-through. These binary scores were used to quantify LB read-through frequencies, as summarised in Fig. 2c (“LB read-through”). DNA size marker: NEB 1 kb Plus ladder.

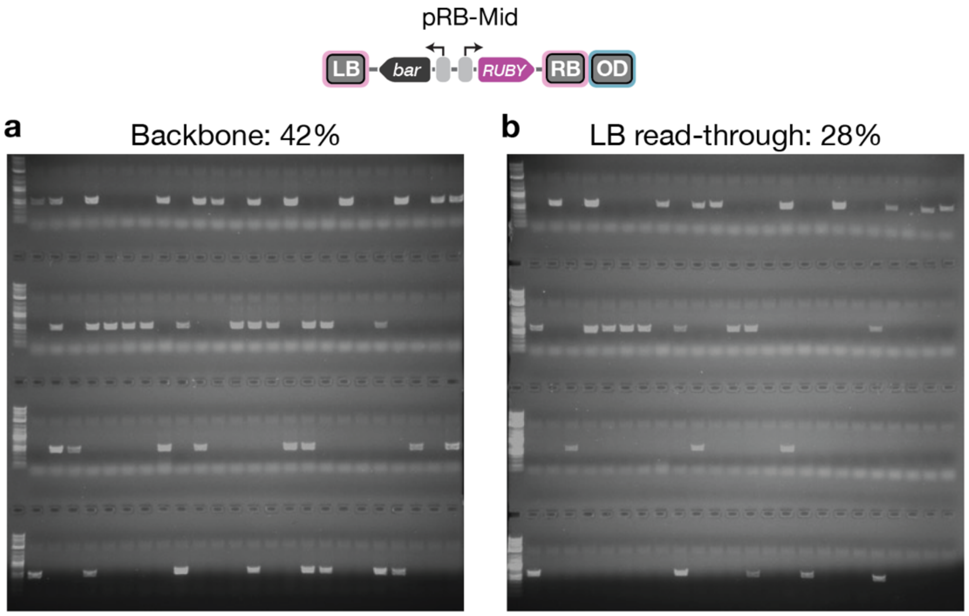


**Extended Data Fig. 12: Detection of backbone transfer in Arabidopsis transformants using the pRB-Mid vector. a,** PCR amplification of the ColE1 backbone replicon was performed using the optimised low-cycle protocol (28 cycles), generating an expected 503 bp product. **b**, PCR amplification across the LB sequence was performed using the optimised low-cycle protocol (28 cycles), generating an expected 629 bp product. Leaf tissue was collected at 21 DPG from 96 randomly selected transformants, without selection bias for RUBY+ or RUBY− phenotypes. PCR products were resolved by agarose gel electrophoresis, and individuals were scored for the presence or absence of a band with the indicated size. DNA size marker: NEB 1 kb Plus ladder.

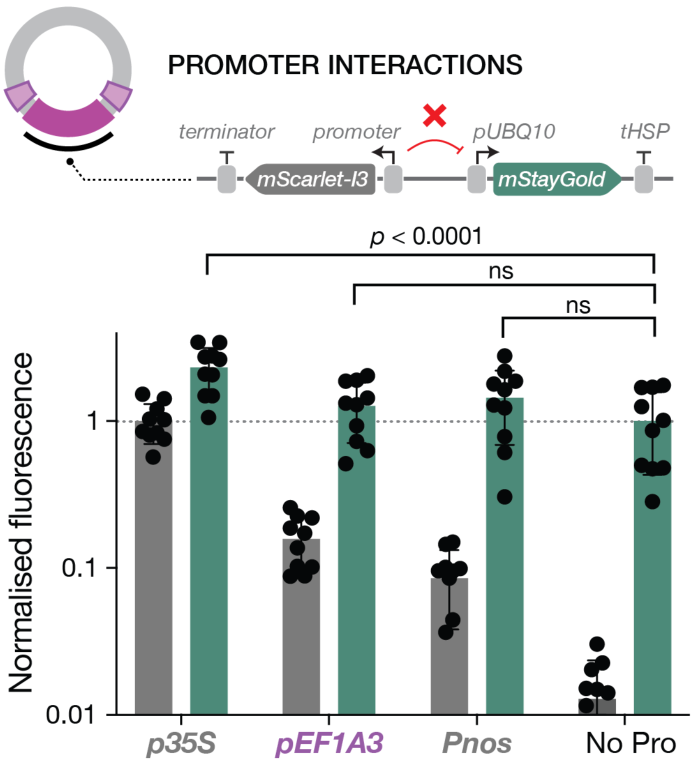


**Extended Data Fig. 13: Evaluating the effect of the *EF1A3* promoter on strong constitutive transgene expression in the divergent arrangement.** The *EF1A3* promoter driving mScarlet-I3 (*pEF1A3:mScarlet-I3:tUBQ11*) was cloned in a divergent orientation to the *UBQ10* promoter driving mStayGold (*pUBQ10:mStayGold:tHSP*). Its effect on neighbouring gene expression was evaluated alongside *35S* (*p35S:mStayGold:t35S*) and *nos* (*Pnos:mStayGold:Tnos*) promoter controls, as well as a no-promoter control lacking divergent mScarlet-I3 expression. No significant change in mStayGold expression was observed with *pEF1A3*, indicating that the apparent absence of enhancer activity is unlikely to result from a repressive effect on neighbouring promoters. Although statistically insignificant, a small increase in mStayGold expression was observed for the *pEF1A3* condition (1.26-fold), which may reflect favourable biophysical effects associated with divergent gene expression[5]. Statistical significance in mStayGold expression relative to the no-promoter control was assessed using two-way ANOVA with multiple comparisons in GraphPad Prism, and significance levels are shown.

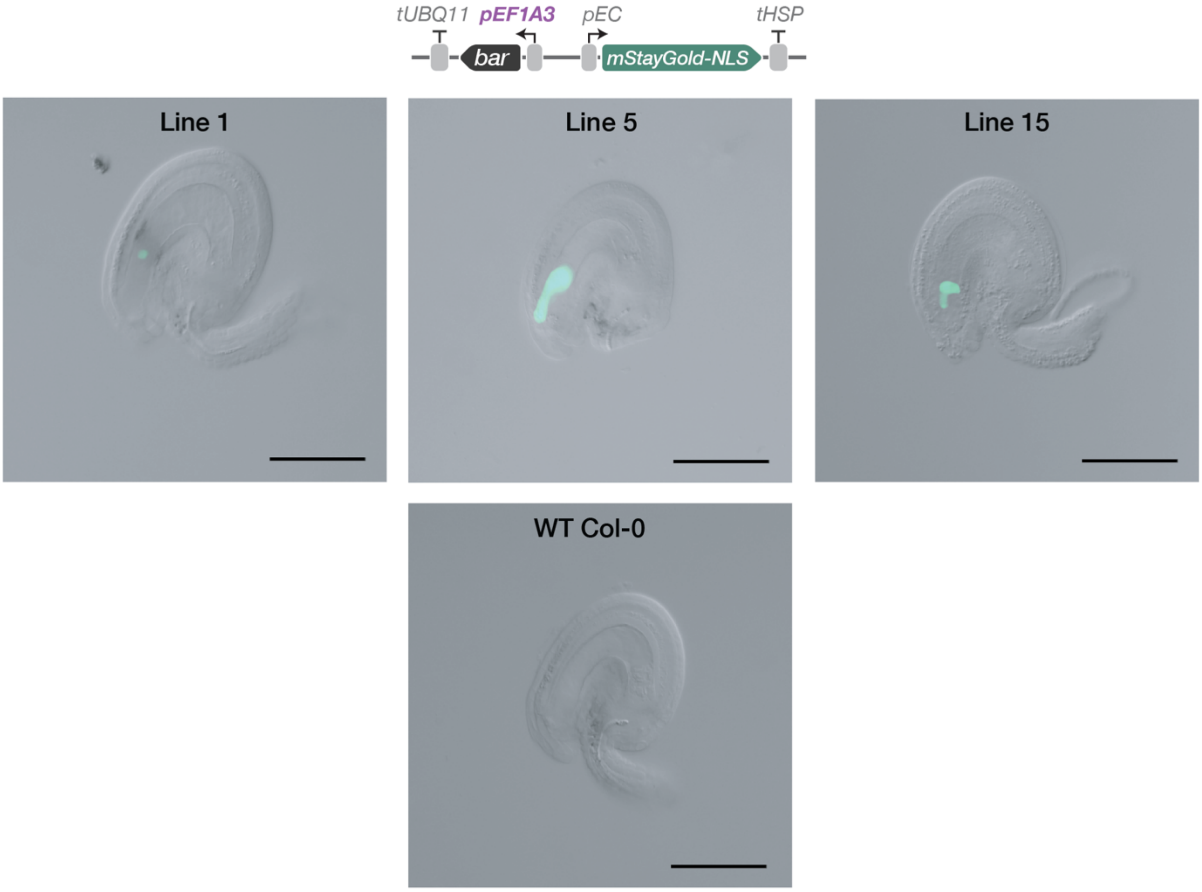


**Extended Data Fig. 14: Egg cell-specific expression of nuclear-localised GFP using the *pEF1A3*-based selection marker system.** Representative images of unfertilised ovules from four independent transgenic lines carrying the indicated construct in the pRB-Min T-DNA vector. Images show overlays of DIC and GFP channel. Scale bars are 50 µM.

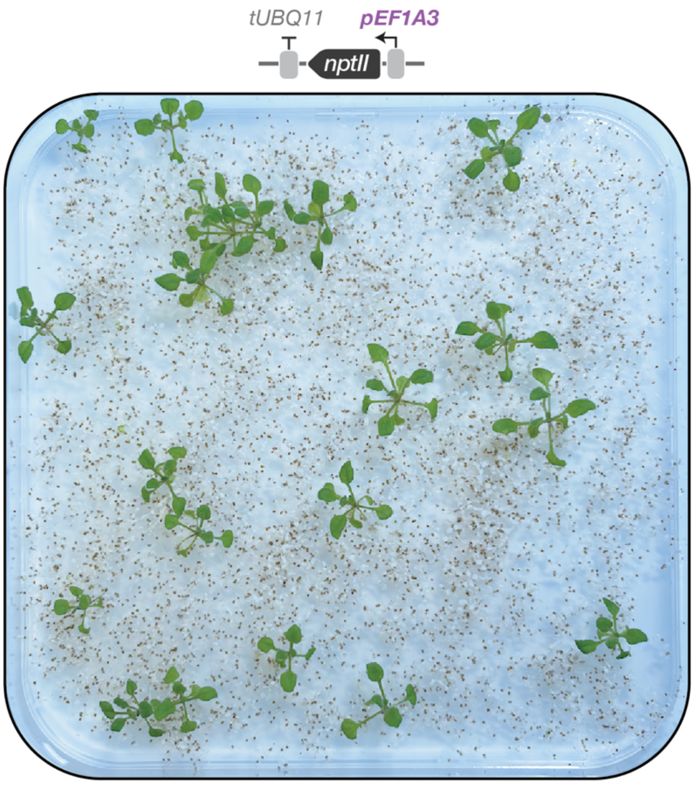


**Extended Data Fig. 15: Robust growth of antibiotic-resistant transformants using the *pEF1A3*-based selection cassette.** Selection of 5,000 seeds transformed with *pEF1A3:nptII:tUBQ11* on 0.5x MS medium with 50 mg/L kanamycin. Image taken 14 DPG.

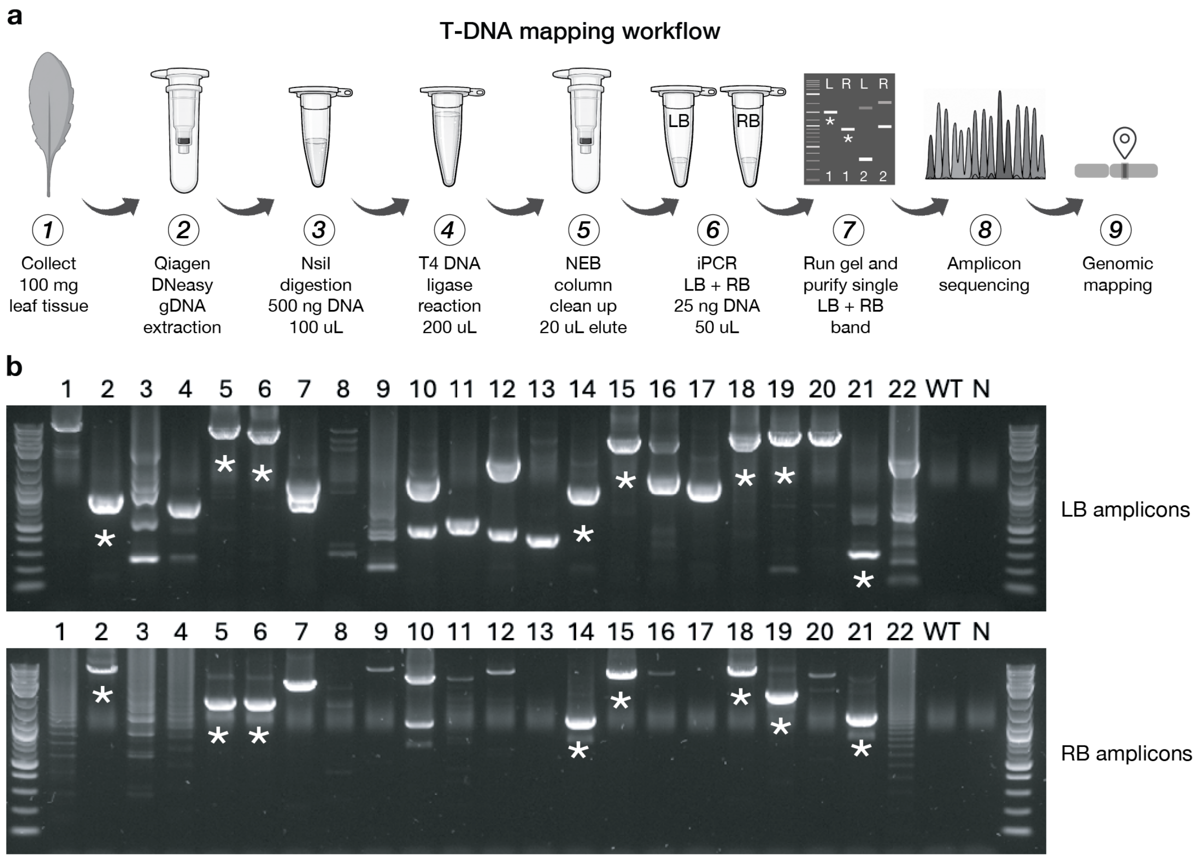


**Extended Data Fig. 16: Inverse PCR (iPCR) workflow and analysis of T-DNA insertion events in randomly selected Arabidopsis transformants.** **a**, Schematic overview of the iPCR workflow used for mapping T-DNA insertion sites. **b**, Agarose gel analysis of 5 µL iPCR products from 22 randomly selected T1 transformants, amplifying junctions at the LB (top) and RB (bottom). Lines exhibiting a single amplification band at both borders were selected for sequencing, using the remaining PCR volume, and are indicated by an asterisk (*). WT, untransformed Col-0 control; N, no-template control. DNA size marker: NEB 1 kb Plus ladder.

Amplicons from eight lines were sequenced to resolve T-DNA insertion structure. Six lines (2, 5, 6, 15, 18, and 21) mapped precisely at both the LB and RB to the same genomic locus without detectable vector backbone integration, consistent with clean single-copy insertion events. All six lines demonstrated consistent RUBY expression. One line (14) mapped to two distinct loci on chromosome 5 separated by 12.2 Mbp (LB, Chr5:1397972..1398927; RB, Chr5:13576233..13577280), consistent with either a double T-DNA integration event or a large chromosomal rearrangement/deletion. Another line (19) yielded a correctly mapped RB amplicon (Chr1:4946719..4948952), whereas the LB amplicon aligned to vector backbone sequence, indicating incomplete border processing and backbone integration. The iPCR workflow therefore enables efficient identification of single-copy insertion events, detection of vector backbone integration, and precise mapping of genomic insertion sites through a simple and scalable workflow. Using the new T1 vector series, 27% of transformants possessed all desired characteristics: clean, single-copy, fully mappable insertions with consistent transgene expression.

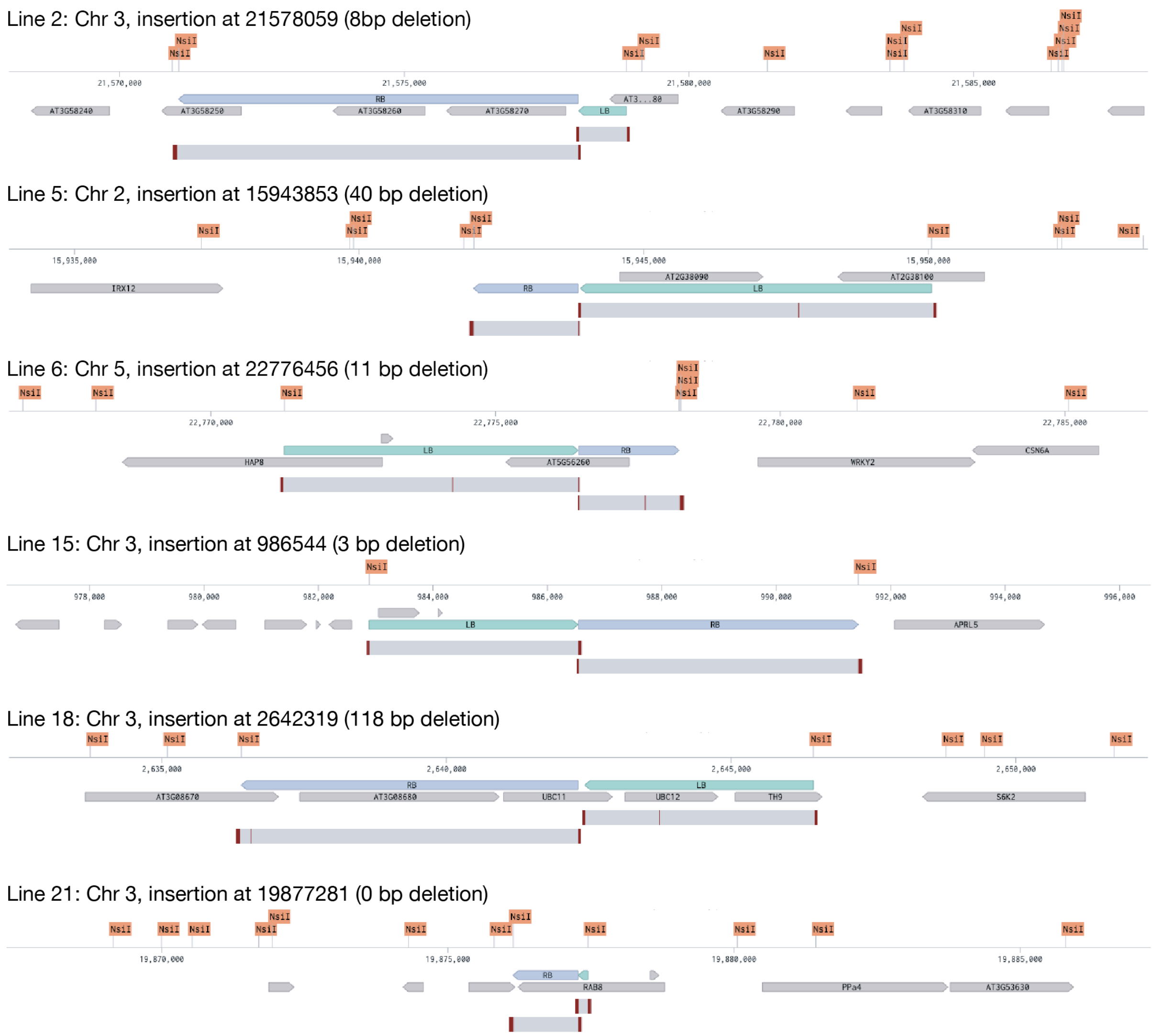


**Extended Data Fig. 17: Genomic mapping of clean single-copy T-DNA insertions by inverse PCR.** Genomic regions surrounding the six successfully mapped insertion events are shown, centred on the insertion site ±10 kb using TAIR10 reference in Benchling. The indicated deletion corresponds to the genomic region absent between the LB and RB amplicons and is presumed to have been lost during T-DNA integration. LB (green) and RB (blue) iPCR amplicons are annotated alongside *Arabidopsis thaliana* gene models (grey). All NsiI restriction sites are indicated in orange. The distal end of each amplicon terminates precisely at an NsiI site for all sequence alignments, consistent with successful restriction digestion and circularisation during iPCR sample preparation. Sequence alignments for LB and RB amplicons are shown below each genomic map. Grey indicates perfect sequence identity, whereas red indicates mismatches. Mismatches at the ends of reads correspond to flanking T-DNA-derived sequences, including the barcode and 100 bp spacer regions. No additional sequences, including vector backbone DNA, were detected in any mapped insertion for the six lines shown.

### Supporting guide for T-DNA MoClo toolkit

This guide provides an overview of the MoClo standard and serves as an operational manual for the T-DNA toolkit. It is designed to support both new and experienced users. The first section outlines the T-DNA toolkit plasmid architectures, introduces a simplified MoClo nomenclature, and describes cloning strategies for assembling multigene constructs. The second section details the essential cloning operations required to build single- and multigene T-DNA constructs, covering the needs of most users. The final section presents advanced cloning strategies available within the toolkit, offering additional flexibility for more complex applications. The toolkit is based on the original MoClo design of Weber et al.[6] and is fully forward- and backward-compatible.

All cloning in this work was performed using TOP10 *E. coli* (Thermo Scientific). Competent cells were prepared in-house using the KCM/TSS protocol (see **Methods**). We recommend using this strain because the toolkit has been extensively validated in TOP10 cells and produces strong GFP and RFP dropout signals for screening successful assemblies, even in small colonies. Fluorescent screening can be performed at low cost using a blue LED lamp and orange-lensed blue-light-blocking glasses. Once colonies are sufficiently large, GFP and RFP fluorescence can also be distinguished by eye.


**T-DNA toolkit and MoClo Basics**
T-DNA toolkit plasmid architectures
MoClo formatting and a simplified nomenclature
Combined Level 1 and Level 2 nomenclature
Alternative cloning approaches for Level 2 multigene constructs

**Essential MoClo cloning operations**
Assembling promoter + 5' UTR (Pro) sequences into the part entry plasmid
Assembling coding sequences (CDS) into the part entry plasmid
Assembling 3' UTR + terminator (Ter) sequences into the part entry plasmid
Assembling a Level 1 transcriptional unit in the forward direction (Forward TU)
Assembling a Level 1 transcriptional unit in the reverse direction (Reverse TU)
Assembling a Level 1 transcriptional unit (TU) into a T-DNA vector
Example Level 2 multigene assembly into a T-DNA vector

**Advanced MoClo cloning operations**
Moving assembled constructs between T-DNA vectors
Assembling multigene constructs for future BsaI transcriptional unit expansion
Assembling a transcriptional unit into a BsaI expandable T-DNA vector
Converting a BsaI expandable T-DNA vector for BpiI multigene expansion
Assembling a multigene construct into a BpiI expandable T-DNA vector
Assembling a multigene construct into the expansion module plasmid

#### T-DNA toolkit and MoClo basics

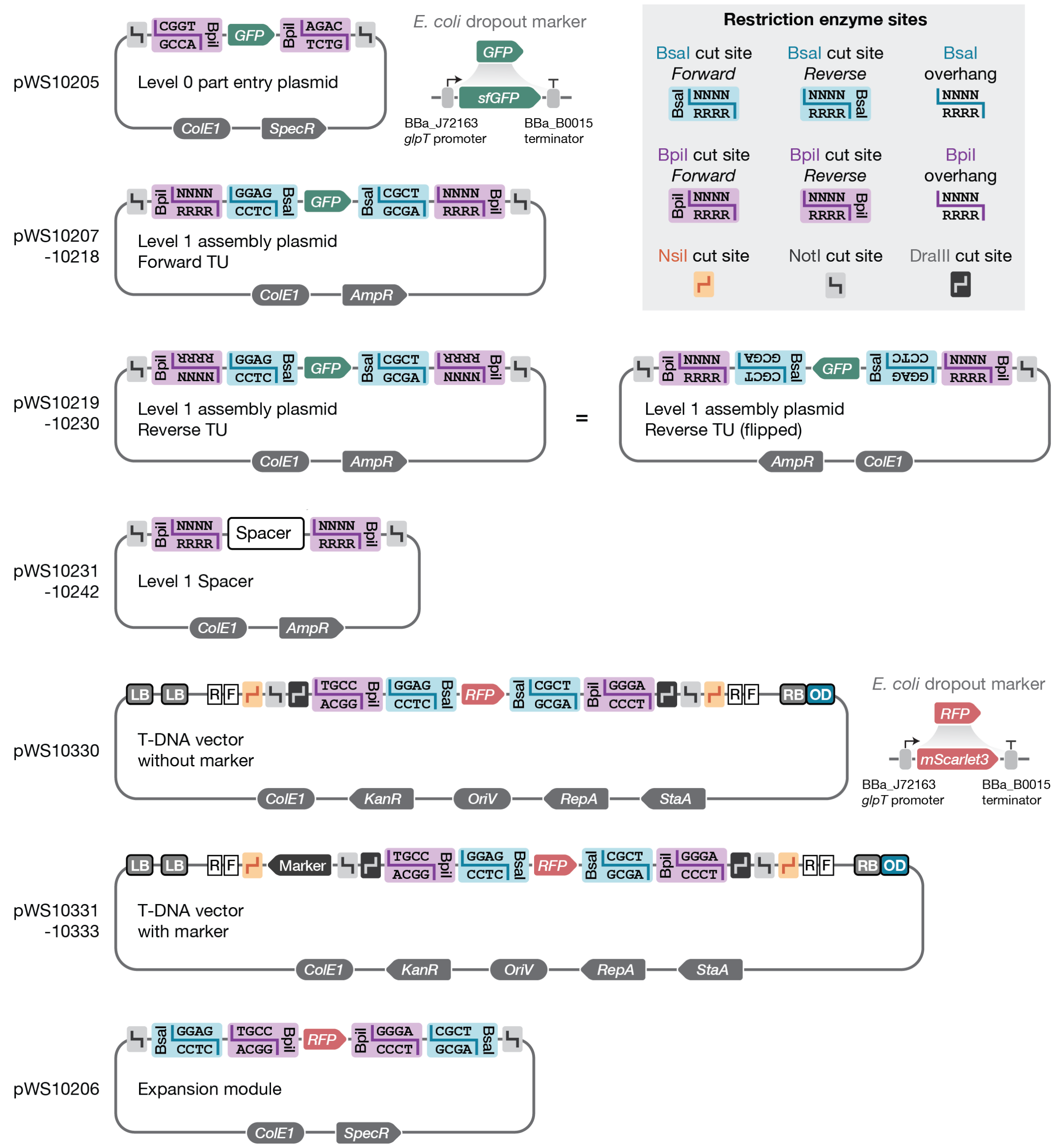


**T-DNA toolkit plasmid architectures.** Plasmid backbones are shown with key features highlighted, including restriction sites and cloning dropout fluorescence markers for screening assembled constructs in *E. coli*. All plasmids contain flanking NotI sites surrounding the Golden Gate Assembly region, enabling rapid validation of assemblies by diagnostic digest. Consistent with the original MoClo system, DraIII sites are included in the T-DNA vector backbone to facilitate the transfer of assembled constructs between T-DNA vectors with different plant selectable markers. These sites also facilitate exchange of the dropout cassette with the original MoClo cloning sequences and alternative cloning systems. All plasmids in the T-DNA toolkit have been cured of all other major Type IIS restriction sites (including PaqCI, BsmBI, and SapI) and common restriction sites (including BamHI, BglII, EcoRI, HindIII, KpnI, PstI, SalI, SpeI, SphI, XbaI, XhoI, SphI, and SacII). GFP and RFP dropout cassettes drive strong constitutive expression of sfGFP and mScarlet3, respectively. Colonies retaining the dropout can be readily distinguished by fluorescence and are optimised for rapid screening using a blue-light source and blue-light blocking glasses, enabling straightforward identification of assembled plasmids.

#### T-DNA toolkit and MoClo basics

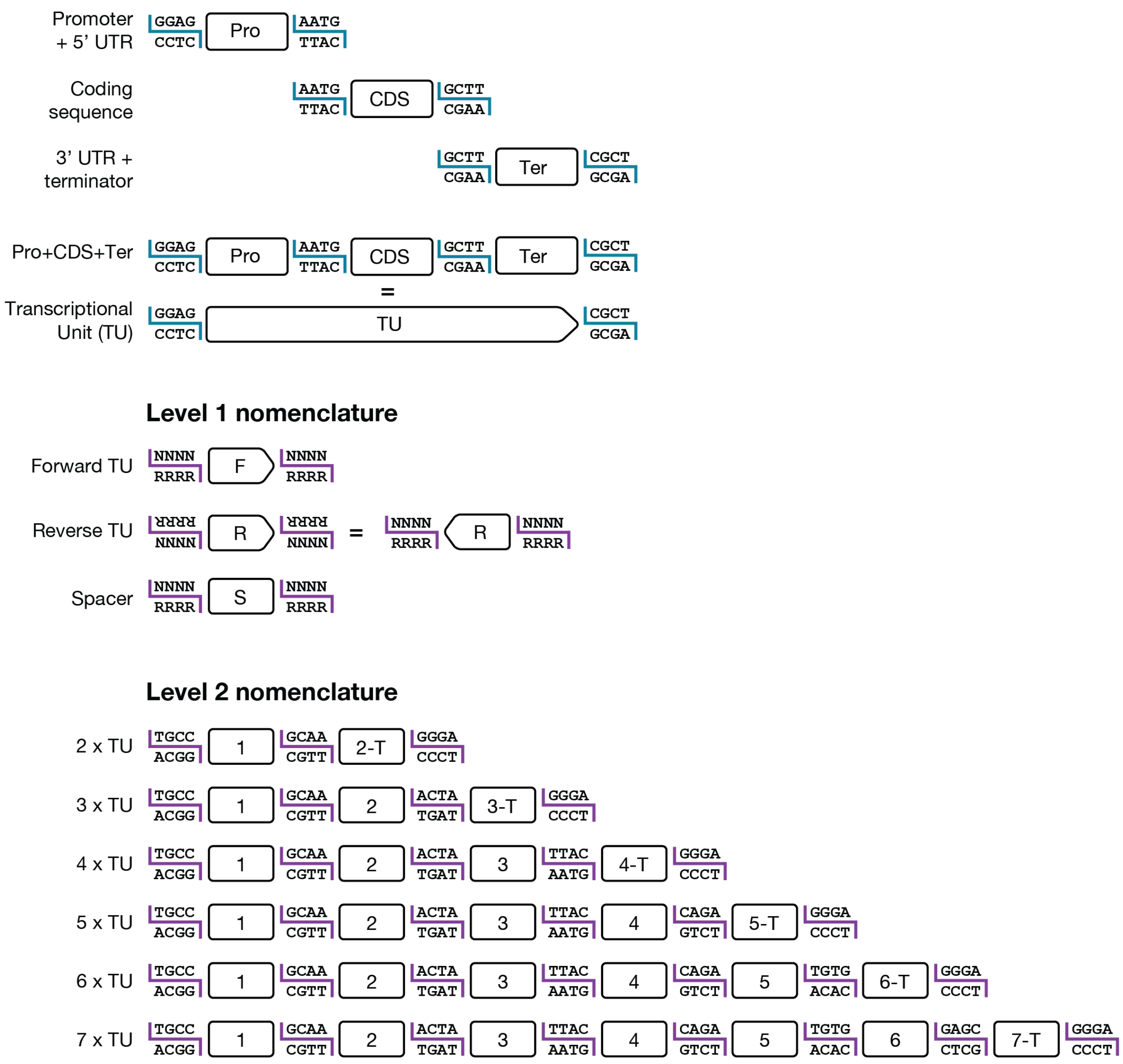


**MoClo formatting and a simplified nomenclature.** To improve accessibility and encourage adoption beyond the synthetic biology community, we propose a streamlined MoClo nomenclature that retains the original framework while simplifying routine use. The Golden Gate Assembly overhangs are shown for Level 0 parts (BsaI, blue) and Level 1 cassettes, (BpiI, purple). All overhangs are taken directly from MoClo for full compatibility of Level 0 parts, Level 1 cassettes, and T-DNA destination vectors.

Firstly, we recommend consolidating Level 0 parts into three core modules: (i) promoter + 5′ UTR (Pro), (ii) complete coding sequence (CDS), and (iii) 3′ UTR + terminator (Ter). Together, these components form a transcriptional unit (TU). Although the original MoClo system maximised modularity by separating promoters from 5′ UTRs and signal peptides from CDS elements, this granularity is less critical in the era of inexpensive full-plasmid sequencing where Level 0 parts can be validated in a single sequencing run regardless of length. While naming Level 0 parts, we find it useful to append them with their Level 0 part definition, e.g. p###_Pro, p###_CDS, and p###_Ter. This simple convention improves organisation during *in silico* construct design and reduces errors when setting up experimental assemblies by making Level 0 part identity and compatibility immediately clear.

We recognise that increased modularity remains valuable in specific contexts, such as combinatorial screening of promoters and 5' UTRs, and the original MoClo format remains fully compatible with this toolkit for advanced users. However, for most basic users, consolidating parts into Pro, CDS, and Ter modules simplifies construct design, reduces cloning complexity, and accelerates assembly workflows.

Secondly, we propose a simplified Level 1 nomenclature that clearly conveys the functional and positional attributes of each cassette. Each module is defined by: (i) its type, transcriptional unit (TU) or spacer (S); (ii) its orientation, forward (F) or reverse (R); and (iii) its position within the multigene construct (positions 1–7). To indicate whether a cassette contains the appropriate overhangs required to terminate a Level 2 multigene assembly, the suffix "–T" is appended. An example illustrating the combined Level 1 and Level 2 nomenclature is provided below for a representative Level 2 assembly. This approach and notation should simplify the design of multigene constructs.

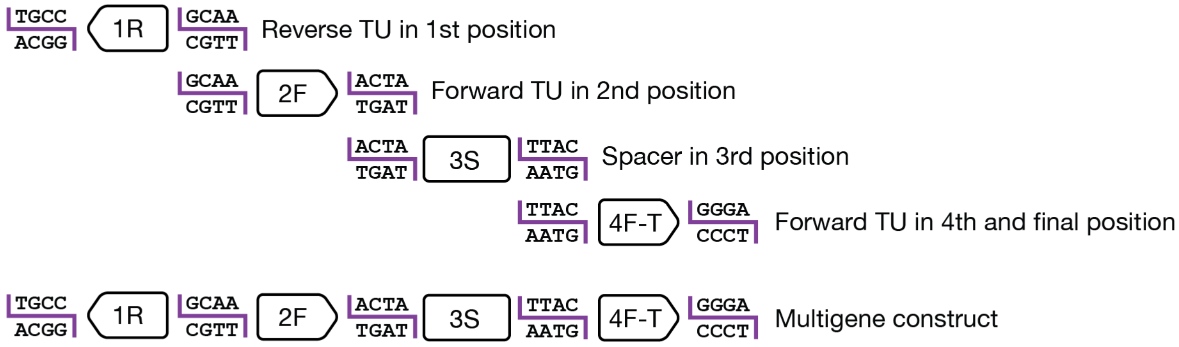


**Combined Level 1 and Level 2 nomenclature.** Example multigene assembly, showing the Level 1 cassettes and their combined nomenclature depending on their type and position within the Level 2 assembly. While naming Level 1 plasmids, we find it useful to append them with their Level 1 definition, e.g. p###_1R, p###_2F, p###_3S, and p###_4F-T. This simple convention improves organisation during *in silico* construct design and reduces errors when setting up experimental assemblies by making Level 1 cassette identity and compatibility immediately clear.

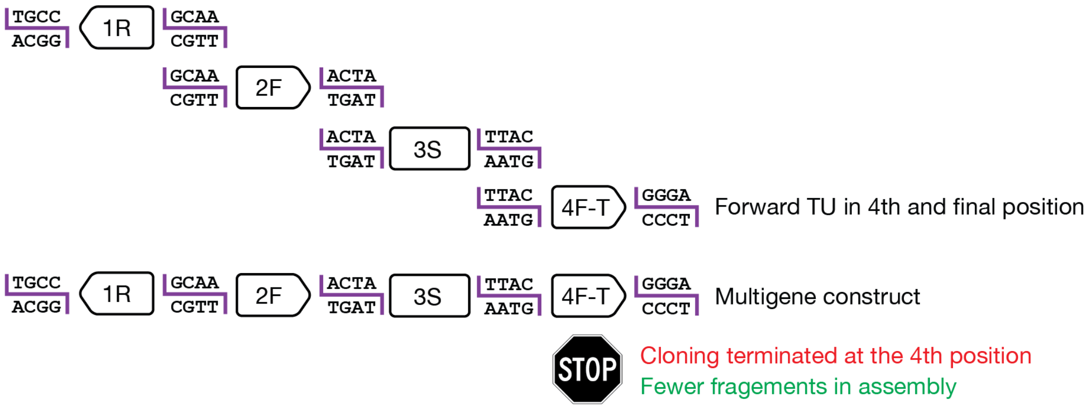


Strategy 2: use of terminal Level 1 spacer in Level 2 assembly

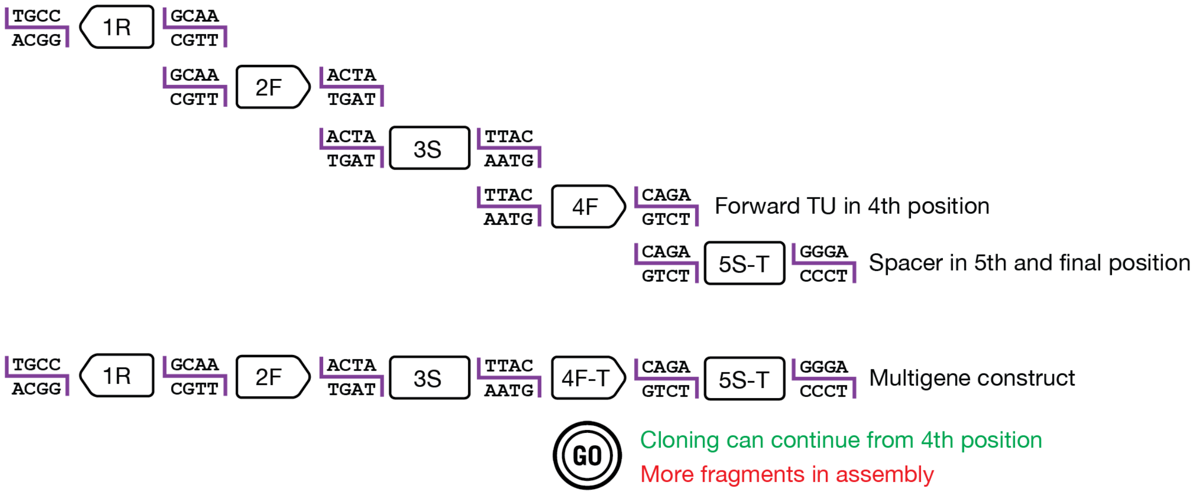


**Alternative cloning approaches for Level 2 multigene constructs.** Within the MoClo framework, Level 2 assemblies can be generated using two distinct approaches:

1. **Defined assembly strategy** – The exact number of required Level 1 transcriptional units (TUs) is assembled, with the final cassette formatted to terminate the multigene construct.
2. **Expandable assembly strategy** – An indeterminate number of Level 1 cassettes is assembled, and the construct is terminated using a dedicated terminal Level 1 spacer.

The defined strategy minimises the number of fragments in the Level 2 reaction, typically improving assembly efficiency and cloning success. However, it offers less flexibility for future expansion. In contrast, the expandable strategy introduces an additional fragment (a terminal spacer) into the Level 2 assembly, which can slightly reduce assembly efficiency due to more parts. Its advantage is modularity, allowing additional TUs to be incorporated later without redesigning existing Level 1 cassettes. The optimal strategy depends on downstream goals. If future expansion of the construct is anticipated, inclusion of a terminal spacer is advisable. If the number of transcriptional units is fixed, reducing fragment number will simplify cloning and maximise efficiency. Importantly, constructs generated using the defined strategy can still be adapted later by reassembling the final cassette in a non-terminal Level 1 format if expansion becomes necessary.

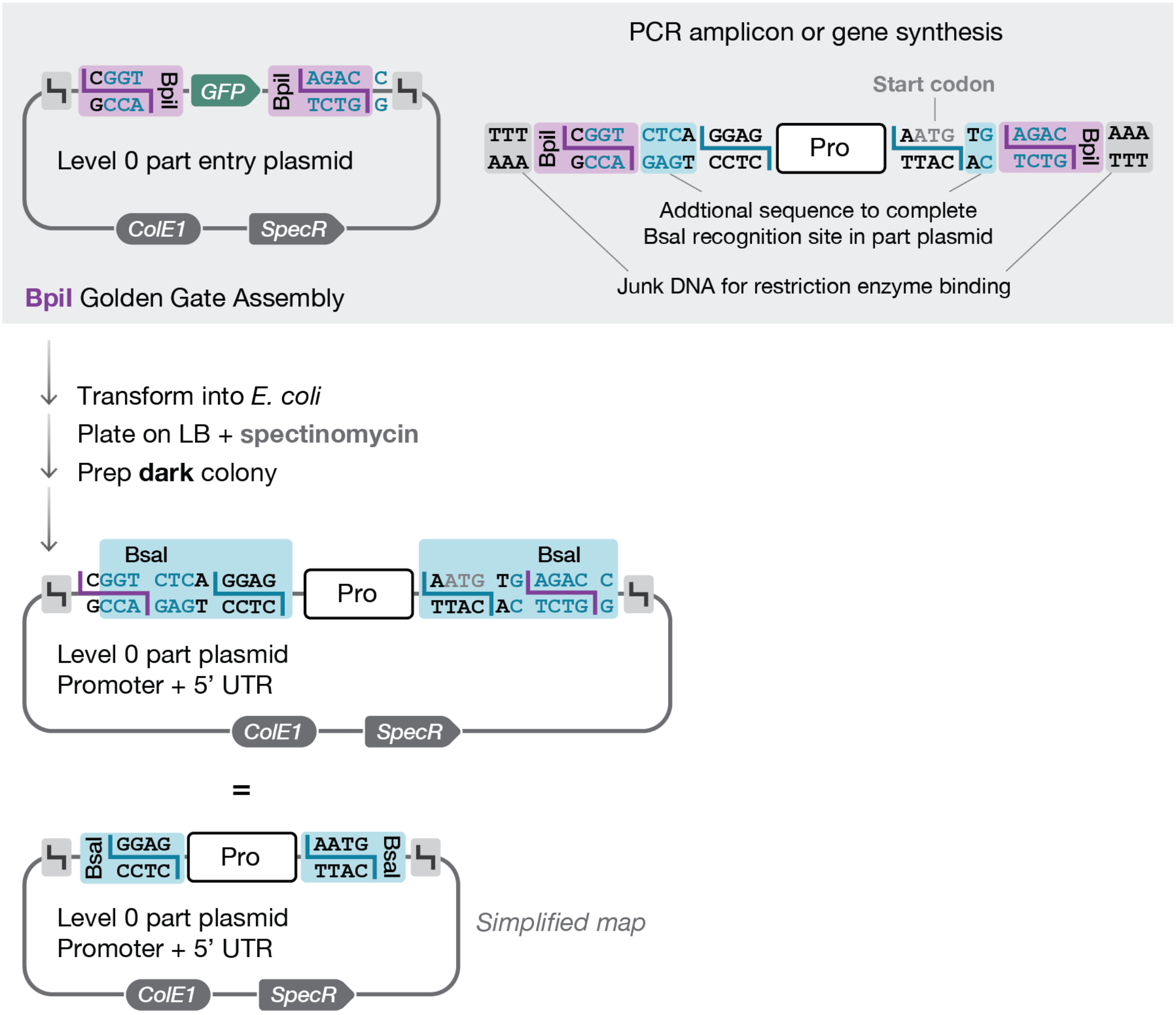


**Assembling promoter + 5' UTR (Pro) sequences into the part entry plasmid.** New promoter + 5′ UTR (Pro) parts can be generated by PCR amplification or DNA synthesis and cloned into the Level 0 part entry plasmid using BpiI Golden Gate Assembly (GGA). In both cases, the Pro part sequence must be flanked by the appropriate MoClo formatting sequences to introduce BpiI recognition sites and the correct Level 0 overhangs, as illustrated above. The start codon (ATG) for the downstream coding sequence is included within the 3′ overhang. Successful cloning replaces the GFP dropout cassette with the new Pro part sequence and generates inward-facing BsaI sites for subsequent Level 1 assembly. Following BpiI GGA, constructs are transformed into *E. coli* and selected on LB agar containing spectinomycin. Non-fluorescent (dark) colonies (indicating loss of the GFP dropout) should be selected for plasmid preparation. A NotI diagnostic digest can be used to screen plasmid preps for the correct size insert. However, all newly generated Level 0 parts must be fully sequence-verified. Errors introduced during PCR amplification or DNA synthesis will propagate through higher-level assemblies, as Level 0 parts form the foundation of all downstream constructs.

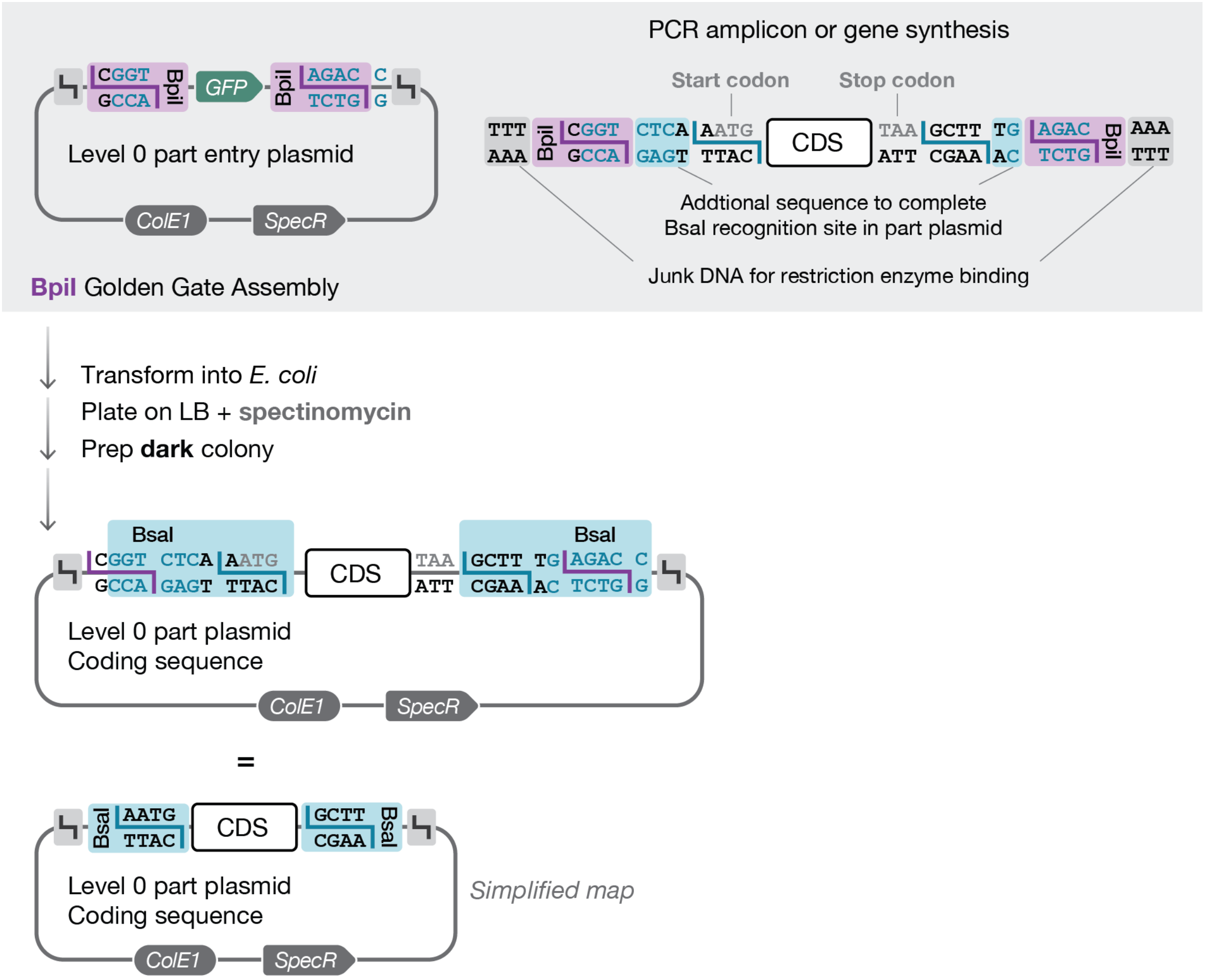


**Assembling coding sequences (CDS) into the part entry plasmid.** New coding sequence (CDS) parts can be generated by PCR amplification or DNA synthesis and cloned into the Level 0 part entry plasmid using BpiI Golden Gate Assembly (GGA). In both cases, the CDS part sequence must be flanked by the appropriate MoClo formatting sequences to introduce BpiI recognition sites and the correct Level 0 overhangs, as illustrated above. The start codon (ATG) of the CDS is included within the 5′ overhang. An in-frame stop codon (e.g. TAA) should be included at the end of the CDS. Successful cloning replaces the GFP dropout cassette with the new CDS part sequence and generates inward-facing BsaI sites for subsequent Level 1 assembly. Following BpiI GGA, constructs are transformed into *E. coli* and selected on LB agar containing spectinomycin. Non-fluorescent (dark) colonies (indicating loss of the GFP dropout) should be selected for plasmid preparation. A NotI diagnostic digest can be used to screen plasmid preps for the correct size insert. However, all newly generated Level 0 parts must be fully sequence-verified. Errors introduced during PCR amplification or DNA synthesis will propagate through higher-level assemblies, as Level 0 parts form the foundation of all downstream constructs.

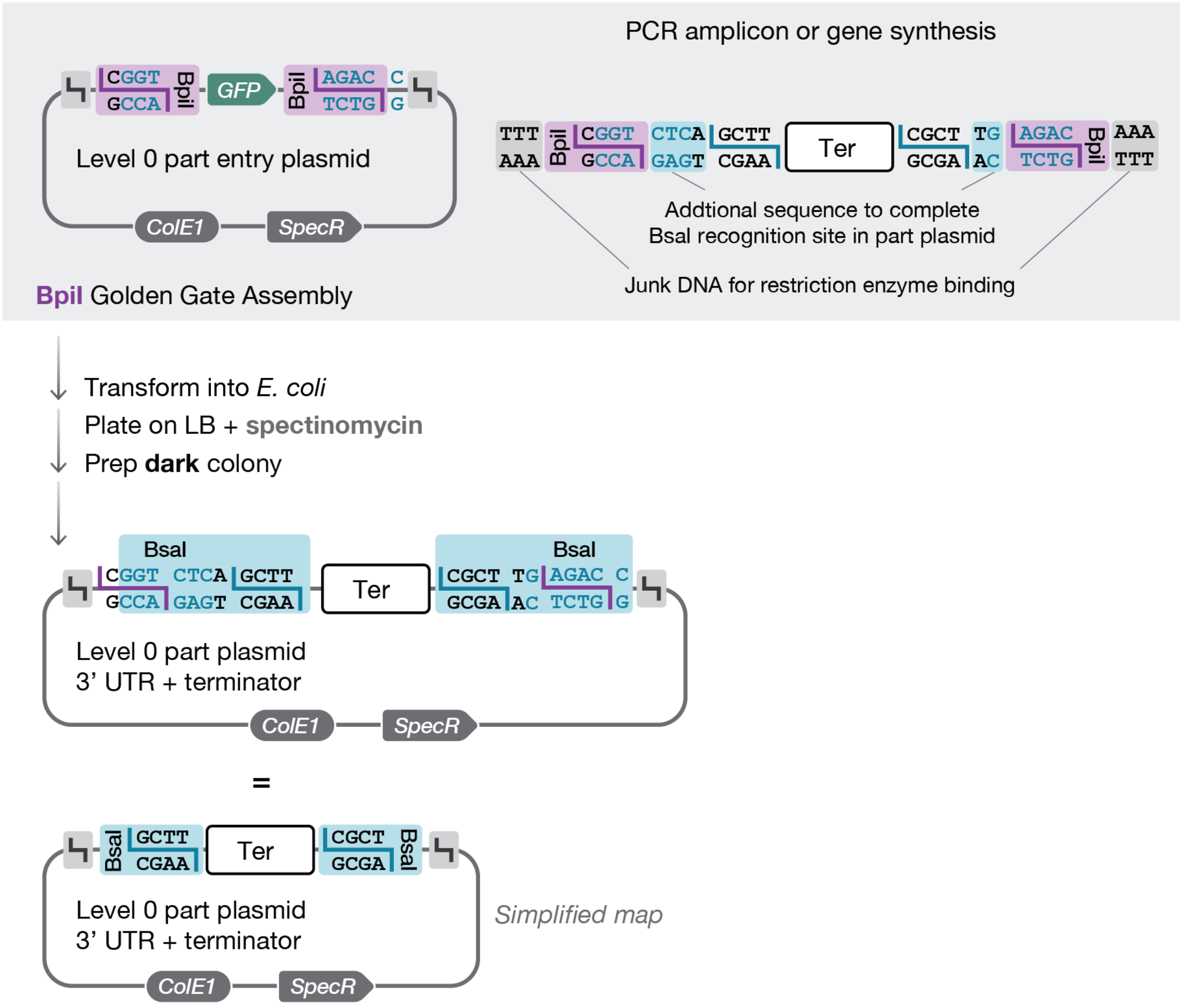


**Assembling 3’ UTR + terminator (Ter) sequences into the part entry plasmid.** New 3’ UTR + terminator (Ter) parts can be generated by PCR amplification or DNA synthesis and cloned into the Level 0 part entry plasmid using BpiI Golden Gate Assembly (GGA). In both cases, the Ter part sequence must be flanked by the appropriate MoClo formatting sequences to introduce BpiI recognition sites and the correct Level 0 overhangs, as illustrated above. Successful cloning replaces the GFP dropout cassette with the new Ter part sequence and generates inward-facing BsaI sites for subsequent Level 1 assembly. Following BpiI GGA, constructs are transformed into *E. coli* and selected on LB agar containing spectinomycin. Non-fluorescent (dark) colonies (indicating loss of the GFP dropout) should be selected for plasmid preparation. A NotI diagnostic digest can be used to screen plasmid preps for the correct size insert. However, all newly generated Level 0 parts must be fully sequence-verified. Errors introduced during PCR amplification or DNA synthesis will propagate through higher-level assemblies, as Level 0 parts form the foundation of all downstream constructs.

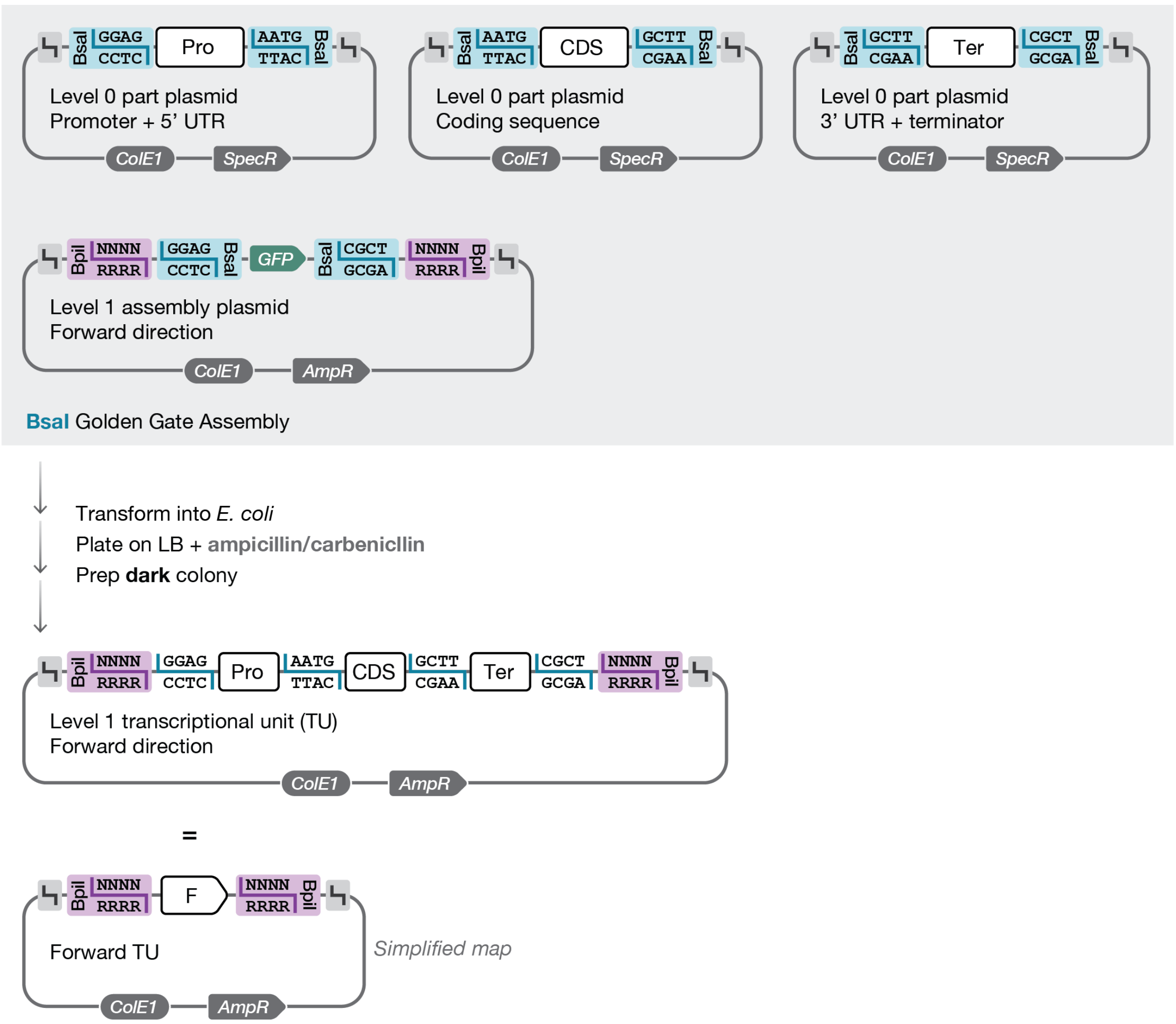


**Assembling a Level 1 transcriptional unit in the forward direction (Forward TU).** Forward transcriptional units (TUs) are assembled by combining a Pro, CDS, and Ter part into a Forward Level 1 assembly plasmid using a BsaI Golden Gate Assembly (GGA). Successful assembly replaces the GFP dropout cassette with the complete TU and introduces inward-facing BpiI sites flanking the insert, enabling subsequent Level 2 multigene assembly. Following BsaI GGA, reactions are transformed into *E. coli* and plated on LB agar containing ampicillin/carbenicillin. Non-fluorescent (dark) colonies (indicating loss of the GFP dropout) should be selected for plasmid preparation. Level 1 assemblies can be validated with a NotI diagnostic digest or full-plasmid sequencing. Additionally, assembled Level 1 transcriptional units can be used in protoplast transformation for the expression of a single gene.

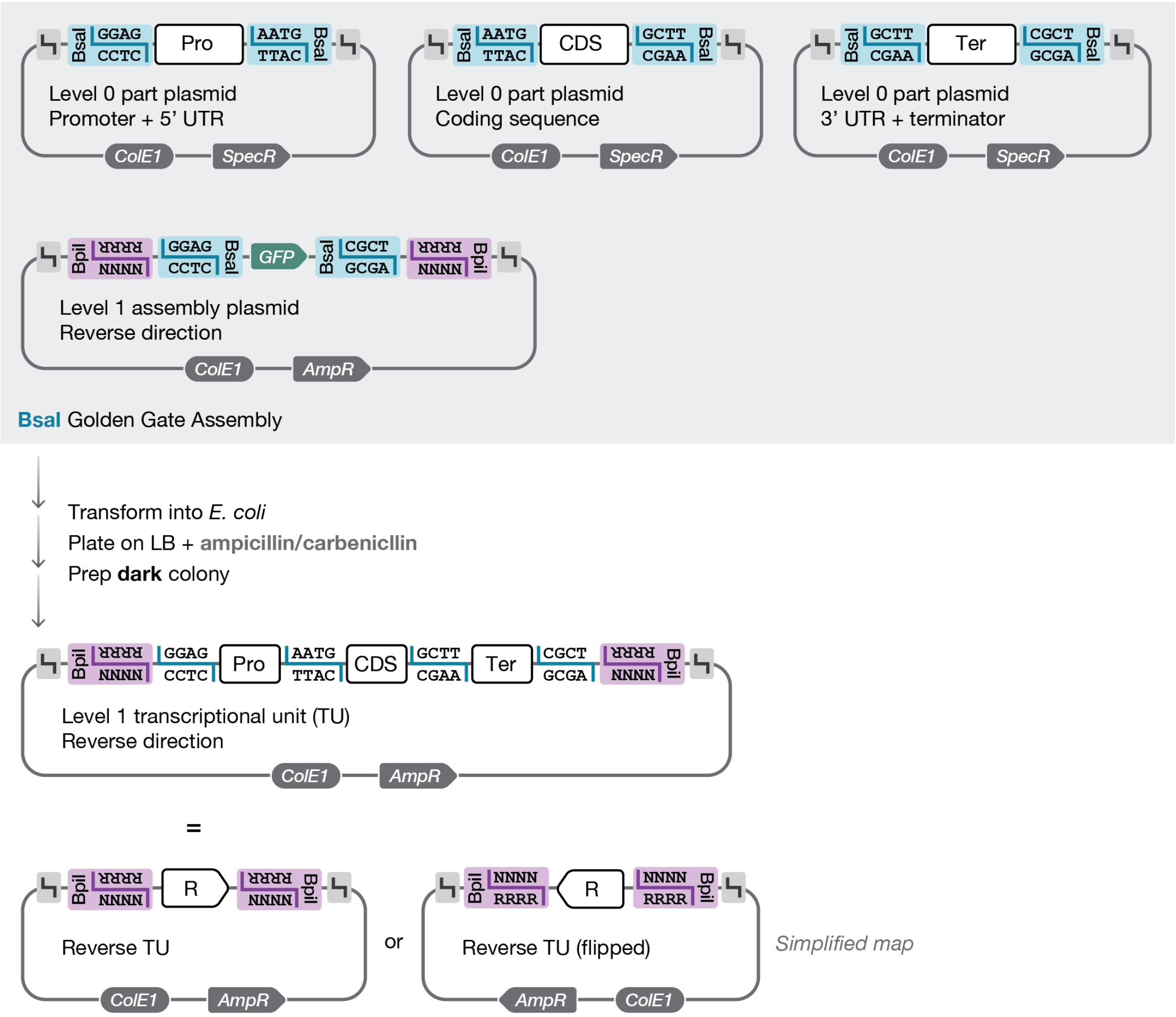


**Assembling a Level 1 transcriptional unit in the reverse direction (Reverse TU).** Transcriptional units (TUs) in the reverse direction are assembled by combining a Pro, CDS, and Ter part into a Reverse Level 1 assembly plasmid using a BsaI Golden Gate Assembly (GGA). Successful cloning replaces the GFP dropout cassette with the assembled TU, flanked by inward-facing BpiI sites for subsequent Level 2 assembly. Following BsaI GGA, constructs are transformed into *E. coli* and selected on LB agar containing ampicillin/carbenicillin. Non-fluorescent (dark) colonies (indicating loss of the GFP dropout) should be selected for plasmid preparation. Level 1 assemblies can be validated with a NotI diagnostic digest or full-plasmid sequencing.

#### Essential MoClo cloning operations

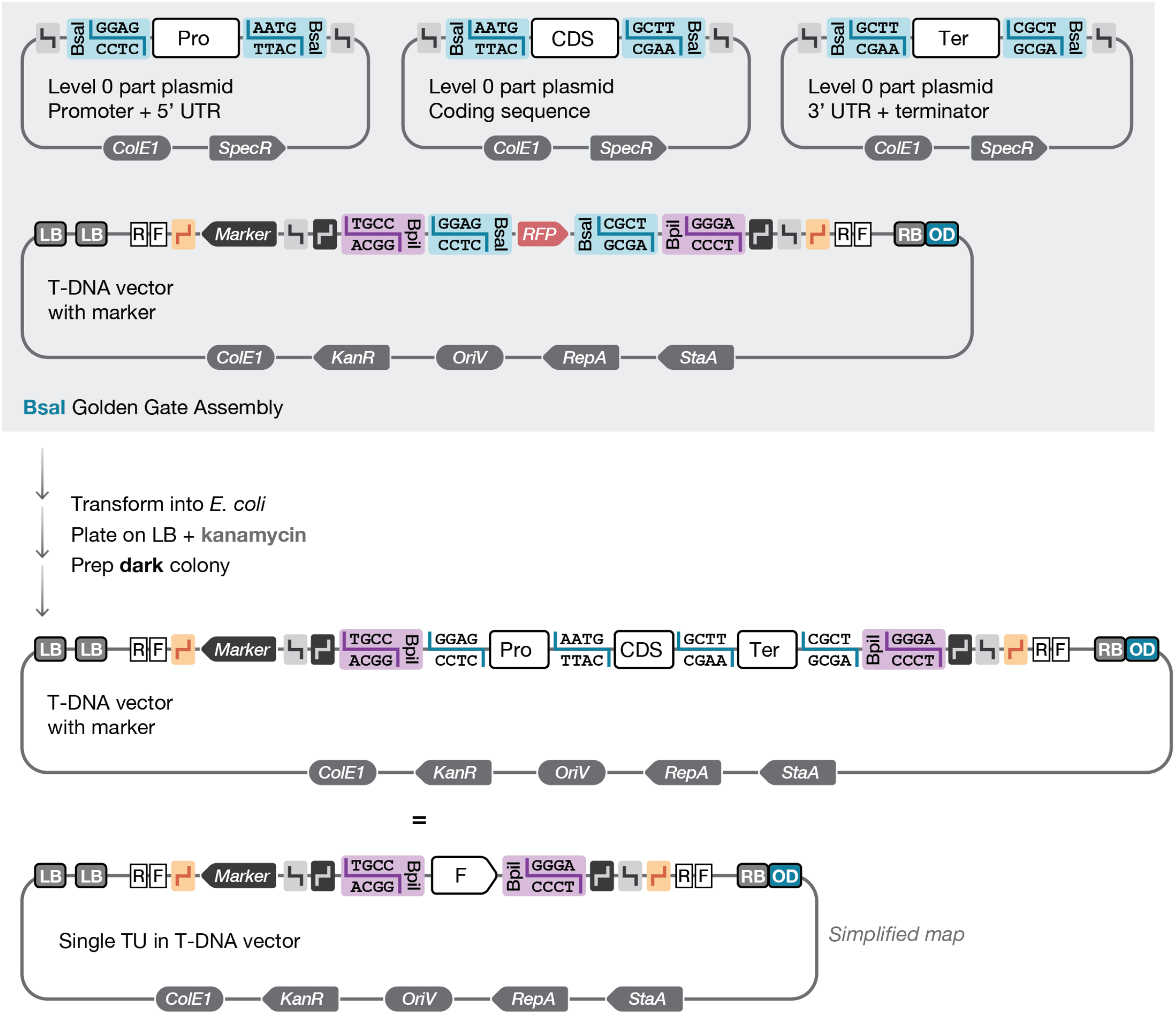


**Assembling a Level 1 transcriptional unit (TU) into a T-DNA vector.** A single transcriptional unit (TU) can be assembled directly into a T-DNA destination vector using BsaI Golden Gate Assembly (GGA). In this assembly, the TU can only be in the forward orientation. If reverse orientation is required, a reverse, non-terminal Level 1 TU (e.g. 1R) should first be generated and then combined with a terminal spacer (e.g. 2S-T) in a subsequent Level 2 assembly. Successful cloning replaces the RFP dropout cassette with the assembled TU. Following BsaI GGA, reactions are transformed into *E. coli* and plated on LB agar containing kanamycin. Non-fluorescent (dark) colonies (indicating loss of the RFP dropout) should be selected for plasmid preparation. Assemblies can be validated by diagnostic NotI digestion. However, we strongly recommend full-plasmid sequencing of assembled T-DNA constructs. Seemingly minor human errors, such as swapping similarly sized parts, can go unnoticed at the cloning stage and result in substantial delays during downstream stable plant transformation experiments.

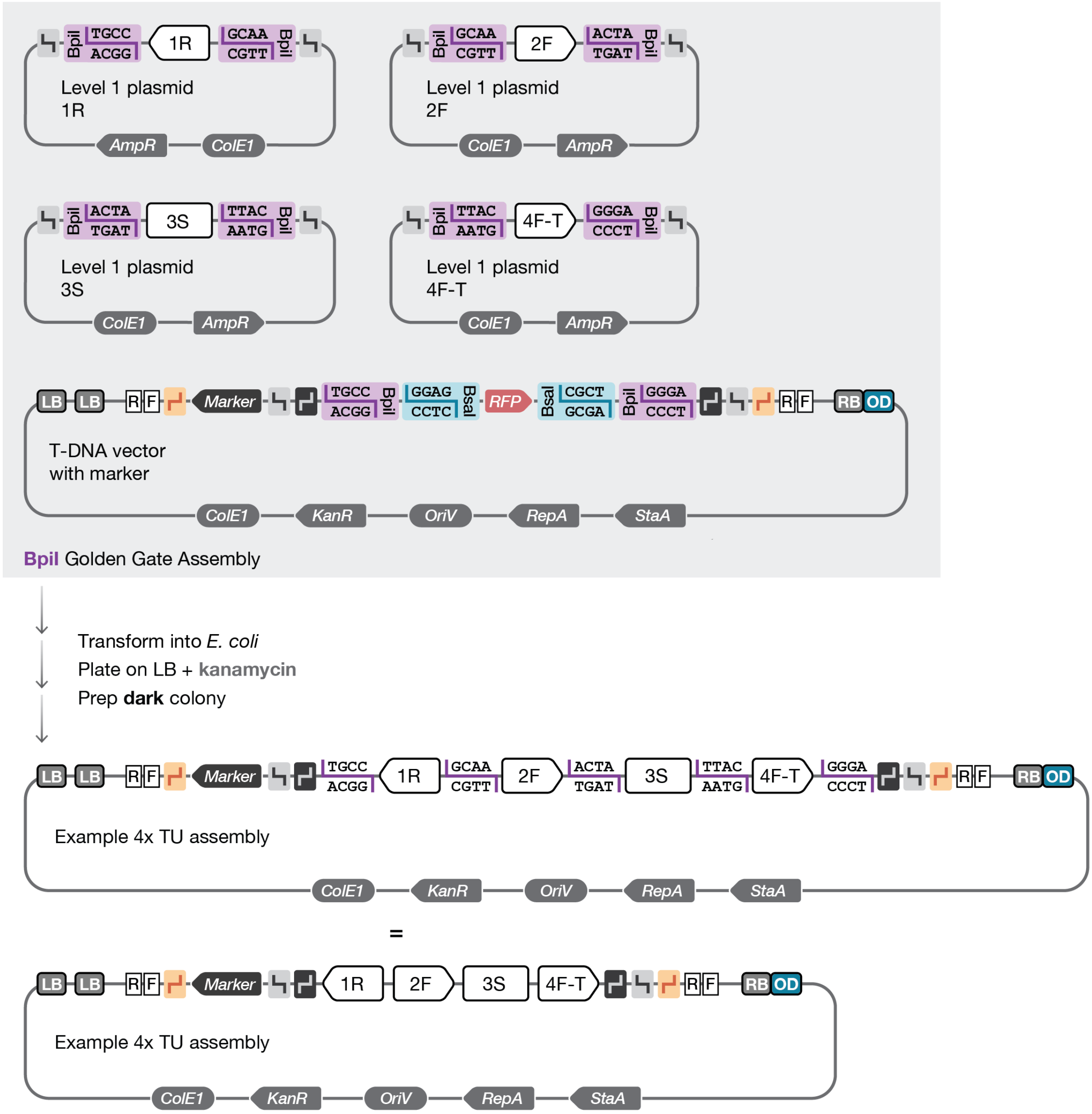


**Example Level 2 multigene assembly into a T-DNA vector.** Multigene constructs are assembled into a T-DNA destination vector from multiple Level 1 cassettes using BpiI Golden Gate Assembly (GGA). Level 1 plasmids must be arranged in consecutive, uninterrupted order (starting from position 1) and the series must terminate with a terminal cassette (T), whether this is a spacer (S) or a transcriptional unit (TU). Successful assembly replaces the RFP dropout cassette with the complete multigene construct. Following BpiI GGA, reactions are transformed into *E. coli* and plated on LB agar containing kanamycin. Non-fluorescent (dark) colonies (indicating loss of the RFP dropout) should be selected for plasmid preparation. Assemblies can initially be screened by diagnostic NotI digestion. However, we strongly recommend full-plasmid sequencing of assembled T-DNA constructs. Seemingly minor human errors, such as swapping similarly sized Level 0 parts or Level 1 cassettes, can go unnoticed during the cloning stages and result in substantial delays during downstream stable plant transformation experiments.

#### Advanced MoClo cloning operations

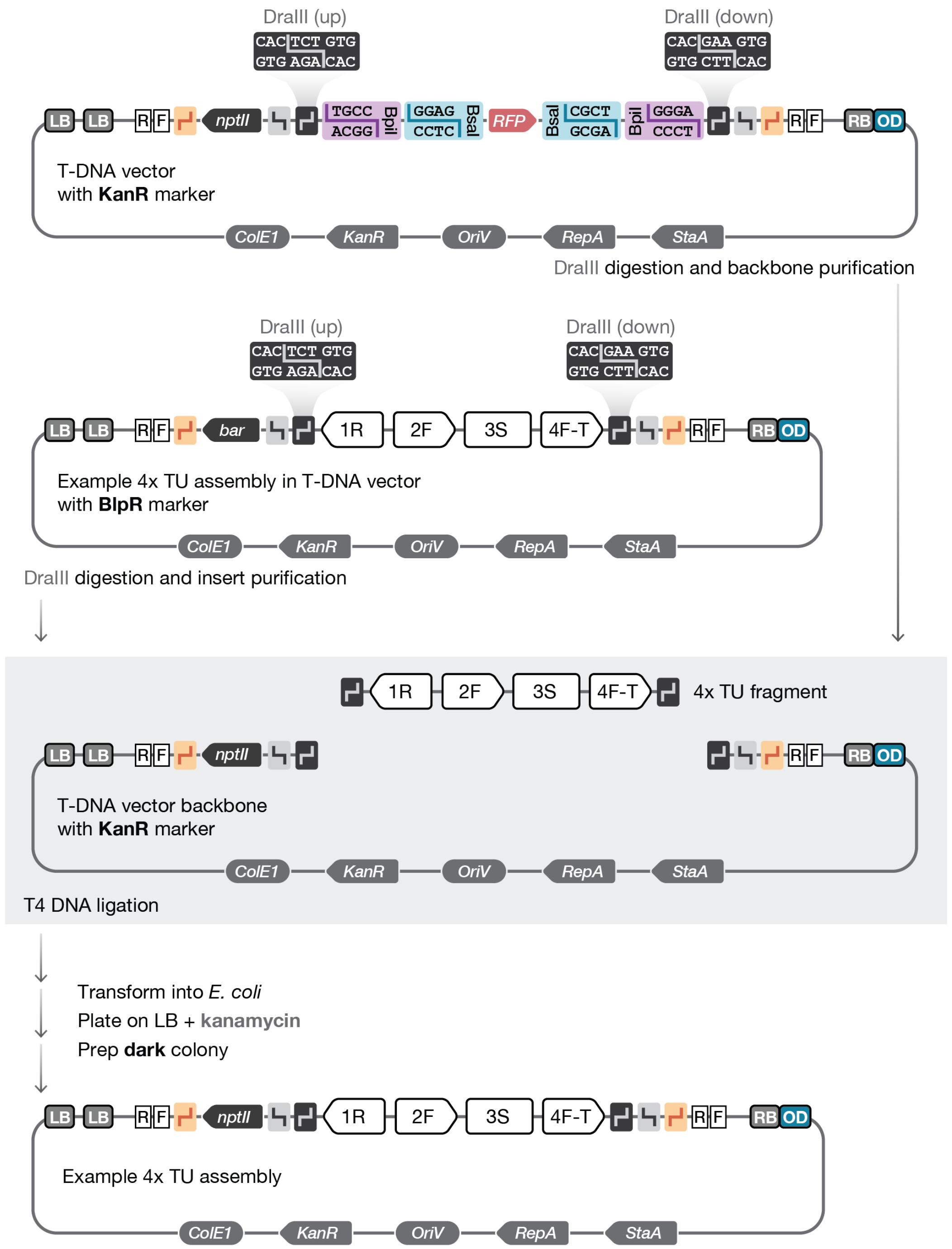


**Moving assembled constructs between T-DNA vectors.** Assembled transcriptional units (TUs) or multigene constructs can be transferred between T-DNA destination vectors using a DraIII digestion–ligation reaction that preserves the same 3-bp overhangs used in the MoClo system, as shown above. In this approach, both the T-DNA insert and the destination vector backbone are digested with DraIII and gel purified. The purified insert is then ligated into the destination backbone, and the reaction is transformed into *E. coli* and plated on LB agar containing kanamycin. Non-fluorescent (dark) colonies (indicating loss of the RFP dropout) should be selected for plasmid preparation. If the insert has been previously sequence-validated, a NotI diagnostic digest can be used to confirm the expected insert size. However, we recommend full-plasmid sequencing after transfer into a new backbone. Residual plasmid carryover from the original T-DNA vector can be difficult to distinguish by restriction digest alone, and colony PCR may yield false positives due to the abundance of the destination backbone following transformation. The DraIII cloning sites can also be used to introduce the original MoClo cloning sequences into the new T-DNA vectors or to integrate alternative cloning systems if required.

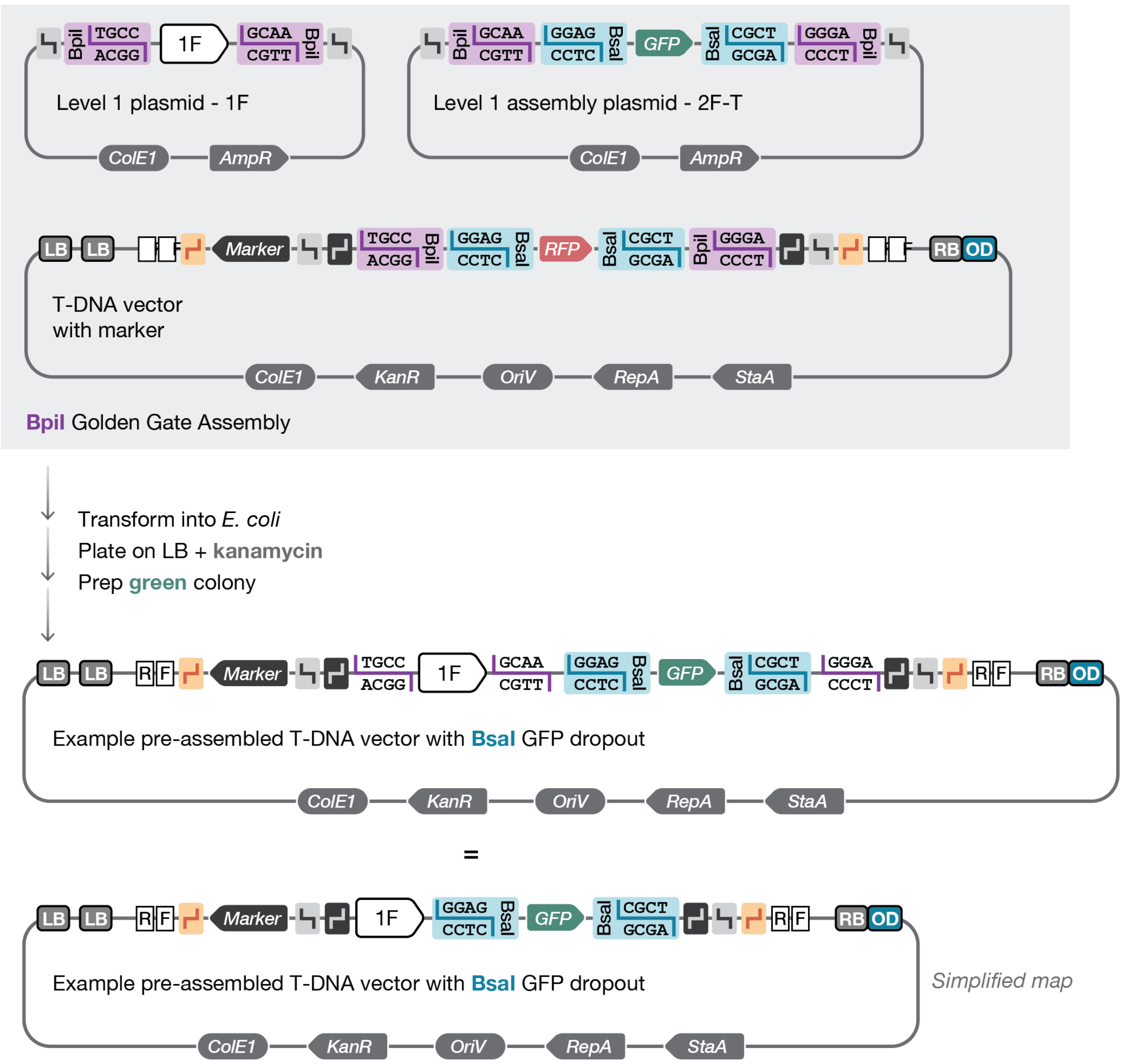


**Assembling multigene constructs for future BsaI transcriptional unit expansion.** Level 2 multigene constructs can be assembled with an unassembled Level 1 plasmid. This allows for future transcriptional unit (TU) assembly or multigene expansion. Introducing the expansion module resets the T-DNA plasmid for future multigene assembly, which is necessary for introducing more than 7 genes. Future expansion is also useful for creating a new T-DNA vector that has common cassettes preassembled, if they are likely to be used a lot, reducing the number of fragments in later Golden Gate Reactions (GGAs). Multigene constructs with BsaI expansion are assembled into a T-DNA destination vector from one or more Level 1 cassettes and an unassembled Level 1 assembly plasmid using BpiI GGA. Following BpiI GGA, reactions are transformed into *E. coli* and plated on LB agar containing kanamycin. Green-fluorescent colonies (indicating the exchange of the RFP dropout for GFP) should be selected for plasmid preparation. Assemblies can be validated by diagnostic NotI digestion at this stage, but we recommend full-plasmid sequencing once the final T-DNA has been cloned.

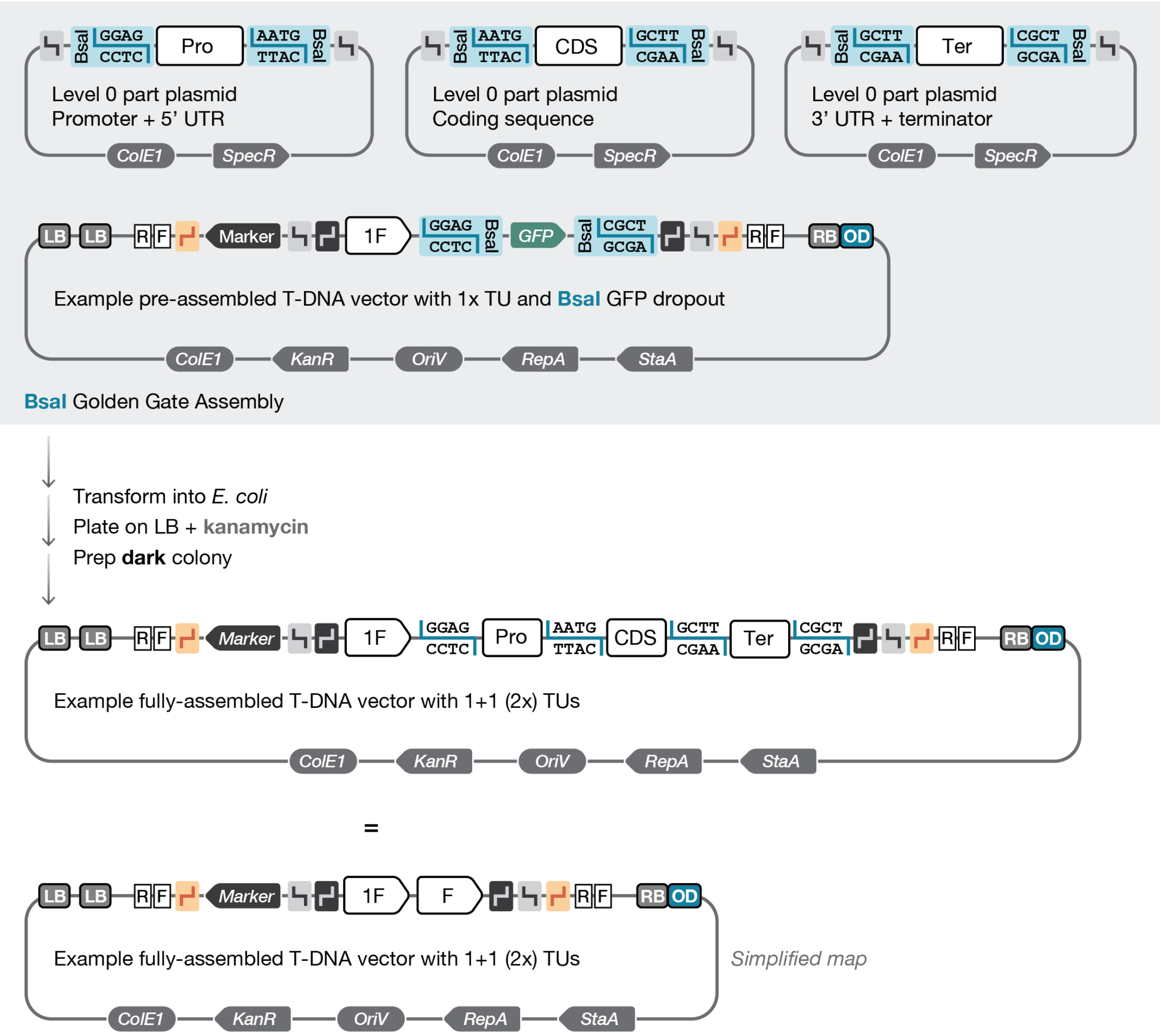


**Assembling a transcriptional unit into a BsaI expandable T-DNA vector.** Following construction of a pre-assembled multigene vector containing a BsaI dropout cassette, transcriptional units (TUs) can be assembled directly into the vector using BsaI Golden Gate Assembly (GGA). Each TU is generated by combining a Pro, CDS, and Ter part and the BsaI expandable T-DNA vector. Successful assembly replaces the GFP dropout cassette with the complete TU. After the BsaI GGA reaction, assemblies are transformed into *E. coli* and plated on LB agar containing kanamycin. Non-fluorescent (dark) colonies (indicating loss of the GFP dropout) should be selected for plasmid preparation. Assemblies can be validated by diagnostic NotI digestion. However, we strongly recommend full-plasmid sequencing of assembled T-DNA constructs. Seemingly minor human errors, such as swapping similarly sized parts, can go unnoticed at the cloning stage and result in substantial delays during downstream stable plant transformation experiments.

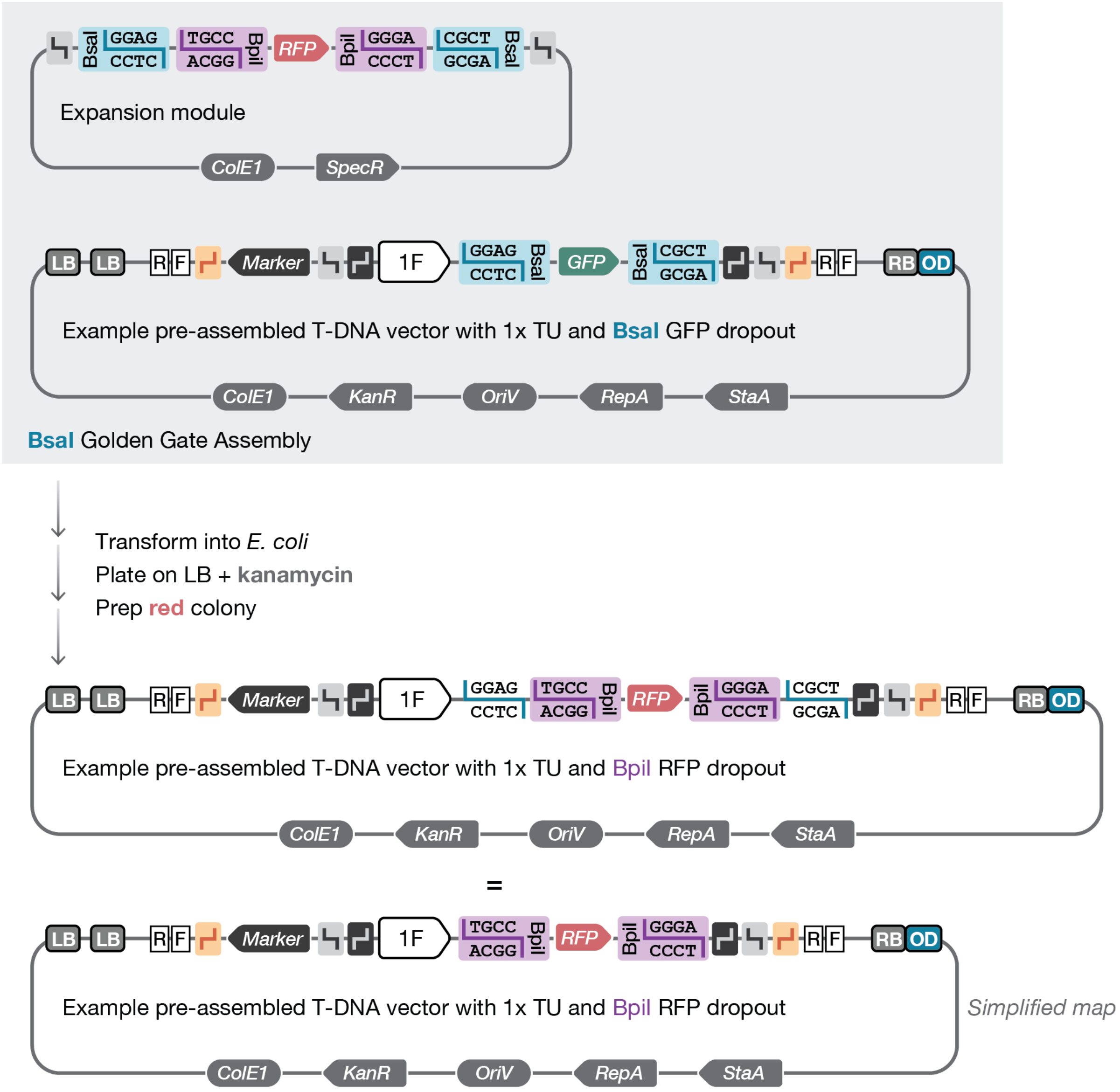


**Converting a BsaI expandable T-DNA vector for BpiI multigene expansion.** To extend a T-DNA construct design beyond the seven cassettes supported by the available Level 1 assembly plasmids, a BsaI-expandable T-DNA vector can be converted into a BpiI-expandable vector by introducing the expansion module. This conversion is achieved by assembling the expansion module plasmid together with the BsaI-expandable T-DNA vector using a BsaI Golden Gate Assembly (GGA). Following BsaI GGA, reactions are transformed into *E. coli* and plated on LB agar containing kanamycin. Red-fluorescent colonies (indicating the exchange of the GFP dropout for RFP) should be selected for plasmid preparation. Assemblies can be validated by diagnostic NotI digestion at this stage. This conversion resets the T-DNA vector for further rounds of multigene assembly, enabling iterative expansion of the construct. In principle, this process can be repeated multiple times; however, cloning efficiency typically decreases as plasmid size increases.

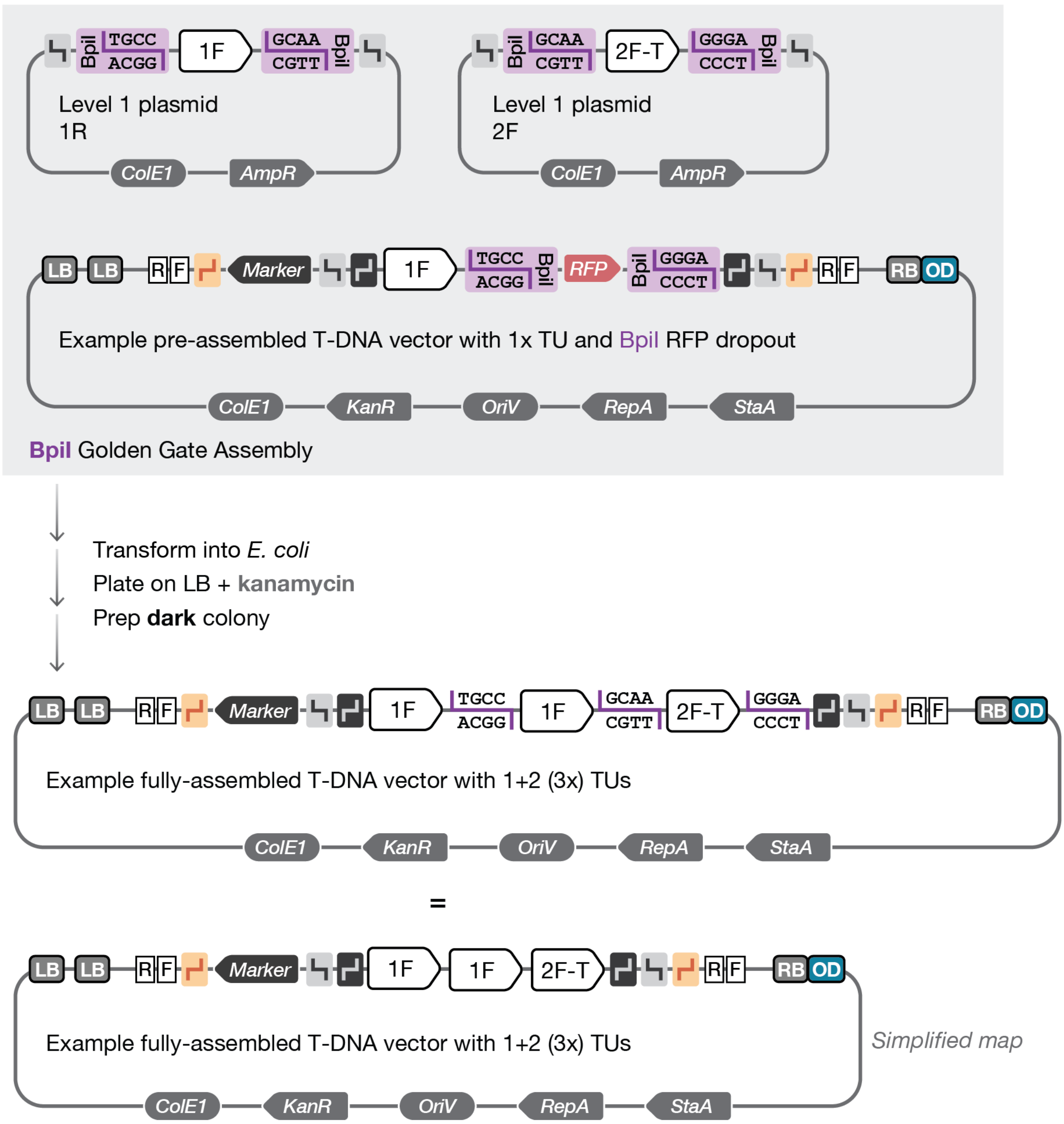


**Assembling a multigene construct into a BpiI expandable T-DNA vector.** Multigene constructs are assembled into a BpiI expandable T-DNA destination vector from multiple Level 1 cassettes using BpiI Golden Gate Assembly (GGA). Level 1 plasmids must be arranged in consecutive, uninterrupted order (starting from position 1) and the series must terminate with a terminal cassette (T), whether this is a spacer (S) or a transcriptional unit (TU). Successful assembly replaces the RFP dropout cassette with the complete multigene construct. Following BpiI GGA, reactions are transformed into *E. coli* and plated on LB agar containing kanamycin. Non-fluorescent (dark) colonies (indicating loss of the RFP dropout) should be selected for plasmid preparation. Assemblies can initially be screened by diagnostic NotI digestion. However, we strongly recommend full-plasmid sequencing of assembled T-DNA constructs. Seemingly minor human errors, such as swapping similarly sized Level 0 parts or Level 1 cassettes, can go unnoticed during the cloning stages and result in substantial delays during downstream stable plant transformation experiments.

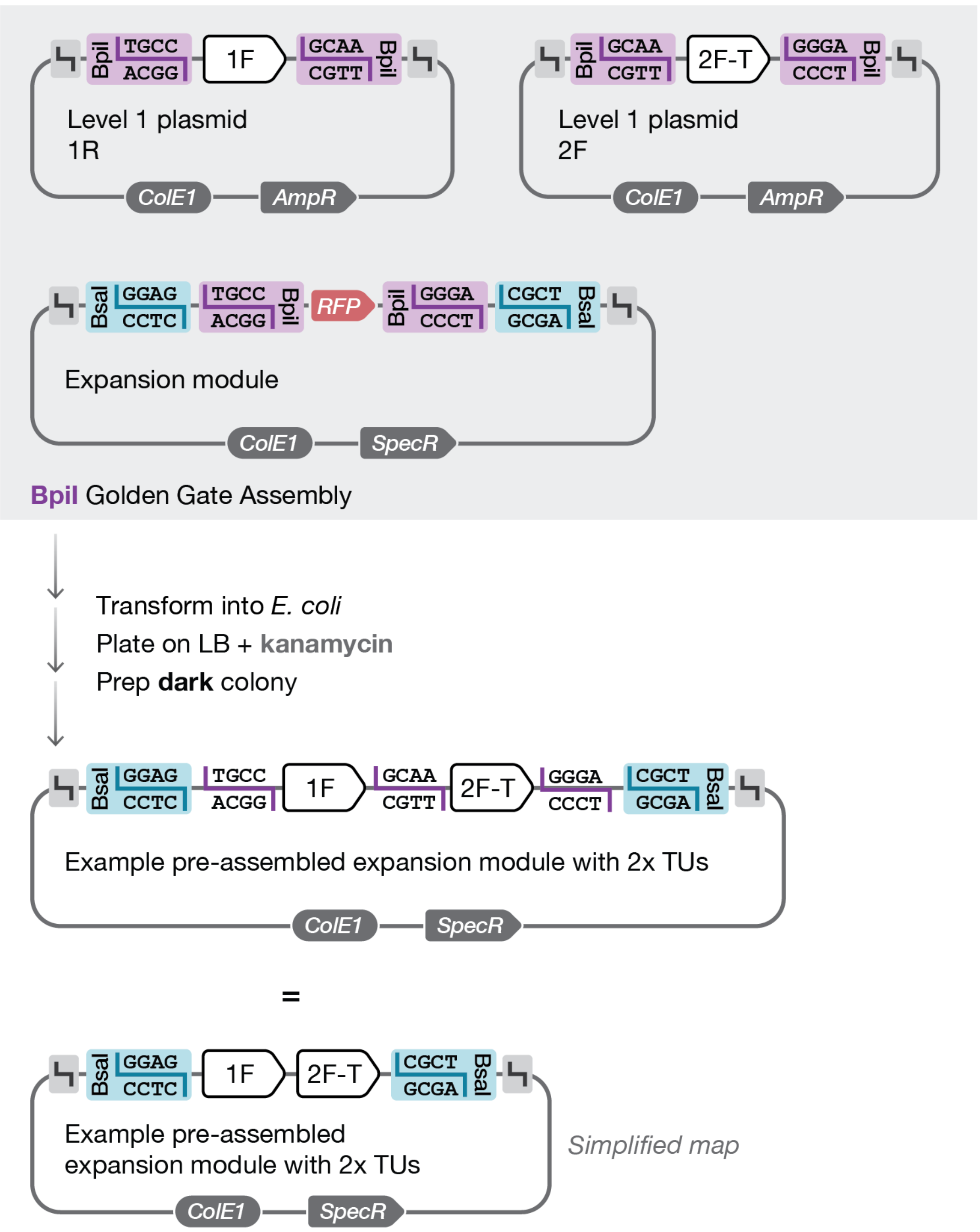


**Assembling a multigene construct into the expansion module plasmid.** Multigene constructs can be assembled into the expansion module plasmid prior to transfer into a T-DNA vector. This approach allows multiple genes to be combined into a single reusable module that can function similarly to a Level 0 transcriptional unit (TU) part in subsequent assemblies. In addition, the expansion module plasmid can be used directly for plant protoplast transformation to enable the transient expression of multiple genes. Multigene constructs are assembled into the expansion module from multiple Level 1 cassettes using BpiI Golden Gate Assembly (GGA). Level 1 plasmids must be arranged in consecutive, uninterrupted order (starting from position 1) and must terminate with a terminal cassette (–T), either a spacer (S) or a transcriptional unit (TU). Successful assembly replaces the RFP dropout cassette with the complete multigene construct. Following BpiI GGA, reactions are transformed into *E. coli* and plated on LB agar containing spectinomycin. Non-fluorescent (dark) colonies (indicating loss of the RFP dropout) should be selected for plasmid preparation. Assemblies can be screened by diagnostic NotI digestion. However, if this is going to be treated as a new part, it is worth validating at this stage to provide propagating cloning errors to later stages.
