## Supplementary Tables for "Rational design of T-DNA vectors enables predictable, single-copy integration in *Arabidopsis thaliana*"

**Extended Data Table 1:** Transformation counts

| Figure | Description | Line | Seed weight (mg) | Seed count | RUBY- | RUBY+ | Total | Tx RUBY- (%) | Tx RUBY+ (%) | Tx total (%) | Fraction RUBY+ (%) | RUBY+ per 10000 seed |
|---|---|---|---|---|---|---|---|---|---|---|---|---|
| Fig. 1i | Starter vector | AWS429 | 900 | 45000 | 730 | 58 | 788 | 1.62 | 0.13 | 1.75 | 7.36 | 12.89 |
| Fig. 2b | Wild type | AWS493 | 400 | 20000 | 257 | 28 | 285 | 1.29 | 0.14 | 1.43 | 9.82 | 14.00 |
| Fig. 2b | Border Var 1 | AWS494 | 400 | 20000 | 419 | 40 | 459 | 2.10 | 0.20 | 2.30 | 8.71 | 20.00 |
| Fig. 2b | Border Var 2 | AWS495 | 400 | 20000 | 394 | 37 | 431 | 1.97 | 0.19 | 2.16 | 8.58 | 18.50 |
| Fig. 2b | Border Var 3 | AWS496 | 400 | 20000 | 326 | 26 | 352 | 1.63 | 0.13 | 1.76 | 7.39 | 13.00 |
| Fig. 2b | Border Var 4 | AWS497 | 1000 | 50000 | 12 | 90 | 102 | 0.02 | 0.18 | 0.20 | 88.24 | 18.00 |
| Fig. 2b | Border Var 5 | AWS590 | 400 | 20000 | 302 | 35 | 337 | 1.51 | 0.18 | 1.69 | 10.39 | 17.50 |
| Fig. 2b | Border Var 6 | AWS499 | 600 | 30000 | 30 | 91 | 121 | 0.10 | 0.30 | 0.40 | 75.21 | 30.33 |
| Fig. 2e | RB-Max | AWS619 | 400 | 20000 | 226 | 9 | 235 | 1.13 | 0.05 | 1.18 | 3.83 | 4.50 |
| Fig. 2e | RB-Mid | AWS620 | 400 | 20000 | 32 | 69 | 101 | 0.16 | 0.35 | 0.51 | 68.32 | 34.50 |
| Fig. 2e | RB-Min | AWS621 | 400 | 20000 | 8 | 32 | 40 | 0.04 | 0.16 | 0.20 | 80.00 | 16.00 |
| Fig. 2g | OD 2.0, p-RB-Max | AWS601 | 400 | 20000 | 115 | 4 | 119 | 0.58 | 0.02 | 0.60 | 3.36 | 2.00 |
| Fig. 2g | OD 2.0, p-RB-Mid | AWS602 | 400 | 20000 | 40 | 72 | 112 | 0.20 | 0.36 | 0.56 | 64.29 | 36.00 |
| Fig. 2g | OD 2.0, p-RB-Min | AWS603 | 400 | 20000 | 12 | 47 | 59 | 0.06 | 0.24 | 0.30 | 79.66 | 23.50 |
| Fig. 2g | OD 0.8, p-RB-Max | AWS607 | 400 | 20000 | 215 | 8 | 223 | 1.08 | 0.04 | 1.12 | 3.59 | 4.00 |
| Fig. 2g | OD 0.8, p-RB-Mid | AWS610 | 400 | 20000 | 29 | 50 | 79 | 0.15 | 0.25 | 0.40 | 63.29 | 25.00 |
| Fig. 2g | OD 0.8, p-RB-Min | AWS613 | 400 | 20000 | 6 | 22 | 28 | 0.03 | 0.11 | 0.14 | 78.57 | 11.00 |
| Fig. 2g | OD 0.2, p-RB-Max | AWS604 | 400 | 20000 | 165 | 6 | 171 | 0.83 | 0.03 | 0.86 | 3.51 | 3.00 |
| Fig. 2g | OD 0.2, p-RB-Mid | AWS605 | 400 | 20000 | 30 | 68 | 98 | 0.15 | 0.34 | 0.49 | 69.39 | 34.00 |
| Fig. 2g | OD 0.2, p-RB-Min | AWS606 | 400 | 20000 | 14 | 29 | 43 | 0.07 | 0.15 | 0.22 | 67.44 | 14.50 |
| Fig. 2h | EHA105, p-RB-Max | AWS439 | 400 | 20000 | 72 | 6 | 78 | 0.36 | 0.03 | 0.39 | 7.69 | 3.00 |
| Fig. 2h | EHA105, p-RB-Mid | AWS441 | 400 | 20000 | 19 | 5 | 24 | 0.10 | 0.03 | 0.12 | 20.83 | 2.50 |
| Fig. 2h | EHA105, p-RB-Min | AWS442 | 400 | 20000 | 0 | 1 | 1 | 0.00 | 0.01 | 0.01 | 100.00 | 0.50 |
| Fig. 2h | LBA4404, p-RB-Max | AWS625 | 400 | 20000 | 30 | 23 | 53 | 0.15 | 0.12 | 0.27 | 43.40 | 11.50 |
| Fig. 2h | LBA4404, p-RB-Mid | AWS626 | 400 | 20000 | 1 | 2 | 3 | 0.01 | 0.01 | 0.02 | 66.67 | 1.00 |
| Fig. 2h | LBA4404, p-RB-Min | AWS627 | 400 | 20000 | 0 | 0 | 0 | 0.00 | 0.00 | 0.00 | 0.00 | 0.00 |
| Fig. 2h | AGL-1, p-RB-Max | AWS628 | 400 | 20000 | 69 | 7 | 76 | 0.35 | 0.04 | 0.38 | 9.21 | 3.50 |
| Fig. 2h | AGL-1, p-RB-Mid | AWS629 | 400 | 20000 | 39 | 17 | 56 | 0.20 | 0.09 | 0.28 | 30.36 | 8.50 |
| Fig. 2h | AGL-1, p-RB-Min | AWS630 | 400 | 20000 | 1 | 4 | 5 | 0.01 | 0.02 | 0.03 | 80.00 | 2.00 |
| Fig. 2i | WS, p-RB-Max | AWS608 | 400 | 20000 | 91 | 15 | 106 | 0.46 | 0.08 | 0.53 | 14.15 | 7.50 |
| Fig. 2i | WS, p-RB-Mid | AWS611 | 400 | 20000 | 20 | 34 | 54 | 0.10 | 0.17 | 0.27 | 62.96 | 17.00 |
| Fig. 2i | WS, p-RB-Min | AWS614 | 400 | 20000 | 0 | 10 | 10 | 0.00 | 0.05 | 0.05 | 100.00 | 5.00 |
| Fig. 2i | Ler, p-RB-Max | AWS609 | 400 | 20000 | 25 | 4 | 29 | 0.13 | 0.02 | 0.15 | 13.79 | 2.00 |
| Fig. 2i | Ler, p-RB-Mid | AWS612 | 400 | 20000 | 10 | 24 | 34 | 0.05 | 0.12 | 0.17 | 70.59 | 12.00 |
| Fig. 2i | Ler, p-RB-Min | AWS615 | 400 | 20000 | 2 | 0 | 2 | 0.01 | 0.00 | 0.01 | 0.00 | 0.00 |
| Fig. 2i | C24, p-RB-Max | AWS529 | 400 | 20000 | 28 | 4 | 32 | 0.14 | 0.02 | 0.16 | 12.50 | 2.00 |
| Fig. 2i | C24, p-RB-Mid | AWS531 | 400 | 20000 | 5 | 8 | 13 | 0.03 | 0.04 | 0.07 | 61.54 | 4.00 |
| Fig. 2i | C24, p-RB-Min | AWS532 | 400 | 20000 | 2 | 7 | 9 | 0.01 | 0.04 | 0.05 | 77.78 | 3.50 |
| Fig. 3e | pRB-Mid (WT) | AWS340 | 400 | 20000 | 44 | 64 | 108 | 0.22 | 0.32 | 0.54 | 59.26 | 32.00 |
| Fig. 3e | pRB-Mid (R106H) | AWS339 | 400 | 20000 | 86 | 69 | 155 | 0.43 | 0.35 | 0.78 | 44.52 | 34.50 |
| Fig. 3e | pCAMBIA (WT) | AWS338 | 200 | 10000 | 81 | 9 | 90 | 0.81 | 0.09 | 0.90 | 10.00 | 9.00 |
| Fig. 3e | pCAMBIA (R106H) | AWS337 | 200 | 10000 | 138 | 9 | 147 | 1.38 | 0.09 | 1.47 | 6.12 | 9.00 |
| Fig. 5a | pRB-Mid (1xLB) | AWS313 | 400 | 20000 | 65 | 94 | 159 | 0.33 | 0.47 | 0.80 | 59.12 | 47.00 |
| Fig. 5a | pRB-Mid (2xLB) | AWS317 | 400 | 20000 | 54 | 76 | 130 | 0.27 | 0.38 | 0.65 | 58.46 | 38.00 |
| Extended Data Fig. 7d | pUBQ10, Divergent | AWS429 | 200 | 10000 | 231 | 16 | 247 | 2.31 | 0.16 | 2.47 | 6.48 | 16.00 |
| Extended Data Fig. 7d | pUBQ10, Reverse | AWS430 | 200 | 10000 | 202 | 28 | 230 | 2.02 | 0.28 | 2.30 | 12.17 | 28.00 |
| Extended Data Fig. 7d | pUBQ10, Forward | AWS433 | 200 | 10000 | 142 | 24 | 166 | 1.42 | 0.24 | 1.66 | 14.46 | 24.00 |
| Extended Data Fig. 7d | pUBQ10, Covergent | AWS434 | 200 | 10000 | 79 | 31 | 110 | 0.79 | 0.31 | 1.10 | 28.18 | 31.00 |
| Extended Data Fig. 7e | RUBY | AWS500 | 300 | 15000 | 183 | 17 | 200 | 1.22 | 0.11 | 1.33 | 8.50 | 11.33 |
| Extended Data Fig. 7e | Linker | AWS501 | 300 | 15000 | 264 | 20 | 284 | 1.76 | 0.13 | 1.89 | 7.04 | 13.33 |
| Extended Data Fig. 7e | Recoded | AWS504 | 300 | 15000 | 222 | 17 | 239 | 1.48 | 0.11 | 1.59 | 7.11 | 11.33 |
| Extended Data Fig. 7e | Intron | AWS553 | 200 | 10000 | 107 | 16 | 123 | 1.07 | 0.16 | 1.23 | 13.01 | 16.00 |
| Extended Data Fig. 7f | pRPS5A, Divergent | AWS427 | 200 | 10000 | 197 | 13 | 210 | 1.97 | 0.13 | 2.10 | 6.19 | 13.00 |
| Extended Data Fig. 7f | pRPS5A, Reverse | AWS428 | 200 | 10000 | 175 | 32 | 207 | 1.75 | 0.32 | 2.07 | 15.46 | 32.00 |
| Extended Data Fig. 7f | pRPS5A, Forward | AWS431 | 200 | 10000 | 157 | 24 | 181 | 1.57 | 0.24 | 1.81 | 13.26 | 24.00 |
| Extended Data Fig. 7f | pRPS5A, Covergent | AWS432 | 200 | 10000 | 151 | 44 | 195 | 1.51 | 0.44 | 1.95 | 22.56 | 44.00 |
| Extended Data Fig. 7g | tHSP | AWS643 | 200 | 10000 | 160 | 10 | 170 | 1.60 | 0.10 | 1.70 | 5.88 | 10.00 |
| Extended Data Fig. 7g | tFAD | AWS644 | 200 | 10000 | 114 | 6 | 120 | 1.14 | 0.06 | 1.20 | 5.00 | 6.00 |
| Extended Data Fig. 7g | tNOS | AWS645 | 200 | 10000 | 134 | 12 | 146 | 1.34 | 0.12 | 1.46 | 8.22 | 12.00 |
| Extended Data Fig. 7g | tOCS | AWS646 | 200 | 10000 | 120 | 26 | 146 | 1.20 | 0.26 | 1.46 | 17.81 | 26.00 |

**Extended Data Table 2:** iPCR sequencing results

| Line | Border | Alignment | Sequence |
|---|---|---|---|
| 2 | LB | Chr3:21578067:21578910 | CGTCGCAGGCACGTCTTACAGTGGTTTACACCACAATATTTCAAATTCAACTTGTAGCATTTGCCATCGACTTAAAACTT<br>TTAAATAAAATATTTTAAAATTAATATATGCAAGTTCTACAAGTTGATTTTGACCACTACATGTTGGTCTACAGAGAAATG<br>AAACTATGAATTTTAATATCCTTTCTTGGAATGAAAGGTTCCTTTTCGTACTCAAATTTATATTATAAAATTTTGATTCAAC<br>AGTTTCATCATAATCACTTACTCTCTGCTTTAATTTTAATTTCGTATTCAAAACCGCAGGTGGATGAAGACAAATTAGCAG<br>AGACCAAAACGTAGTTGAGGATGAAACAATTACTTAGCAGAAAATGAACAGAAACAGAGGAGTCTTACTTATTCACCA<br>AGAAACTCTTATATCACTTAATTTTCTCAAGTTAAAAATAGGATATGTATATGTTTTTTAACTCTACCTAAAAGGAAGATA<br>CTGAAATGAGGAAAGCTGAAATAAAAAGACTAGAAGTAGGAAACACTATTCCTCATTAAACAATGGAGAAGACTCAAA<br>TCAAGAAATCACAACTTCATCTTAAAATCTCAGTCAAACAACATCAATGAAGCCGAAGCTGCAGCTTTCTCTTTCTCCAG<br>GTCGGCTTCAAGTTCTGTCACTGCCAGCTTTCGCTTCTTGACCTGTTTCTCGAGTTCTCCGATCCGTGTGTCGTAAGCTA<br>TCTGTTTTTCCAAAGCCTGATCAAACTTCTGACGCAACCAGTCCAAGTTCAAGCCTGCTTTAATCAGATCCGTAATCGTA<br>TTGTCTGCTTCGTTCATGTCTTCCACTGTGAGTTCCTTAGTAGATTTGGACAGTGTCTTCGTAAGGTCAAGAAGAGCAT<br>GCATAGCATCTTGTTACCGACCAATGCTCAGAACTCACGACTCC |
| 2 | RB | Chr3:21571044:21578058 | AACTTCTCTCAAATACACACCGCGTGCCGTTTAGTGCTCTGCTTATTGGACAACGTGTGGTATAACAGAACGTGAACGC<br>TTAAACATTTTTCAACTGGGCCGGGCTCTTTGGCCCAACTCTATGACTTGAATGATACCATCAGTGAAGTCAGGTCGGT<br>CGGTGTTCGTCGTATCAAAGAAGTTTCCTTTAGAGCTTTTTCTCTCATTCTGTCTTCGTCTAAAAAGGTGAAATCTTTAAC<br>AACTGATCATATTACTCTTAATGATTTCGTAATCTCCTGTTCGTTACGTTGATTATTGTTATCTGTTAAAATTGATGTTTAA<br>GAGTGTCATCGTACTACTGGTAGCTTGATTGCAGATAACAATGGGGAAAGAAGTTGATAACAAGTTCACTTGGGTGAT<br>TAAGAATTTCTCCTCTCAGCAGTCCAGGAAAAATTATTCTGATGAATTCTTCGTCGATGGCTGCAAATGGTAAGAAGTG<br>AATTTTTTCGAATGATTTCATCAATTTAGTTTTTTTTTTGTTTTCCGTGAAATTGAGGATTGACTAAATTTTCTTGAAGGCG<br>TCTTCTGGCCTTTCCTAAAGGAAATGGTGTTGAGAAGTTGTCACTGTATCTGGCCGTTGCTGGTAGTGAATTTTTACCT<br>GATGGATGGAGAAGACACGCATATTTCCACTTCAGTGTAGTGAATCAACTCTCTGATGAACTCTCTCAAGCTAGAGGT<br>GATTGTTTTTCTACACAATGCTCGTTACTTGATATGCGTTGGAAATTGATTGCAACAGTTTTTTTTTTCCTTATGTTGTTTG<br>AGTTGGTTGGTTTAAGACTAGTGTCAATGTTTCGTAGCAAGTTTAGAGCATGGTTCATGGATGGCTTTATTTTATGTGAA<br>GTGAAGTCAATGATGAAATGGAATTTATTTCGTTGTTGTTACAGAAACAAAAAATTGGTTTGATGCGAGTACTAGTGAC<br>TGGGGTTTTACATCAATGCTTTCCCTTAAAAAACTTCATGACAAAGATGGTGGATTCCTGGTTAATGGAGAGCTCAAGA<br>TTGTTGTCGATGTTTCTGTTCTTGAAGTTATTGGCAAATTAGACGTACCAGTGGAATCTGAAGAGACAACCACCAAAGC<br>TCTAAGCGAGCTAGAGGAAAATGACGTACCAGAGGAATCTGAAGAGACAACTAAAGCTCTAAGCAAGGTAGACGAA<br>AACGATGGTGCAGAGTCTAATGATTCGCTCAAGGAAGCTTCATCAGTAAAGGAAAGCATGGATGTTAATGGCTTTCGA<br>GTTCTTCCTTCACAGGTATGAAATCAAGAATGGTGTAGTTGTTTCTTAAAATATCTAACCGACATATGTTCTATTCTGTTT<br>TGTGAATTGTTTCTCAAGTAGCAGAGAGCACACTTGTATATTCAGTAAAAACAAGAAATGAAATTGTAAAAGTTTCTTGA<br>TACATGTATGATGTATCTATGATTTGCAGAATATGAGCGAAACTGGAATATTGGAGATGTTTTTGTCTGTTCTGATTTATT<br>CAAGTCTCTGATAACTGCCTCCTCAATTTGTTTTTTTCTACTTAATTCATCTTTCTATCACTTTCAGGTGGAAACTGTGAGC<br>TGTATATTTGAAAGACACCCCGACATAGCATCAGAGTTCGGTCCAAAGAATCAGCATCTGAGGTCAGCTTACATGAAC<br>GTCCTTCTCAGCCTAATCAAGACTATGTGCCAGTCGACTCAGGAACTTTCCAAGGATGATCTGTCAGACGCAGATGCT<br>GCTCTTGCATACTTGACGGATGCGGGGTTGAATCTAAATTGGTTGGAGGAGAAACTGGAAGAAGTGTCAGAAAAGAA<br>GGAGAACGAGGAAGCTGGTGAGACTCGGGTGCATGAAATTGAAGAAGAGTTGAAGGAATTAAAGCTGAAGTGCTCA<br>AACCTGGAAGCTCAGCTGGAGAAGGAGAAAGCAGATGTGTCTGTGGCCAGAGCTCCTTTCTCCTTCGATGATATTGTT<br>TAATGGCTTCAGTATTCAGATTCCTTCTTATAGTTCTGTTTGGTGTTTTAGTCCTCTTGAAACTCTGCAAGATAAGTGGCT<br>TAGGTTGGATGTTTCCGCATTTTGTGGCTCTTGACTCGTTAGGTTTCTTTAATCCAGTGTGTTTCTTGTAAAGAAGTATTG<br>TGGTTCTAAGACTATTAGTCAGATTTTTTGGTTGTTTCGTTTAGGTGTGTTTCTTTTGAGTAAGGATCATTTTTTCCTTCTC<br>TTTTCTAACTAAACAGATGGGTAACGTTCACATATGGATCAAGTAACACAAAAAAAATCACTTTATCATTGAATTTGACT<br>ATTTCTAAATATGTAAAATTACCTAACAATTACCATTACCAAATCTTTGGGGTTAAAGTATCAAAATATATACTGGGTTCT<br>GTTCATGGTAATTATACGTATGAAGAGGACTAGAGGAGTGTGACGGTACTGCAAACGAGGTCTACAACGGGCAGGG<br>ATTGAACCAAAATTGTAGGAAACCAAAAAACAAAAAATAGAGATAATTTGATTGTTAATCGATTAATTATTAGACCAGAA |
| 5 | LB | Chr2:15943893:15950062 | TACTTTAGCATTCAGTTTCGTAAGACCTTGACCTCCTCTTTAACTTTGGTTATGTAGTATGGTTCTAACTTATATTCTTTGT<br>CTAGAATTGAAATTTACGCTCTCTTCTGTTCAGGGTTTCGGGTTCTTGGCAATATCAGCGTCACCAATACTTTCAGGCAA<br>TGGACAAAAGGGCCTGTTCTACGTAGCGCTAACGGTTATATCCGTTGGTATATTTGGACGGTCTATCTCTTTGGGAGTA<br>TTTACAGAAGACCAACTAGAAGACGGTAGGAACAAGGGAAACCCCGCCAAGCTCGTTAGCTTTGTCATTGGAAATGT<br>GGGGAATTTTGTGTTCTTGTTACTTGCTGCTATTGCAATGCCTCAGATAAGTCCGTGGTTTGTTAGGTTCACGATCCCCT<br>CGGGCTGTGAGGTATTAGCAATGCTGATTTTTATTTCCGGTGCGTGTTCATACAAACGGGTTAAACCAGGTGGGAGCC<br>CTTTGACAACGGTTTTCCGGGTCTTTATGGCTTCTGCTTCCAAAATGTCTTGTGCATACTCGAACAACTCAAGCCAGCTC<br>TATGAAAAAGCTGAGTGTGATCAAGATATTAAGCCCCACACAAGCAGCTTAAGGTTCAATAAAAATTCACTTCATTAAG<br>AGTTGTTAACCCACACTACTGTATAACACTCGTAACTCTTGATTTTTTCAAACACAGGTATCTAGATAGAGCTGCAATGA<br>TACTCCAAACAGAGTCTTTGGAACAACAGAGGAAAAACAGATGGAAGCTTTGTCGGGTGACAGAAGTGGAGCAAAC<br>GAAAAGCGTCATACGCACGGTTCCTTTGTTTGCGACATCTCTCATTTCGGGGATTGTCTTCTCTCTTGGTAATACTTTCTT<br>CCTTGAACAAGCAAACCATATGGACTCCAAGTTCGGATCATGGAACCTTCCTCTCCCACTCCTCTTACTCTTTAGTGAAG<br>CTGCAAGGCTAGGATCCCGAGAACTATGTGTCATGGCTGCAAAACGCCATGCCATTGACTTCCCTGAATCACCTAAAC<br>AAACCAAAACGCCTTATGGTATACCAGTCTCCATCATCCTATCGATATTTTGCTGCTCCATCGCTGCACATGTTGAGTCT<br>AGGAGGCTCAAAGTAGTTAGTACTCAAGGTTTACTACATGAAACTGTCCCCATGAGTGTATTCTGGCTGCTCCCACAGT<br>ACATTCTACTCGGTTCAATCACAGGAATCTACGAAAACAGCTTTGCGCTTTACCTAGAAGAAACGGTGCCTGAAGAGTT<br>GAGCCAATACATGGTTCTATTAAACGTGGGAGTGTGCGGGGTCGGGATCATGAGCAACATTGCTCTTGTTAGTTTGGT<br>GGGCAGTGTAAGCGGAGGCAAATGGTTTCAAGACACAATAAACAAGAGCCGGGTGGATAACTATTATTGGGTTATAA<br>CAGTGTTTTGCATGTTCAATCTCTTGTTATATTTCATTGTAACCTATAGGTACACAGTGTGTAACAAGAAAGATGGTGCA<br>ACACAAGAAAATGATCGAAGGATAATCGCTTCTGTGTAGTACTTGTGTTGTGTGCCCAATGTTTCAATTATACATGTATG<br>AGGTTAGATGAGGTTATTGCGTTTGTGGCTTCGGAAAAACATAGAAGAGAAATTAAAAAGGAATTCTCTCAATGTTTTT<br>GACCAAATCCTCAAATCATCTTCAGGCAACTAAACCATGTTTGCACACACATTAATAAACCAAAGACTAAATGGGACTG<br>GGACATCAGCTCACTTTAAAACTGAACCATTAAAACCTTCAATGACTTTTAGAAACAAATAAAGTAAGTTATGACTTAAA<br>TTGGAGGACAAGATTTAGAAAAACAAATCGAATTTTATGGAAAATAAAGAATTTAGTCCAAAATAGATTTACAAGAATA<br>ATTGGGAATCAACGGTCTTCCTTCAAATTCAAATTTGCAATTACATGGAGATACTTCTCGTATTACTCACAGTCACAAGG<br>CATCAAAATGGAACCCACCGAACATCAAGTTAAAGAACAAATTCACTAAAATTTTGAAAGTTTCCTAAATCTTAAAAAAA<br>GTAACTGCAAACTATTATTCTAGTTAAATAATCAGAAAGTCATTTTAAAGCCTGATACCTCTAAATACCATCATTATCTTT<br>GCCTATGTTCGAACCCCTTTGATGAGTGTTAAAATGTTGCTCTTATGAAAATCACATTATTATTTTAGTATCTATCCAAAG<br>GTTGGAATAAAAAGTTATTTTCTTCTGAATCTAATTGTACTGTGTATGGTTGGAAAGAAGATACAATACAAAAGATGAAC<br>AAAAAAAAAAAAAAAAAAAAATGTTAATTTCCCAAGATGAGGATTTAAAATTGCCAACTTGGTTGAGGAACAAAAGCAT<br>ATCAATCATTGTGGCTTTTTATTTATGGTCTTTCTTTAGACTCTCTGTAGATTCTCATTCTCTTGTCAACTTCGAACTTCAG<br>ACAGTCCTATTAAATAAAACAGTTGGGAAAATAAAAGTAATTGAAAACTAAATTCTAATTACATAAGAAAAAGGGTTTA |

| | | | |
|---|---|---|---|
| 5 | RB | Chr2:15942019:15943852 | CATCTCCCATTGGCTAACAAGATTTAAGGCGGTTATAAGGGTACTTCGGTAATTTGCTATTGGTGTTGACATTTAACCTT<br>TTGGGCACATGAGTTGTTTGTTTGTATATCGAGATACTTTCTCTTTTTTTCCTAAGGGGAAATGATTTTGTGACAATGAAA<br>CGTTTAATACTATATAAATTTGGATCATAATCTCGATAACAAAAAGCATTGGAGAATGTGGTTAAACTAGCATGCTAGG<br>AAACATAAAGGGGCATATCCTATCACAAACTATACTTATCTACGAAAATATGAATTACAAAATAAATAAATAAAATGAAC<br>TACAAAATTTAGTACAATGCCAAAATACGTATAAACTACAAACCCTAACAAAATTTTACAAAAATAGTTGTACTCATCTAA<br>ATCAATATTGATCTAAAAGGCGTTAATTTTTTGCATTGACATGTGACAGAATAAATTGCACAAGAAATGGCAATAAATAT<br>TTTTAGGCGTACTATTTGCTAAACCGACATTAAAGAACTGTCAAACAAATATAATTTTAATATGCAATGATGATTCCTTGC<br>TTAGCTTCTTGGTTCAACGGCCACAGCAAAAAAATTCAACATCAAGGAAGTAATTAAATAGTATAGTAAATGTTTAGTGAT<br>CATAATTGTGTGTCTAAGATAACAAGGTCGTAATAATGGAGTTTAGCGAGAGAGAGGTCGAACCTTAATTATTAATTAA<br>CGTATGAAATATAATCATGATGATATTCTAACAACATTTTTTTTTTGTTGTGTCTGCACATTCTAACAATATCTAAATTCCA<br>AAAACTGGTACTAAGGTTTATTATGAGTTATGACCATAAGAATTTACGTAGCATATATATAAGATGATTTTCCGAAATTTC<br>AAGAATAATTTCTTTATAGGAAATCTTTAGAGTTAAATCAGTTTTTTGTTTTCTTTTTGTAACTCGACTAATTTAATTTATGT<br>ATAGTTTACTACTATAAGATTAATGTATATTCGAAACACATTATGAACGGCAATCACACAAAACGTAAATTAAATACATA |
| 6 | LB | Chr5:22771295:22776455 | GATTCAAATCTCAACCTCTTCTTGTCTTCCAAGTCTGGCTTTAACACTGTTTGTATGCAAACAGTTGCACATGTCTTTGTT<br>TACTTTGTTTTAGATGCTTCTCTCAGTGTCATCAGCATTCAAGTGCGTTTTAAGTCAGTAACAAGATGAATACCTCTTGTT<br>GGCATCTTAAATCACACGAATTGTTCATGTTAAAGAAGAAGTTTCAAATTGTTTGTTGATATTGAAGAGAATGAGAACTA<br>TTTTCTATGTTTACTTTATTTTGCTATTTTCATTTTTTTCTTTCAGTTCCCTACTCTTATCTGTCTTGTGATTCCAAATGCAGA<br>ACACACACAGAATCTCCCATGGCTGCATTTGCAACGGCAGAAGCGTGTGACAGCAACGCAGAACTAATATCAAATGG<br>AGACTTACGCGCTCTTCACCCAATCTTCAAGATCTATGGCCAAAGAAGATGCTTCTCAGGACCAATCGTGACTCTCAAG<br>GTCTTTGAAGACAATGTCCTCGTCAGAAACCAACTAGAAACGAAAGGAGAAGGCGGAGTCTTAGTTATAGACGGTGG<br>TGGAAGCATGAGATGCGCGCTTGTTGGAGGAAACCTCGGACAGTTAGCTCAGAACAACGGGTGGTCGGGGATTGTT<br>GTGAATGGATGCGTTAGAGATGTGGATGAGATCAATGACTGCGATGTTGGGGTCAGGGCATTGGGATCTAACCCGTT<br>GAAATCTACTAAGAAAGGTCATGGTGAGAAGAATGTGCCGGTTCATATTGGAGGAACTTTGATTAGAGATGGAGAGT<br>GGCTATATGCTGATAGTGATGGTATCTTGATCTCCAAGACCGAACTCTCTGTTTGAGTCAGAACGTTTCCGTGATAGTG<br>CTAAGAGACTACAGCTTCTTGGAGCAAGGTGGTTTATTCGAATTCTTAAGAATCTGTTTAATACTCCAAATAATCATTTTT<br>ATTACAATATACTTACGTTTTATGAAATTACAAAATAAAAGTCTATTTAGTTCAACAAGTTTGGATCATGTGATATGTATA<br>AGAAACTTTCATTGAATTGAAAGAGATCAATAAGTCAAAACATGTGTATCAGTGAATACATTATACAACAAGGATATAAT<br>TTCTAAAGATGTTATTTTCGTCTTCCAGCATTGTAATGATTCTTGTACGAATGGATTTTAGTTATTTTGAAATTTAATTGAT<br>AGACATGTATAAATTATAATTGTATACACAAGAGCGTACTATTTTACTTGAATTTGAATACAAAATTCTGCTATGAGGTTT<br>GAAACTTGAAAGAATACATATCGTGGTGGACCAATTACTACTTTTATGGAATTGAAACAAATGAATGAATACTATTTCAA<br>AATTTGACCAAAAAAATAAAAACACCATTTCAAACTAAAACAAAATTCATACCGAACCAAACAAAACAAGATCAGATTTA<br>CTTGTCACGTTAATTTCGGTTCACTCATTTTTCTCGACCAACACACTTCGAAATCGTACGGAAAAATCGGTGCTCCCCGT<br>TAATAAAAAAAAATAAACGGTGTTTCCCAAGAATAATAAAAAGAATAAAAACATATATATTATTCAATGCTAGAGAAACT<br>ATATTATTAATATGTTCCGTCACATTAATAAATAAAAGCATCATAAAAAGTTAGACAGACCACACCATCACCATGTTCAT<br>GCCATTGATAATTTGACGTATATGTGGACATTTCATGTTTTCTCTTTAACACTCCTTTGTTTTTATACATAGGATCTACAAT<br>AAGTCTATTATTTCTAGCATATTTTGTATCTATCGAATCTTCTCAATCAAAAGTTAAATTATATGTTGAAATATGTCTTTGAT<br>CACATATTGATGTTGTAAAATAATTTGGATGATATGACACTCTAGACTAAAATAAGTAAATTCTAGACCATACAAATCTAT<br>AATTGATTCCAATTAGACACGAATATATTATACCACGACCACTTGTTTCTCAATCTAATTTGATGGAACTTATTAATCAAA<br>AGCTAAAAACGGCTGCTCTTTAGATTTATCATTGCGATTAAGAAGCTATAATTTGTTTTTCTTGTTTTGGTGTCTTGCTTG<br>AATCCATTACTTATCAATTTGTCATTCATTACTTCACCAAAAAAAAAAAAAAGAATAGTGTCGTGTAGCGAAGGAATTATG<br>GGACCTCCAATAAGTCCATAGGGCATGGAATATGACTTAACGAGGAAAATGATGTTTGACTAAAACCATAATTTCACGT<br>TTCCACAAAATGGCACTATCCTCTCTTATTTAATTCTATGATTTTTCTTAATTTCTCTTAATAAGGTGATGATTTAACGATAA<br>TTCTGGCTGTTGATACATTCGTTTTAAAAAGATCAAAATATGAGATATCTCATGGTTTATACTTTATAGAACTTGTAATTTT<br>GAGAAAAAGTGGTTATGAGTGCCTACATAATTCTATATGCACTAACATATACATGACCAACTTTTTCAAACAATTGGATC<br>CTTTTTTTCGTAAACATTTGGCTTGCTCTCAAAAACAGTCAAGACATCATGTTCTTATTTATTTTCACCTTTAAAAAGGACA |
| 6 | RB | Chr5:22776467:22778231 | TGAAACGTATTAACCTAATTAAACTGCAATATAAGGAGATACGTTTACCAACCATTGTGAAATTCAGATATTCAAATATA<br>GATTGAATCTAATATATATAACACCTATTAAGTAGTTAAGTACTGGTTAAAGAAACCTAAAATAAAATCATTAAACGAAC<br>TAAATAACATCAGACAAATATTCTTTACAAATATATGAGATACGTTAACGCTTATATCTTTTCTTTTCTTTCTTACATGTTAA<br>CGTTTATAGTCTGAAAAGAGGAAGAAACAAAATGACTAACGTGCGTTAAGGATGATCTAATCTAATTGTATGTAGAGA<br>CATTTAATAAAATTGGGAATAATCAATCAATGACTTTTAATTATACCTTAAATTGAATATTAAATTGGAAATTGAGGAGGC<br>AGTTTTTCCAAAAAAAAACGGTCAAAAAAGAAACATTGACCAAGCCTCTTTTAGTATTTTGTTTTTCGCTTTCACTTTTATT<br>CTTCTTTGTCTTTTTACGAGCCGCAAAATATCCCGTTACGGAGCTGTGACCTTACACTCATGTGACAGCCAAAGCACGG<br>TTGTGGCGTGGTGTACCTTTTTTTTTTTTTTTTTTTCTTCTTACAGATATTTTTATTTTACATCGTGGGGCCCTCAAAACGA<br>ACTTTGAAAGGACGGTGGATCGACGGCTCTGATGTCTTGTCAGTTTTAAGTATTCAATTATCCATGTAAACCGATCCCA<br>TCCGAACCAAAAATCGAACTGTTATCTAGGGGCTACTAGGCTCCCATTTCCACTGTGTAAACTTGGTTTAATTTCTACAT<br>CACTTGGCCCATTAGGCCCATTATAGCTTAATAAAAATAAAATATTTGGAATGGACCCAGCAGACTTGTTTTGACTGTCC<br>GATGATCGAACCAATAGTCAATAGAGAAACATGTCAACGATCCGACCCGAGAAAAGCGGGTCGGATTGAAAATACCC |
| 14 | LB | Chr5:1397972:1398927 | GCTTCTGTTTAAATCTTTTACTACTTTCAGTTTTATATCTCTGATAAACTCTATAATATTGACTGATTCCTAAAGGACACTG<br>TCTCAGGATCTATCCCCTACATTCTTGTTCTACCTTGATGGATCATATATTTTACCTTAGTAAAGAAAATCGTCCAAGAAA<br>TTCATCATTAATTAAAAACATACATATAACATAGTCCTTCTTCGAATATAAGACTTCAATTCCTCAAAAAAAAATTAAACGA<br>AATAAATTATTTTTAAATATATATTCAAGAAACTGATGTTCATCGAGTTTAATTATTCAGTTGAATTACAATTCTTACTTTTA<br>GAAATTTGTATTGCGACCAAGAAGAAACATTCTTTAAGCTCAAAGCTTCATATAATTTACACATACGTAATTTATAGATAA<br>CACGTTGACCGAGAGTATTAGAAGCTCACAGTGCTTATAGGAAAGTCAAAACTGACTGCATCAGTACGATTTTTACGT<br>GTTCGTTTATGGATTATATAATACAAAATAGCTTATACATCAGTTTTAAAAAAATTAAACAAATTAATTAAGATGTAAAAC |
| 14 | RB | Chr5:13576233:13577280 | TTTCTTCTCAATTAGGGTTCTCCCAATTTTCAATTTCATTTCCTTTGATATTAGGGTTCTTTCGAGTTAGTGTACGAATTAG<br>GGATCTTTACTAATTTCAGTTCTATGTATTTCAGATCTTGCTAAGAACTTATATAATTCAACGCTTTCTTTCAAAGGTTTGT<br>CCAAGTCGATAGATTTCTTATGCCTTTGATTTCTTACACAAAAACTTTATTGCGAATGTGATTGTTTTTTGTATCATACATC<br>TTGTGTGTTGCTGTCTGACTTGTTTTCTTTCTTATACATGTTTTGTGTTGCTGTCTGATTCATGCTTGTTTTGGGATACTGT<br>CTCAACGGTTCAAAATAAGGATAACAGATAAAATTAGGCCTCGACATAAAAACCGTACTCGCATACCCGAACCGGAAC<br>CGAACAAAAAAAAACGATTCAGGGCGGGTACGAGTATAAGGATTTTACACTGATGGGTCCTATGTAAGTGTACCCGT<br>GGGTATCGGTTCGGGTTCGGGTATTACCTGATATCCGAAAGGGTAAACCGAACACCCGAACCAAACACATTAACTCAT<br>TAACCACCCTTAACCAGTTAACCCTTATGTCGATTTAGGTTTTTTACTTTTGTTGATTTAGCTTCTTCCCTAACCCCTACTT |

| | | | |
|---|---|---|---|
| 15 | LB | Chr3:982891:986543 | CCTATGTGAATACAACTTGTAATGGTTTTGCTTAAAATGTGAGAACATAACCATGTGAGGAAAATTTTGTTCATGTATGC<br>CCCACACTTTTAGACACCATTGTTTTCAATTGATAAGTCAAGTTTGAAGTTTCAACTCACTCACAGATCTATATATACAGA<br>AGAAACGAAGTCTTTTAACATCATCAACTTACAAGCAATCTAGAATCTCTCTCATTTGTATCCATACATCTTCACAAGCTT<br>CAAGAATCAAACTTCAAGACAAGAAAATGGCTTTGGTGAGAAGTCTCTTTAGCGCAAAGAAGATTCTTGGCGGTTCTTT<br>AGTAAAAACAAGCAAGGCACCGCCAAAAGGGTTTCTTGCAGTGTACGTCGGCGAGAGCCAGAAGAAGCAGAGACAT<br>TTTGTACCAGTCTCATACTTGAACCAGCCTTTGTTTCAAGATCTTCTAAGCAAATGTGAAGAAGAGTTTGGTTTTGATCA<br>TCCGATGGGCGGCTTGACAATCCCTTGTCCTGTAGATACTTTTATCAGTATAACATCTCAGCTCCAAGGATGAAGATGA<br>TGATGATCCAACAAAAATACATTAACAATTTTTTTTTTTTACTCAAACTAGAGATGGAGGAGTATTCCATGTGAATAGATA<br>ACATTTTTTTCTCCTCTTTCCTATTTGATAGAGTTTGTTCTAGACGATTAGAAAATTTCGAATGTGATTGCCGTTTTTGTAT<br>AACAAAGATCAAATCATTTACAAGTTGCGGTAGCTTTTTGCTTGATCCAATTGTTCAACGAGTATGAAACAAACTAATCA<br>AAGATATCACTTGACGACTTAGCAATCTAAATATCAGTTATTGCAAAAATGTAAAATGTCATAAAAGATCTCTTCGTTGC<br>AGCAAATCTTGAAACTGAGATTGGATTAGGAACGAAGATGGAACAAAGAATCTTTTATCTTCTCTATATTTATTTATATT<br>GAAAAACTCAAATGTAAAAATCAATACAAATATCAGATTACACAATCATGTCAGACTCTATATCATCAAGTTTGTTGGTC<br>GACGTCGATCAGTAAACAGAGACAGTTCAAGAGTAAAGAGAATCAAGTTTAAGCAATTTCACCACTACAGAAACATAA<br>TTATGTGACTATGATTAGCGTTATGATTAGGGTTAAGGTTAAGAGTATTATCCGGATCCGTGGCGCAATGGTAGCGCG<br>TCTGACTCCAGATCAGAAGGTTGCGTGTTCGATTCACGTCGGGTTCAAATCCCCCGAACAATAAATTCCGGAATATTTT<br>TTTACGTAGTGTCGGTCGGCACATTCAACATAGCGTTCGCCGTTAGGCTATTTCCAGCTTTGCCCTTTGCGTTTCAATTC<br>AACCTGTAATCTCTTAAGGGGCAGAGATAAGATTTCGACTGTCGATGATCTCTCCTTGAAAGAGTTCTGAAACAGTAAA<br>TTGTGATTTGCTTGAATTCCAAATTCAGTTATTTCTAGAGGACGAACAATCATGCAAGTATTTTTGCAACCATCCTTCTAT<br>GGATGCGGATAACGTAAACTCTAAGTCTCTAAGTCCATTCACGTTCATGGACACGGAGAGAACATCAGAGGTAAATG<br>GGGATGAATGTGCAGAGACAAATTAAGAAGGGCCATAAACGAATGGAAGTTTAATAAACGATGAGATTTATCCCCTTT<br>TTAAGACTTTCATTTAGATAAATCTTACCAAATTGATCCATGTTGGATTAGATTTCATTTTGGCGGGAACTTTTCCATTTTT<br>CAAATCAAACGAAAAAAGGAAGAAAAAAAAAACTGAAAGTGAAATGAAACTAATATTCAACAGTAATTTGCGTGCTTAT<br>GAATATGATATATTTCAGTGTGTATTTGAAAATTTAAAATGGAAAATAACGAAAGGAAGTTACAGAAAAATCATAGTTAA |
| 15 | RB | Chr3:986547:991441 | [illegible]<br>TTGGCGGAAGATCCATACACTATAAAAGTATAAAGAAAGAGCCCTACACTATATAGCAATAATTATCAGAATAGCTTTA<br>ACAAAATAATTATCAAGATATCGAAACATAATTACGTTGTTATCAACCTAGTCCGTTTAATAGATATACGATATTGTTTCC<br>TATTAAATTGGAAATTTCGGTTACAATCCGGATTCAATACACTTTTGACTTTTGATTTGGATTCGGTATAGTTTGGATAAA<br>CTAAAAAGTAAGGGGAATTATTTGTAATAAATAAAAGTAGAAGGATATTAGAAAAAAAATTCAAATTGTAACAAGTCGT<br>CTTTTTCTCTGTTCATAATAAAAAATTAAAAACCCTTGTCTTTCCAACTGTTGACCAAGACAAGCCCTCTCCACTTCAGAT<br>TGTTTCTAAAAGCAATTCTCAATCTTTTCTACACCGAAATAGCAACAAAATGAATTCAATCTCTTCATCTTCCATGCGTCG<br>TAATTGGATACACGATGTCTTCCCGAGCTTCCACGGTGCAGATGTCCGCAAATCGTTTCTGAGTCACACTCTGAAGGA<br>GCTCAGAAGCAAAGGTATCGACTTATTCATTGATAATGATATCTTGAGAGGAGAATTTATCGGCCCTGAGCTCAAAAAA<br>GCGATCCATGGATCGAGGCTCGCGATTGTCTTGCTCTCGAAAAGATACGCTTCTTCGTCGTGGTGTCTTGACGAATTG<br>GCGGAGATTATGAAGTGCAAGGAAGCTTTTGGTCAAACAGTGATGGCCATTTTCTACGAAGTGGATCCAACTGATGTA<br>GAGAAGCAGACCGGAGGTTTTGGGGAAGTCTTCACAGAAACTTGTGAGGGTAAAACAGCTGAGGACATTGAGAAAT<br>GGAGTCAAGCTCTTGCAAAAGTGGCAACTGTCACTAGTTACCTTTCAAGCAGCTGGTTCGTCCCTTTTAATTTCAACTTG<br>CTCTTGAACTTTCTGAACACATCAAACTTTGAAACGATATGCAGCCAAGAACTTGAGTTTGTGATTTTTGGATGACAATT<br>TAATCAGATGTATTCGTCTTTGTTAGGGATACTGAAACAGAAATGATCGAAGAAATTGCCACTGATGTTTCGAATATGTT<br>GACTAAGTCAACGCAATCAAGGGATTTCGTCGGCATAATTGGAATGGGATCTCATATGGAGAAGATGAAACCGTTGCT<br>ATGCCTAGAGTCAGAGGAAGTGAGGATGATAGGGATCTGGGGTCCTCCTGGAATTGGCAAGACCACCATTGCCAGA<br>TTTCTTTTCAACCAACTATTTGGCAATTTTGGTTTGATTGTCTTTATGGATAATATCAAAGCAATGTATACAATACCAATTG<br>GCTCAGATGACTATAGTGCAAAGTTGGATTTGCAGCAGAAGTTCATGTCTCAAATTATCAACGAAAAGGATATCAAGAT<br>TCCTCATTTAGGGGTTGCACAAGAAAGGCTCAAGGACGAGAAAGTTCTTGTTGTTCTTGATGGCGTTGATCAGTTATTT<br>CAATTGGAAGCCATGGCAAAAGAAACTCGGTGGTTCGGTCATGGAAGTCGTATTATCATTACAACACCAGATCTAAAA<br>CTTTTGAAGGCACATGGGATCACCCACATTTATAAGGTGGATTTTCCACCACCTGCTGAGGCTCTTGAAATCTTTTGTAT<br>GTATGCTTTCGAGCAAAAATCCCCTAAAGAGGGTTTCGAGATGCTTGCCTGGGAAGCTACAATTCTTGCTTGTCAACTC<br>CCTTTGGGACTAAGGGTTATAGGCTCTTATTTTCGAGGAAAGTACTCGAAGAAGATGTGGGAAGAGGCACTACCAGA<br>GTTAAGGAGTCACCTTGACGGAGAAATTGAAACCGTTTTAAAGTTCAACTACGATGCCTTATCGGAGAAAGATAAATC<br>GTTATTCCTTCATCTAACTTGCTTTTTCAACTTTGGAAGTATTGAGAATATAGAAGTTTATCTTGCCAAGACGTTCTCGGA<br>CTTCAGGCAACGGCTTGATCTCTTAGTTGAGAGATTTCTCATATCCATAGTCTCTGGATATTTAGAGATAAATCATTTGC<br>TAACCCTACTGGGAAGAGAAATTGTGCGAAAAAAATTATGGTAGTGAGGTCTCCATTCGTGAGCCTGGGAAACGCCA<br>GTTTTTGGTTGCTGCAGGAGATATTTGTCAAGTACTGAGTGATGATACAGCAGTAAGTTTTACTACTAATATTTCGTTGC<br>ATTGCTCCCTACAAGCTGAGTTTAATTGATAACTAACAGTTTATTTGATATATTGTATTGTATCCTTTTCAGGGTAGCAGC<br>ACTGTTATAGGAATAGATCTCACATTACCTGAGAAGGCCGACGAGGACGAAGAATTTTATATAAGTGAAAGAGCATTT<br>GAAAATATGACTAACCTCCAATTCTTAAAAATTAGTGGTGATTGCAGTAGATTGTACTTTCCGCCACGTCTGAACTCCAT |
| 18 | LB | Chr3:2642437:2646454 | TTCGTCGCAGGCACGTCTTACAGTGGTTTACACCACAATATATCCTGCCAAGTACGAGTCAACTGCGAGAAGCTGGAC<br>ACAAAAGTACGCAATGGGTTGATGATATAAGACAAAGCAAAGGATATGTTATTGCGCATCGGAATAAAATAATAACTT<br>AAAAAAAATGCACTTGTTTGGTTTATCAAACGTTAGCAAGTATCGAACAACTTAGCTGCTCTTACTCCTCTTCTGTTCTGA<br>TCACTATAAACTAGTATTAATTTCGATTGTTGGGTCTACTTTTATTCTCTTGTTTCCCTCTCCTCACTTGGTTGCAGGAAAC<br>TGCAATTCTAATCGCCCCAAACGCAATGTTTGGACTGTTTTGGTACACTCTTGACATAAGGGGTTCCAATCTACACATG<br>GATCAACCACAACCAAAACCAAACCAACTATGAGCCTTCCATAGTTAGGATTATACATTGCTCCCAAATACAAAACTCG<br>TTCCTAATCTTCTATTAACTTATACCAAAGTTGGTATATTTATGATTAAATTATTGATTTTCTTAATTTTTTTAATATATTTCA<br>AAAGTGTACATGGTTCTAATAAAACACTTAAGAAATTTTCCATATTGTTTACAATAGCTGTTTTAGGAAATATCATATCTA<br>TAGACCAAACCAAGAATATCTTTAGACCAAACCAAGAATAGGTGTTTTAGAAAATAAACATATCTTTGGTTGCAATTTTC<br>CAATATCATCCGCCATGGGTCCCTACTAATAGGTCAGGCGGTACTTGCCGACAGAATCAGAAGCCTCGAGACTGACTT<br>GTATTCTCCGACTATAAAAAAGAAGTAGAGGGTTTTAGGCTTTTCACGAATCTCTTCTCTCTTTTGTAGTGGCCTTGAAA<br>AAAAGATTCAATCTCCTGTTGTGGTATGTTTCTTTTGATCCATCTTTCCTTTATGTTTTCTGTTTCAACTACTTCTTAGGGTT<br>TATCATCACTTACGTTACGTATGTATTGTTTTATTCGTCTTCCTCTATGATATGTTATGTTCTTAGTTGAATTCCTTAATACT<br>CATTGTGTGTTGCTTATCTCGTCCGCAGGTTTTCGAGTCTAAAAGCATGGCTTCTAAGAGGATTTCAAGGGAACTGAG<br>GGATATGCAGAGACATCCTCCAGCAAACTGTAGCGCAGGTAATCTCCCTTTTCTTGGCTGATTTTGACTGATAGATTCA<br>CACTAAAAACACTTTTAGGGTTTGATATATGTCACACTTTTATTTCATCAATCCACATTATATGTTTCTGAGTTTTTTTTGGT<br>AACGCTATGATTTGTTAGACTCTCTTTGGTTCTGTTATATATATGTACTCTGCATGAACCTTTTTTTTTTTTTTTTTGTATTC<br>GAGAAAACTCGTTAAACCTTTGCCTGGTACGATTAAGAATATCCAGAGGTTTAGCTTGCTTTTATTTCAAGTGTCTCCTT<br>TTGTACTTCTTATTTATTCTGCCAATTTGTCTCTGTTCTATAAGAAGTATCACTTGATCATGCAGGTTTTATTGTCTTTTTCG<br>ATGAATAAATGGGTCTGATTGCAACGTCTTTCTTATAGGTCCTGTGGCTGAGGAGGACATATTCCATTGGCAAGCAACT<br>ATCATGGGACCGCATGACAGTCCGTATTCCGGAGGCGTATTTACGGTTTCTATTGATTTCTCTTCGGATTATCCTTTCAA<br>GCCACCAAAGGTGTTGTGTTTTTTCAACACTAATTTAATCATCTGATGTGTAATGAAAATCTATGATCATCAGGCTTATTA<br>GCGAGTGTACGATTTTGCAGGTGAATTTCAAGACTAAGGTGTACCACCCAAACATCGACAGCAAAGGAAGCATTTGCC<br>TTGACATATTGAAAGAGCAATGGAGTCCTGCTCCTACCACATCCAAGGTTTGAGTTTTGTCTCTCTACTACTATTTTTTAC<br>CAAAGTCATGTTCTGTAAAAAAAAAAACTAGTATATTGGTGAACTTTGTGCGAGTGATATTGATGGTGTGGAATTATATA<br>GGTTTTGTTGTCGATTTGCTCGCTGCTGACTGACCCGAACCCAAATGATCCTCTTGTGCCGGAGATAGCTCATCTCTAC |

| | | | |
|---|---|---|---|
| 18 | RB | Chr3:2636403:2642318 | TGAAAGAGGACGTCAAAAACACAAACTGGTTACATAGCATCGATGACCGGTCGTCAGATCACGGACTCATGTTTGACA<br>ATGGAGGATTCGAGCTACTTCCAGAGCCATTTGCCCCACTATAACACACACACACACACACCCCACAAGTAAACACCTT<br>TTTTTCTTTCTAAAATCACTTCTCTTTTTTTTTTTTTTTTTTTAAATTTTATCTTACACAATTTGTTTGCTTGTCGATGGTGG<br>AAAGATATAAGGCTCAAGTGGTTCTTGGTTCTTTGGTTATTGATTCTTGATTCATGGCTATTGAGTGAGGTTTGTAACTT<br>TGTAGTTATAATATTTGAAGAGATGGTAACGTATAAGAGGGTACCTCATGTTCACTCTTGTTGGAAACGATCATGAACG<br>AAAAAACAATTAGAATGTGAAAAGAGGAAAGAGAGGGTCACACGACTCACGCGCCTAACGTGAATCAATGACAGAG<br>CTAAAGTTAATAATGGGTGACAACGAAGCACACAACCAACAACAGGCCAGTGTAGTTCTGATAAAAGAAAAAAATTAA<br>TTCCAGAAATTGACTTTCATTAATGTCATATTTGAGATTTAAGAAGTTTTAAATTTTTGTTATTTATTTTCTCTTTTATGTTAC<br>CCCAAAATAAATAAATACCCTCATTAATGTCACATACCAAACTCCTCCTCTTCAGGGGGTCCATATTTAAGAAAGATACC<br>CTCTTACCCGAATTAAAAATATCAAATAAAGTAAGCTCAATTTGCAACATATAATAATTTGTTGAATATTCCCCACTCTTG<br>TTGTTTTCCTTGCCCATTTTTTGATCTTGAAATACTTTTATCTTCTTATTGTTTAACGAATATCACAATCATCCATTTTCATTT<br>ATACATTGCATATTTTCAAAAAAGAAGCCATATATTTATTTAATCAACTGGTGACCCATTCAATAGAACATTACGATGCTT<br>TTAAAAGAATAAGGAAATTAAATGTTGGGAATTCAGTAAATGCCGTTTATTAAAAGCTCTTATGTATAAATTCATTTAGA<br>ATAAAAACAAAGAGAAATTAACAAAAGGACAAATATTTAATTGAGCTTAGCAATAAAAAAAAAATCTAAAAAATTCGGA<br>AGAGAGTCAAAGGGGGACTTAAAGCCAAAGAGAGCAAAGCAAATCGGAGTACAGAGAAGAAGAAGAGAGATTGGA<br>ATCTGAAACTGTAACATCGAGATTTACGTTTCTCCGTCGGCGTCACGCCGAAGCTTCCATACTCCAGAAACCCCAAAAC<br>GCTTATTTCTCAAAAATCTCCACTCTCTCTCTTCTGAATCTTCAATCACTTGGGGTTGTGTCACTGAATCTGCATCAATGG<br>CGATTCCAGATTCCTCACTCTTTTCTGCTTTTGACTTTAGCTCTAGCTTCTTCTTCACGAAATGCCGTTGTTGTTAAAAAGT<br>GATTTCTCCACTATCCCATTTCCTTTGGCATTTATTCTCCACTCTCTCTCACTTGTCTTCTTCTTCTTCTACGATCAGCTCAC<br>CAATGTCCTGAACCATTCTCCTCTGAATCAACCTCTAAGGTCTCTCTCACTTTCGATCTCTGTTTTCCCTCTTGTTTTGGG<br>GTTTATTAGCTTCGTTGTTTATTCCTTTAAAAGATTTCTTCTTTCTCAATTTCAATTACTCCGTTTAGCTTAGCTGGAGAAA<br>CCCTAGTTTGTAGATTGCAGATCTGGGTTAATCAAGTATCTGTGCACACATTTATGTCTAAAATTAGTTACTTTCGTAATT<br>GGATTTGTTTTTTTTGTGAAGATTTATTATTTGAAGTCAAACAGATTTCAGCTGATTGAATCAATGTGTGCATATAGCTTC<br>CATAATTTCATATTTTGATTTGAGAATCGAGTAATGTTGCTTTGGACTTTATCATCAGCATTCCCACTTTGGTTTGAGAAA<br>TTACGCTATTGTATCTTTGTCATTATTAAGGAAGCTTTTCGTTGTGTTGATGATCTTTTTACTCATGATACCTGTGATGTGT<br>AGGTGTCATTCTTGGCTACTTTATCCATAGATACATTTCTCATATTGAGTTTAGCTTTTATGTGGCTATAAATCATTACATG<br>ATGTGTTTTGTGTTCTCTTGCAGTTGCTGTTGAATGAGAAAGCTTTGATGTGAGCGAGCCTGAGAGTGTAACAAAGGA<br>CTGATGATGAAGATTATTGCGGCGTTTCTCTTCCTCTTGGTCACGACTTTTGTTTCCCGTTGTTTGTCTGCAGACATAGA<br>ATCGGACAAGCAAGCACTGCTTGAGTTTGCATCTCTTGTTCCTCACAGTCGCAAACTCAACTGGAACTCCACTATTCCA<br>ATCTGTGCCTCTTGGACTGGTATCACCTGCTCCAAGAACAACGCCCGTGTAACCGCACTCCGTTTGCCTGGATCTGGA<br>CTTTATGGACCTTTACCAGAGAAAACATTTGAGAAGCTTGATGCCCTCAGGATCATTAGCCTTAGATCCAACCACCTTC<br>AAGGGAACATTCCATCTGTCATTCTTTCACTTCCTTTTATCCGATCACTATACTTTCATGAAAACAACTTTTCTGGTACTAT |
| 19 | LB | Backbone | AAGGAGTCGTGAGTTCTGAGCATTGGTCGGTAACAAGATGCTATGCATTTCTTGTCTCCCCTATCCCCCCGTTACTTTG<br>GACTGGTTCCCATGCGTTACCGTAGACCGATCCACCTTACACTTCGCTTTACAATAGCCTCGTGAATAATCAGGGTGAC<br>TCAGACACAGTAGGAGACTTACGCCAGACGTGACAGGATATATTGGCGGGTAAACTAAGTCGCTGTATGTGTTTGTTT<br>GCAGCCGTGCGGCTGCATGAAATCCTGGCCGGTTTGTCTGATGCCAAGCTGGCGGCCTGGCCGGCCAGCTTGGCCG<br>CTGAAGAAACCGAGCGCCGCCGTCTAAAAAGGTGATGTGTATTTGAGTAAAACAGCTTGCGTCATGCGGTCGCTGCG<br>TATATGATGCGATGAGTAAATAAACAAATACGCAAGGGGAACGCATGAAGGTTATCGCTGTACTTAACCAGAAAGGC<br>GGGTCAGGCAAGACGACCATCGCAACCCATCTAGCCCGCGCCCTGCAACTCGCCGGGGCCGATGTTCTGTTAGTCG<br>ATTCCGATCCCCAGGGCAGTGCCCGCGATTGGGCGGCCGTGCGGGAAGATCAACCGCTAACCGTTGTCGGCATCGA<br>CCGCCCGACGATTGACCGCGACGTGAAGGCCATCGGCCGGCGCGACTTCGTAGTGATCGACGGAGCGCCCCAGGC<br>GGCGGACTTGGCTGTGTCCGCGATCAAGGCAGCCGACTTCGTGCTGATTCCGGTGCAGCCAAGCCCTTACGACATAT<br>GGGCCACCGCCGACCTGGTGGAGCTGGTTAAGCAGCGCATTGAGGTCACGGATGGAAGGCTACAAGCGGCCTTTGT<br>CGTGTCGCGGGCGATCAAAGGCACGCGCATCGGCGGTGAGGTTGCCGAGGCGCTGGCCGGGTACGAGCTGCCCAT<br>TCTTGAGTCCCGTATCACGCAGCGCGTGAGCTACCCAGGCACTGCCGCCGCCGGCACAACCGTTCTTGAATCAGAAC<br>CCGAGGGCGACGCTGCCCGCGAGGTCCAGGCGCTGGCCGCTGAAATTAAATCAAAACTCATTTGAGTTAATGAGGTA<br>AAGAGAAAATGAGCAAAAGCACAAACACGCTAAGTGCCGGCCGTCCGAGCGCACGCAGCAGCAAGGCTGCAACGTT<br>GGCCAGCCTGGCAGACACGCCAGCCATGAAGCGGGTCAACTTTCAGTTGCCGGCGGAGGATCACACCAAGCTGAAG<br>ATGTACGCGGTACGCCAAGGCAAGACCATTACCGAGCTGCTATCTGAATACATCGCGCAGCTACCAGAGTAAATGAG<br>CAAATGAATAAATGAGTAGATGAATTTTAGCGGCTAAAGGAGGCGGCATGGAAAATCAAGAACAACCAGGCACCGAC<br>GCCGTGGAATGCCCCATGTGTGGAGGAACGGGCGGTTGGCCAGGCGTAAGCGGCTGGGTTGTCTGCCGGCCCTGC<br>AATGGCACTGGAACCCCCAAGCCCGAGGAATCGGCGTGAGCGGTCGCAAACCATCCGGCCCGGTACAAATCGGCG<br>CGGCGCTGGGTGATGACCTGGTGGAGAAGTTGAAGGCCGCGCAGGCCGCCCAGCGGCAACGCATCGAGGCAGAA<br>GCACGCCCCGGTGAATCGTGGCAACCGGCCGCTGATCGAATCCGCAAAGAATCCCGGCAACCGCCGGCAGCCGGT<br>GCGTCGTCGATTAGGAAGCCGCCCAAGGGCGACGAGCAACCAGATTTTTTCGTTCCGATGCTCTATGACGTGGGCAC<br>CCGCGATAGTCGCAGCATCATGGACGTGGCCGTTTTCCGTCTGTCGAAGCGTGACCGACGAGCTGGCGAGGTGATC<br>CGCTACGAGCTTCCAGACGGGCACGTAGAGGTTTCCGCAGGGCCGGCCGGCATGGCCAGTGTGTGGGATTACGAC<br>CTGGTACTGATGGCGGTTTCCCATCTAACCGAATCCATGAACCGATACCGGGAAGGGAAGGGAGACAAGCCCGGCC<br>GCGTGTTCCGTCCACACGTTGCGGACGTACTCAAGTTCTGCCGGCGAGCCGATGGCGGAAAGCAGAAAGACGACCT<br>GGTAGAAACCTGCATTCGGTTAAACACCACGCACGTTGCCATGCAGCGTACGAAGAAGGCCAAGAACGGCCGCCTG<br>GTGACGGTATCCGAGGGTGAAGCCTTGATTAGCCGCTACAAGATCGTAAAGAGCGAAACCGGGCGGCCGGAGTACA<br>TCGAGATCGAGCTAGCTGATTGGATGTACCGCGAGATCACAGAAGGCAAGAACCCGGACGTGCTGACGGTTCACCC<br>CGATTACTTTTTGATCGATCCCGGCATCGGCCGTTTTCTCTACCGCCTGGCACGCCGCGCCGCAGGCAAGGCAGAAG |
| 19 | RB | Chr1:4946719:4948952 | AGCCACATTCTCCACTATCTATACGCTACATACCAGAGTTACAAAATCATTTGAATAGAGGATTAAACTTGGGGTGAGA<br>AGCATGTAATTTGATCAAGACCATTGATTCCTTCAGGCATTTTAAAAGCTATATAACCAACTGCACCCAAATGATGCAAA<br>GGAGCTCACCAAAAATACGATCTTGCACAACATTCACTACATTCTCAATCTATCCTCTCTTTCAAGTGTTCTTTTGATCAC<br>AATATACAAAAGCTAAGCAAGACAATACTTTGATCAAGATTCAAGACAAATTCACAAACTTAAGCTGGAGCTAACTACG<br>CCAAATCTTAGAGATAACAAATGGAAAGAAACTAAGGATCACTTTGCAATAAAAACATTTCACAACATTCTCAGGTTTAA<br>GCGAGTTATACAAAAACAAAAGAAATCAGTTTTATATGTAAGCATAAAAACTAAGCACTCATACACTAACTAAACGCAA<br>AGATCAATTTAACATAGACGAGAAACTACATCTCTACGTTCCAATCAAACGAAGAAAAATTCAAAATTAGCGACACAAA<br>ATGAATCAATCATCAAAGATTTACCTCATCGAAGAAGAAAGATGCTCTTCAGTGAGACGGAGCAGTCTGTTCCTGGTG<br>GTGGTGAGATGATTTGGAAGGAGCCAAAGCTTTGTTGTAGATCTGCTCACCGACGAGATAAACGCAGAAGGCGGCG<br>ACGCCGATTCCAAGTCCGGGAAGAGCGTGTCTCATTTGATTCGAAAGCATCGGATGCTTCCTCCATTCATCTCTTCGCC<br>GGAAAAATTCGCCGGTTGTTCCCAAAGGCTTCGCCATTTTCTCAGGTATTCGATTAATCTCCTGCGACCGTACTATTCCT<br>CTTAACTTTCTTCTCAAATGACAAATCATGGGCTTCTTTGATGTATATGGGCTTTAAATTTATTTTTTGTCGTCATATGGGC<br>TTTAACTTTAGGTTATCATTTGGGCTTCTTTTGAAAAAAAATCGGAAAATAATAATGTCGTAAACTAGTTTTCTTATGGTT<br>CTCTTTAATATCAAAATATAATGTCGGAAATAAGTAATTAAGTTTAATAATATTGCATAGCATCATATATAAATTATATCAG<br>AAATCTATCATTGTAGAGTTGTTATTATTTATTTCTTGTTAAACAGAAAATAAATACAAATGGTAAATCATAGTGAAACAT |
| 21 | LB | Chr3:19877281:19877456 | CTGAGTAGTTTGTTCTTATTGCAGCGTACTACCGTGGAGCCATGGGGATTTTGCTTGTGTATGATGTGACTGATGAATC<br>ATCTTTCAACAGTAAGTACTCTCTTTACTTTTTACTTGCAGTCCTTATGCCACCCAACAAGCTCTTGTAAACCACTGTAAG |
| 21 | RB | Chr3:19876141:19877280 | ATTTGTTTGAATAAAAGGTTTCAATTCTTGCAATGAGTGATAAAACATTAATAGTTGAAAGTTTGAAACAGAACCATTCT<br>CGAATGTATACATAGAATCCAATATCATTGGTGTGTGAGTGAGATAAACTGATATTATCATAAGTCATAATCTGATGTCA<br>GAGAATCCATTCCAGAAATATGGAATACGAATTGTGTCCTTCACAGTCACACTATACCATTAGGGAAAGAGAGAATCTT<br>AAAAGAATACGTAGAAACACCATCTCATATAGAAGAAAAGCAAAAGAAGATTGATTTCCTTTTTCATTTTTTCTTTCCAAA<br>TACCATTTCTCTTTCACTAGCCAGATTCTCTCCTTTGACAGGCGTCGTGGACCCAATTTTTATGTGCCGCAACATGCTGA<br>TTTCTGAGTAGCTTGAGATGTTCCCGCACCTTGGTCGGATTGGTTGATTTTGATTGTTTGCGGCTACAAAAATGATAAG<br>AAATTCGATGACGTTAGTAAGGTTGATACACATACTGAAATGTGATTTTTACCACAATTGCAATGTATGAATGATGACA<br>GTGGAAATTAATTCTATAGCTCAAACGAACCTCAGCTCGTGCATCGGTATCTGCAAGTCTTTGCTTAATGTCTTTAGCAA<br>TAGAGAAGAAAACTTCCTCAACGTTTAAGTTAGTCTTGGCACTCTGCAATTAGCAAACTCATCAAAGTTAACCATTTAAA |

| |
|---|
| Map to same locus |
| Map to different locus/ backbone |

**Supplementary Table 1:** Plasmids used in this study

| Plasmid | Figure | Description(s) |
|---|---|---|
| **pWS9500** | Fig. 1c,d; Fig 1e-i; Fig. 2a-c; Extended Data Fig. 7d,e,g | Divergent; Wild type; Original; tHSP |
| **pWS9501** | Fig. 1c,d, Extended Data Fig. 7d | Reverse |
| **pWS9504** | Fig. 1c,d, Extended Data Fig. 7d | Forward |
| **pWS9505** | Fig. 1c,d, Extended Data Fig. 7d | Convergent |
| **pWS9507** | Fig. 2a-c | Border Var 1 |
| **pWS9508** | Fig. 2a-c | Border Var 2 |
| **pWS9509** | Fig. 2a-c | Border Var 3 |
| **pWS9510** | Fig. 2a-c | Border Var 4 |
| **pWS9993** | Fig. 2a-c | Border Var 5 |
| **pWS9512** | Fig. 2a-c | Border Var 6 |
| **pWS9966** | Fig. 2d-i | pRB-Max |
| **pWS9967** | Fig. 2d-i; Fig. 3e,f; Fig. 5a | pRB-Mid; pRB-Mid (WT); pRB-Mid (1xLB) |
| **pWS9968** | Fig. 2d-i | pRB-Min |
| **pWS9925** | Fig. 3c,d | pRB-Mid (WT) |
| **pWS9928** | Fig. 3c,d | pRB-Mid (R106H) |
| **pWS8872** | Fig. 3c,d | pCAMBIA (WT) |
| **pWS8871** | Fig. 3c,d | pCAMBIA (R106H) |
| **pWS10045** | Fig. 3e,f | pRB-Mid (R106H) |
| **pWS8883** | Fig. 3e,f | pCAMBIA (WT) |
| **pWS8882** | Fig. 3e,f | pCAMBIA (R106H) |
| **pWS9474** | Fig. 4a,b | 35S |
| **pWS9475** | Fig. 4a | pACT2 |
| **pWS9476** | Fig. 4a | pACT8 |
| **pWS9478** | Fig. 4a | pAGP15 |
| **pWS9479** | Fig. 4a | pEF1A2 |
| **pWS9480** | Fig. 4a | pEF1A3 |
| **pWS9481** | Fig. 4a | pEF1A4 |
| **pWS9482** | Fig. 4a | pLEA26 |
| **pWS9483** | Fig. 4a | pPLDA1 |
| **pWS9484** | Fig. 4a | pTUB6 |
| **pWS9485** | Fig. 4a | pUBQ1 |
| **pWS9486** | Fig. 4a | pUBQ4 |
| **pWS9487** | Fig. 4a | pUBQ11 |
| **pWS9605** | Fig. 4a,b | nos |
| **pWS9696** | Fig. 4a,b | Negative control |
| **pWS9682** | Fig. 4b | t35S |
| **pWS9683** | Fig. 4b | tAGP15 |
| **pWS9684** | Fig. 4b | tAPX1 |
| **pWS9685** | Fig. 4b | tEF1A1 |
| **pWS9686** | Fig. 4b | tEF1A2 |
| **pWS9687** | Fig. 4b | tFAD2 |
| **pWS9688** | Fig. 4b | tGADPH |
| **pWS9689** | Fig. 4b | tUBQ4 |
| **pWS9690** | Fig. 4b | tUBQ10 |
| **pWS9691** | Fig. 4b | tUBQ11 |
| **pWS10032** | Fig. 4c | p35S |
| **pWS10033** | Fig. 4c | pEF1A3 |
| **pWS10068** | Fig. 4c | pUBQ10:RUBY positive control |
| **pWS10034** | Fig. 4c | pEC:RUBY negative control |
| **pWS9994** | Fig. 4d | p35S:bar |
| **pWS9995** | Fig. 4d | pEF1A3:bar |
| **pWS9515** | Fig. 5a | pRB-Mid, 1xLB |
| **pWS9519** | Fig. 5a | pRB-Mid, 2xLB |
| **pWS10146** | Fig. 5b | pRB-Mid-2LB |
| **pWS9498** | Extended Data Fig. 7f | Divergent |
| **pWS9499** | Extended Data Fig. 7f | Reverse |
| **pWS9502** | Extended Data Fig. 7f | Forward |
| **pWS9503** | Extended Data Fig. 7f | Convergent |
| **pWS9473** | Extended Data Fig. 7e | Recoded |
| **pWS9464** | Extended Data Fig. 7e | Linker |
| **pWS9838** | Extended Data Fig. 7e | Intron |
| **pWS10143** | Extended Data Fig. 7g | tFAD2 |
| **pWS10144** | Extended Data Fig. 7g | Tnos |
| **pWS10145** | Extended Data Fig. 7g | Tocs |
| **pWS10069** | Extended Data Fig. 13 | p35S |
| **pWS10070** | Extended Data Fig. 13 | pEF1A3 |
| **pWS10071** | Extended Data Fig. 13 | Pnos |
| **pWS10072** | Extended Data Fig. 13 | No promoter |
| **pWS9997** | Extended Data Fig. 14 | pEC:StayGold-NLS:tHSP |
| **pWS10150** | Extended Data Fig. 15 | pEF1A3:nptII:tUBQ11 |

**Supplementary Table 2:** T-DNA plasmid toolkit

| Plasmid | Description |
|---|---|
| **pWS10205** | Plant Entry Vector |
| **pWS10206** | Expansion module |
| **pWS10207** | 1F assembly cassette |
| **pWS10208** | 2F-T assembly cassette |
| **pWS10209** | 2F assembly cassette |
| **pWS10210** | 3F-T assembly cassette |
| **pWS10211** | 3F assembly cassette |
| **pWS10212** | 4F-T assembly cassette |
| **pWS10213** | 4F assembly cassette |
| **pWS10214** | 5F-T assembly cassette |
| **pWS10215** | 5F assembly cassette |
| **pWS10216** | 6F-T assembly cassette |
| **pWS10217** | 6F assembly cassette |
| **pWS10218** | 7F-T assembly cassette |
| **pWS10219** | 1R assembly cassette |
| **pWS10220** | 2R-T assembly cassette |
| **pWS10221** | 2R assembly cassette |
| **pWS10222** | 3R-T assembly cassette |
| **pWS10223** | 3R assembly cassette |
| **pWS10224** | 4R-T assembly cassette |
| **pWS10225** | 4R assembly cassette |
| **pWS10226** | 5R-T assembly cassette |
| **pWS10227** | 5R assembly cassette |
| **pWS10228** | 6R-T assembly cassette |
| **pWS10229** | 6R assembly cassette |
| **pWS10230** | 7R-T assembly cassette |
| **pWS10231** | 1 assembly spacer |
| **pWS10232** | 2-T assembly spacer |
| **pWS10233** | 2 assembly spacer |
| **pWS10234** | 3-T assembly spacer |
| **pWS10235** | 3 assembly spacer |
| **pWS10236** | 4-T assembly spacer |
| **pWS10237** | 4 assembly spacer |
| **pWS10238** | 5-T assembly spacer |
| **pWS10239** | 5 assembly spacer |
| **pWS10240** | 6-T assembly spacer |
| **pWS10241** | 6 assembly spacer |
| **pWS10242** | 7-T assembly spacer |
| **pWS10330** | pRB-Mid-2LB |
| **pWS10331** | pRB-Mid-2LB bar |
| **pWS10332** | pRB-Mid-2LB nptII |
| **pWS10333** | pRB-Mid-2LB hpt |

**Supplementary Table 3:** Arabidopsis promoter design

| Name | Gene | Intergenic | Orientation | Other notes | Acceptable | Promoter + 5'UTR | Final notes |
|---|---|---|---|---|---|---|---|
| ACT2 | AT3G18780 | Yes | | | Yes | 1300 | Tested |
| ACT8 | AT1G49240 | No | Tandem | | Yes | 1040 | Tested |
| AGP15 | AT5G11740 | Yes | | | Yes | 658 | Tested |
| APX1 | AT1G07890 | No | Divergent | | No | | |
| ARF1 | AT1G23490 | No | Tandem | | Yes | 759 | Did not test, synthesis failure |
| AVP1 | AT1G15690 | No | Divergent | | No | | |
| EF1A1 | AT5G60390 | No | Divergent | | No | | |
| EF1A2 | AT1G07940 | No | Tandem | | Yes | 1000 | Tested |
| EF1A3 | AT1G07920 | No | Tandem | | Yes | 1000 | Tested |
| EF1A4 | AT1G07930 | No | Tandem | | Yes | 966 | Tested |
| GADPH | AT1G13440 | No | Divergent | | No | | |
| HTR5 | AT4G40040 | No | Divergent | | No | | |
| LEA26 | AT2G44060 | No | Tandem | | Yes | 827 | Tested |
| PCC1 | AT3G13920 | No | Divergent | | No | | |
| PLDA1 | AT3G15730 | Yes | | | Yes | 1163 | Tested |
| SHM4 | AT4G13930 | No | Divergent | | No | | |
| TPI | AT3G55440 | No | Divergent | | No | | |
| TUB2 | AT5G62690 | No | Divergent | | No | | |
| TUB6 | AT5G12250 | Yes | | | Yes | 1100 | Tested |
| UBC10 | AT5G53300 | No | Divergent | | No | | |
| UBC9 | AT4G27960 | No | Tandem | Alternative splicing in 5' UTR | No | | |
| UBQ1 | AT3G52590 | No | Tandem | | Yes | 683 | Tested |
| UBQ10 | AT4G05320 | No | Divergent | | No | | |
| UBQ11 | AT4G05050 | No | Tandem | | Yes | 720 | Tested |
| UBQ4 | AT5G20620 | No | Tandem | | Yes | 790 | Tested |

Selected for charatcerisation

Promoter selection criteria
1. Not in divergent orientation with neighbouring gene, including lncRNA (within 2 kb)
2. No 5' UTR overlap with neigbouring gene body
3. No alternative splicing in 5' UTR

Promoter region defined by open chromatin in leaf ATAC-seq data from Lu, Z. et al. The prevalence, evolution and chromatin signatures of plant regulatoryelements. Nat Plants 5, 1250–1259 (2019).

**Supplementary Table 4:** Arabidopsis terminator design

| **Name** | **Gene** | **3' UTR (TAIR10)** | **Termination Window (Mo et al. 2021)** | **Longest Read-through (Mo et al. 2021)** | **Intergenic** | **Orientation** | **Notes** | **Acceptable** | **Selection** | **Final notes** |
|---|---|---|---|---|---|---|---|---|---|---|
| ACT2 | AT3G18780 | | 165 | 377 | No | Tandem | Promoter for next gene | No | | |
| ACT8 | AT1G49240 | | 123 | 281 | No | Tandem | Promoter for next gene | No | | |
| AGP15 | AT5G11740 | 460 | 196 | **492** | No | Convergent | | Yes | 542 | Tested |
| APX1 | AT1G07890 | 230 | 181 | **689** | No | Convergent | | Yes | 694 | Tested, accidentally added 5 bp instead of 50 bp |
| ARF1 | AT1G23490 | | 226 | 645 | No | Tandem | Promoter for next gene | No | | |
| AVP1 | AT1G15690 | | 155 | 389 | No | Convergent | Overlap with next gene | No | | |
| EF1A1 | AT5G60390 | **345** | 153 | 308 | No | Convergent | | Yes | 395 | Tested |
| EF1A2 | AT1G07940 | **401** | 124 | 245 | Yes | | | Yes | 451 | Tested |
| EF1A3 | AT1G07920 | | 134 | 223 | No | Tandem | Promoter for next gene | No | | |
| EF1A4 | AT1G07930 | | 117 | 251 | No | Tandem | Promoter for next gene | No | | |
| FAD2 | AT3G12120 | **341** | 120 | 313 | Yes | Convergent | | Yes | 391 | Tested |
| GADPH | AT1G13440 | **388** | 144 | **388** | No | Convergent | | Yes | 438 | Tested |
| HTR5 | AT4G40040 | | 102 | 263 | No | Tandem | Promoter for next gene | No | | |
| LEA26 | AT2G44060 | | 153 | 295 | No | Tandem | Promoter for next gene | No | | |
| PCC1 | AT3G13920 | | 131 | 351 | No | Tandem | Promoter for next gene | No | | |
| PLDA1 | AT3G15730 | | 225 | 577 | No | Convergent | Overlap with next gene | No | | |
| SHM4 | AT4G13930 | | 357 | 1092 | No | Covergent | Too long | No | | |
| TPI | AT3G55440 | | 221 | 510 | No | Convergent | Overlap with next gene | No | | |
| TUB | AT5G62690 | | 129 | 254 | No | Tandem | Promoter for next gene | No | | |
| TUB2 | AT5G62690 | | 129 | 254 | No | Tandem | Promoter for next gene | No | | |
| TUB6 | AT5G12250 | | 142 | 239 | No | Tandem | Promoter for next gene | No | | |
| UBC10 | AT5G53300 | | 126 | 315 | Yes | | Intron in 3' UTR | No | | |
| UBQ1 | AT3G52590 | | 162 | 388 | No | Convergent | Overlap with next gene | No | | |
| UBQ10 | AT4G05320 | 376 | 128 | **416** | No | Convergent | | Yes | 466 | Tested |
| UBQ11 | AT4G05050 | 267 | 170 | **507** | Yes | | | Yes | 557 | Tested |
| UBQ4 | AT5G20620 | 262 | 156 | **304** | No | Convergent | | Yes | 354 | Tested |

Selected for charatcerisation

Terminator selection criteria
1. Not in tandem orientation with downstream neighbouring gene, including lncRNA (within 2 kb)
2. No 3'UTR overlap with neigbouring gene body
3. Does not contain intron

3' UTR from TAIR10 annotation
Termination window and longest read-through from Mo, W. et al. Landscape of transcription termination in Arabidopsis revealed bysingle-molecule nascent RNA sequencing. Genome Biol 22, 322 (2021).

**Supplementary Table 5:** Oligonucleotides used in this study

| Description | Forward primer (5' to 3') | Reverse primer (5' to 3') |
|---|---|---|
| RUBY cDNA for RT-qPCR expression analsysis | CTACCGTGATCAAGAATGGATC | CAGCTTGGTGGTCTGCTG |
| UBC9 cDNA for RT-qPCR expression analsysis | TCACAATTTCCAAGGTGCTG | GTGGACTCGTACTTGTTCTTGTC |
| Arabidopdis thaliana genome for qPCR T-DNA copy number analysis | GGAACATCCTATTCTACTTACCGAG | AACAGCCTGAATAGCCACATAC |
| RUBY CDS for qPCR T-DNA copy number analysis | CACTCATTTCCCTCAGACTCG | CACTATTCGGGATCGTGC |
| Agrobacterium genome for qPCR T-DNA plasmid copy number analsysis | GAGTACCGGAATCTCGTCAAAGCC | CGAAGATCTCTACGGCAACTACCTGG |
| T-DNA plasmid backbone for qPCR T-DNA plasmid copy number analsysis | GATCATCCTGATCGACAAGACCGG | CTGCCGAGAAAGTATCCATCATGGC |
| Low-cycle PCR detection of T-DNA - bar | GAACAGGCAAAGAGAAATCG | GCTGATATCCGTAGAGCTACTG |
| Low-cycle PCR detection of T-DNA - RUBY | GTTCGGCTTCGACATCAAC | TCAAGTTTGGCCTGCATG |
| Low-cycle PCR detection of T-DNA - Backbone | CGTGAGTTTTCGTTCCACTG | CTTACCGGATACCTGTCCG |
| Low-cycle PCR detection of T-DNA - LB read-through | CCACCACTTCAAGAACTCTGTAG | TGCTCAGAACTCACGACTCC |
| iPCR genomic DNA amplification - 1.7 kb | AACCGCGTTATTTTATATGTGCC | GCCGATGTTTTCCAATCCC |
| iPCR genomic DNA amplification - 3.0 kb | GCATGGACCGTTCTCTATTACAG | TCGAATAGATCGTGTAATGGTTATTTGG |
| iPCR genomic DNA amplification - 4.8 kb | CATCTTTATGTCCTTAAAATCGTTGGG | AGGAATACTCTCCCTTGGTCA |
| Inverse PCR mapping at LB (Temporary primers) | TTCAGCACCACGAAATAGTAGC | CAGTCCCGAGCCGTCTTACTC |
| Inverse PCR mapping at RB (Temporary primers) | CCTCGTGAATAATCAGGGTGAC | CTATTGTAAAGCGAAGTGTAAGGTG |
| Inverse PCR mapping at LB (Final barcode primers) | TTCGGGAGCGGATTATACAC | CGGATCGAACTTAGGTAGCC |
| Inverse PCR mapping at RB (Final barcode primers) | ACAAGGAGTCGGCATATCAC | AGACAAGCCTTAACCGTAGG |
